# Intestinal goblet cell loss induces microbiota-independent mucosal and systemic IgA dysregulation

**DOI:** 10.64898/2026.11.16.766412

**Authors:** Sophie Vanhercke, Alexandra Thiran, Gillian Blancke, Maarten Ciers, Ioanna Petta, Maude Jans, Marie Thorp, Karel F. A. Van Damme, Helena Catherine Aegerter, Vanessa Andries, Tino Hochepied, Wendy Toussaint, Victer Verbrugge, Sofía de Penning Aguilar, Laurine Vandewiele, Inés Lammens, Stijn Vanhee, Geert van Loo, George Birchenough, Bart N. Lambrecht, Lars Vereecke

## Abstract

Goblet cells are essential for establishing the intestinal mucus layer and exert diverse immunomodulatory functions. The mucus barrier cooperates with secretory immunoglobulin A (IgA) to regulate host-microbiota interactions and maintain intestinal immune homeostasis. The extent to which goblet cells directly regulate IgA responses remains unclear. Here, we show that depletion of Muc2-expressing intestinal secretory cells in mice caused barrier dysfunction and intestinal inflammation accompanied by profound mucosal and systemic IgA dysregulation, including hyper-IgA, circulating IgA-IgG immune complexes and renal IgA deposition. Selective Paneth cell depletion did not reproduce these IgA abnormalities, and Muc2-deficient mice showed an intermediate phenotype, implicating loss of goblet-cells rather than Paneth cells or of Muc2 loss in this phenotype. Germ-free rederivation markedly attenuated intestinal inflammation but did not prevent mucosal and systemic hyper-IgA or IgA-IgG immune-complex formation, demonstrating that goblet cell loss drives IgA dysregulation independently of live microbial colonisation. Goblet cell loss and IgA deficiency shaped the mucosa-associated microbiota largely independently, as goblet cell depletion promotes expansion of the genus *Akkermansia*, whereas IgA deficiency reduced *Akkermansia* but allowed expansion of *Mucispirillum* and aggravated colonic pathology, indicating that elevated IgA response is partially protective. These findings reveal goblet cells as critical regulators of mucosal and systemic IgA homeostasis beyond mucus production and point to epithelial-immune interactions as potential contributors to IgA-mediated diseases.

## Introduction

Intestinal and systemic immune homeostasis critically depend on the integrity of the intestinal barrier, a multilayered defence system composed of epithelial junctions, mucus, antimicrobial peptides and secretory immunoglobulin A (sIgA). Goblet cells secrete the gel-forming mucin MUC2, which organizes into a double colonic mucus layer with a sterile inner-mucus layer (IML) that spatially separates the intestinal microbiota from the epithelial surface (*1–3*). In addition to producing mucus, goblet cells sense microbial products and deliver soluble luminal antigens to underlying dendritic cells through goblet cell associated passages (GAPs) (*4*). Paneth cells (PCs), located in the small intestine, secrete antimicrobial peptides that restrict microbial contact and shape microbial community composition (*5*). A third essential component of the intestinal mucosal barrier is sIgA, the most abundantly produced antibody isotype in the body. sIgA is secreted by mucosal plasma cells and transported across the epithelium by the polymeric immunoglobulin receptor (pIgR). Beyond neutralizing pathogens and toxins, sIgA promotes bacterial agglutination and enchained growth, modulates microbiota composition and facilitating mucosal colonization of selected commensal bacteria (*6–9*). Intestinal IgA responses are strongly shaped by microbial colonization: germ-free animals produce markedly less intestinal IgA, while commensal bacteria promote IgA-responses through both T-cell-dependent and T-cell-independent pathways. However, the contributions of specific intestinal epithelial cell populations and microbial signals to IgA immune regulation remain poorly understood. Disruption of the secretory barrier components is a hallmark of inflammatory bowel disease (IBD), where goblet cell depletion, mucus defects, Paneth cell dysfunction and dysregulated mucosal IgA responses contribute to chronic intestinal inflammation (*3*, *10–12*). Notably, patients with IBD display an increased risk of developing IgA nephropathy (IgAN) (*13*, *14*) and genetic studies have identified shared susceptibility loci that converge on epithelial barrier integrity, host-microbial interactions and mucosal immune regulation, supporting the existence of a pathogenic gut-kidney axis (*15*). IgA nephropathy, the most common primary glomerulonephritis, is characterized by mesangial deposition of IgA-containing immune complexes that drive progressive renal injury (*16*). Current models propose that dysregulated mucosal IgA responses give rise to aberrantly glycosylated, galactose-deficient IgA1, which is recognized by anti-glycan IgG antibodies, forming pathogenic IgA-IgG immune complexes that deposit in the kidney (*17*). Elevated intestinal permeability, altered mucosal immune responses and intestinal inflammation have been associated with IgAN (*18*, *19*) and the gut microbiota has also emerged as a key player (*20*). Several studies have reported reproducible alterations in the gut microbiota of IgAN patients, including expansion of mucin-degrading and IgA-targeting bacterial taxa such as *Akkermansia* and members of the Ruminococcaceae family (*21–23*). Experimental studies further suggest that intestinal microbes can directly influence the pathogenicity of IgA by modifying its glycosylation state and promoting immune-complex formation (*21*). However, the epithelial, immune and microbial mechanisms that initiate dysregulated IgA responses remain poorly understood and it is unknown whether primary defects in the intestinal secretory barrier are sufficient to trigger them. In this study, we address this gap by investigating the contribution of intestinal secretory cells to IgA immune dysregulation, including the formation of circulating IgA-containing immune complexes and renal IgA deposition, and by examining the role of the intestinal microbiota to these processes.

## Results

### Generation and validation of an intestinal secretory cell-targeting Muc2iCre line

To enable selective genetic targeting of intestinal Muc2-producing epithelial cells, we generated a transgenic mouse line expressing the mammalian codon-optimized Cre recombinase (iCre) (*24*) under the control of the *Muc2* promoter using bacterial artificial chromosome (BAC) transgenesis. Targeted locus amplification analysis (TLA, Cergentis) revealed integration of the transgene on chromosome 3 (Cergentis, Addendum 1). To assess the specificity of Cre activity, Muc2iCre mice were crossed with Rosa26-YFP and Rosa26-RFP reporter mice, hereafter referred to as Muc2^YFP/+^ and Muc2^RFP/+^ mice, respectively (Fig. 1A). In Muc2^YFP/+^ mice, reporter expression detected by immunostaining was observed in goblet cells throughout the small and large intestine and in Paneth cells in the small intestine (Fig. 1B). Reporter activity was not detected in the lung or cervix, although recombination was observed in a small population of distal gastric goblet cells (Fig. S1A). Muc2^RFP/+^ mice showed a similar expression pattern of reporter-positive cells in the ileum and colon (Fig. S1B).

**Figure 1:**
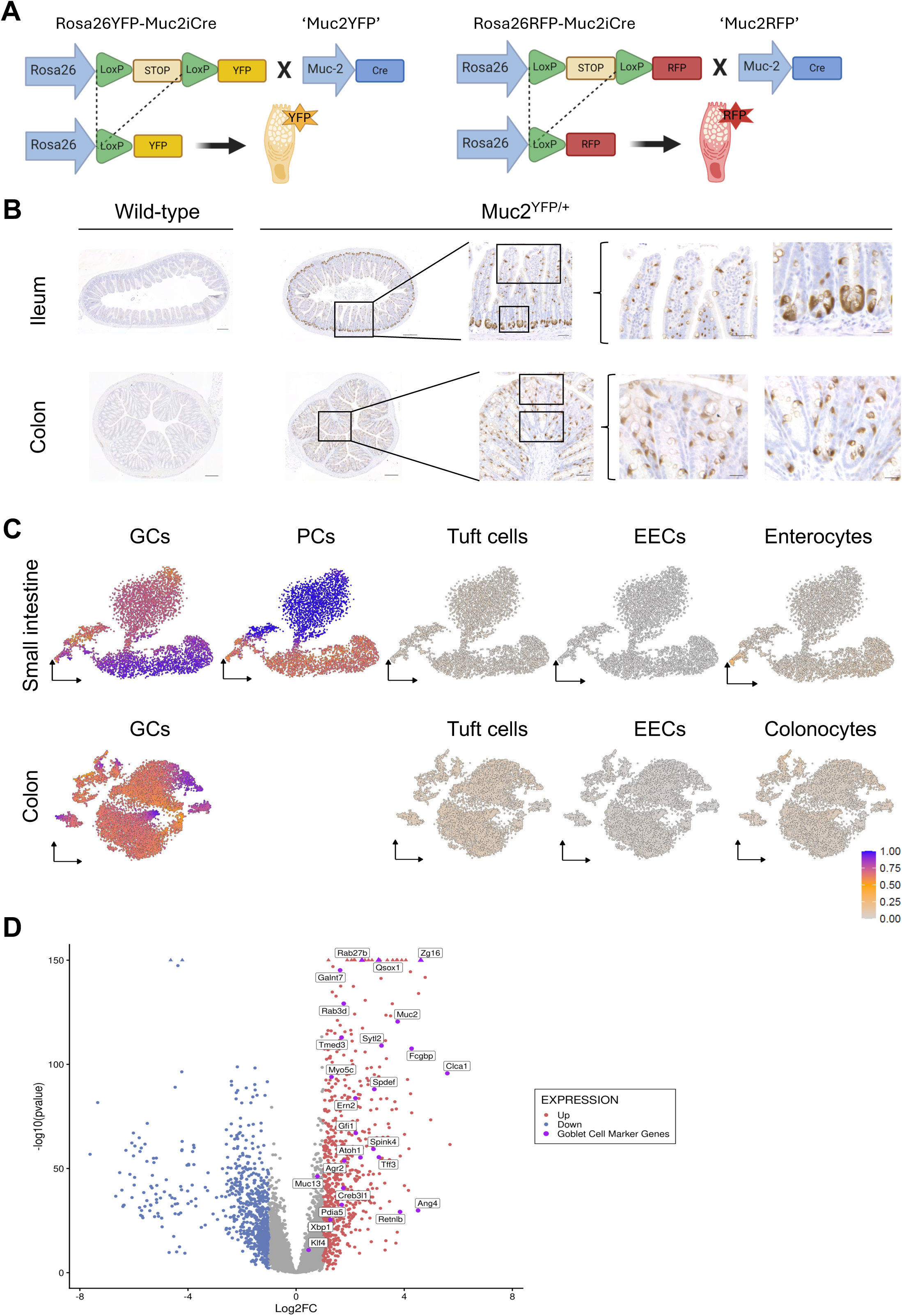
Muc2iCre drives reporter expression specifically in goblet and Paneth cells. (A) Schematic firgure of the Rosa26-YFP (left) and Rosa26-RFP (right) reporter alleles. A LoxP-flanked STOP cassette prevents reporter expression until Cre, driven by the Muc2 promoter, excises it, yielding Muc2^YFP^ and Muc2^RFP^ mice. (B) Representative images of anti-GFP immunohistochemistry (DAB, brown; hematoxylin counterstain) of ileum and colon from wild-type (WT) and Muc2^YFP/+^ mice. WT sections show no staining. Reporter-positive cells were identified as goblet cells and Paneth cells (ileal crypt base) by morphology and position. Scale bars: 200 µm (overview), 50 µm (boxed region), 20 µm (magnified views). (C) tSNE plots of scRNA-seq data from FACS-sorted RFP+ epithelial cells from small intestine (top) and colon (bottom) of Muc2^RFP/+^ mice. Color indicates the expression score of marker gene sets for goblet cells (GC), Paneth cells (PC), tuft cells, enteroendocrine cells (EEC) and enterocytes/colonocytes (n = 1 mouse; colon = 12,647 cells; small intestine = 5,536 cells). PCs are absent from the colon. (D) Volcano plot of differentially expressed genes between YFP+ and YFP-colonic epithelial cells (n = 5 mice per group). Red, up; blue, down in YFP+ (|log2FC| > 1, adj. p < 0.05); purple, goblet cell marker genes.

We next characterized the identity of our reporter-positive intestinal epithelial cells by performing single-cell RNA sequencing on sorted RFP-positive epithelial cells from the ileum and colon (VIB Nucleomics Core - 10X Genomics). Ileal RFP-positive cells expressed transcriptional signatures characteristic of goblet or Paneth cells, whereas colonic RFP-positive cells displayed only goblet cell signatures (Fig. 1C). We detected no distinct populations expressing signatures of enterocytes, colonocytes, tuft cells or enteroendocrine cells. Colonic RFP-positive cells included goblet cells expressing markers associated with both proximal (*Car8*) and distal (*B3gnt7*) colonic regions (Fig. S1C).

As an independent validation, bulk RNA sequencing of sorted colonic YFP-positive and YFP-negative epithelial cells demonstrated strong enrichment of characteristic goblet cell genes in the YFP-positive fraction, including *Muc2, Fcgbp, Zg16, Spdef, Agr2* and *Xbp1* (Fig. 1D). Furthermore, we profiled transcriptional programs of sorted YFP-positive colonic cells during *Trichuris muris* infection (day 10 post infection), a well-established infection model of type-2 immunity characterized by goblet cell hyperplasia and mucus-dependent parasite expulsion (*25*, *26*). Single-cell and bulk RNA sequencing revealed marked remodelling of goblet cell populations and transcriptional programs, including a relative expansion of intercrypt goblet cells, reduced numbers of proliferating goblet cells and induction of immune-related pathways (Fig. S1D-F and S2A-C).

These findings illustrate the utility of Muc2^YFP/+^ and Muc2^RFP/+^ reporter mice for studying goblet cell biology during homeostasis, infection and various physiological or pathological conditions.

### Constitutive depletion of intestinal secretory cells disrupts epithelial homeostasis

To investigate the physiological consequences of intestinal goblet and Paneth cell depletion, we crossed Muc2iCre mice with Rosa26-DTA mice to generate Rosa26DTA-Muc2iCre mice (hereafter referred to as Muc2^DTA^), in which expression of diphtheria toxin subunit A (DTA) selectively expressed in Muc2-expressing cells (Fig. 2A) (*27*, *28*). Constitutive DTA expression resulted in continuous ablation of goblet and Paneth cells. Both heterozygous (Muc2^DTA/+^) and homozygous (Muc2^DTA/DTA^) mice were generated.

**Figure 2:**
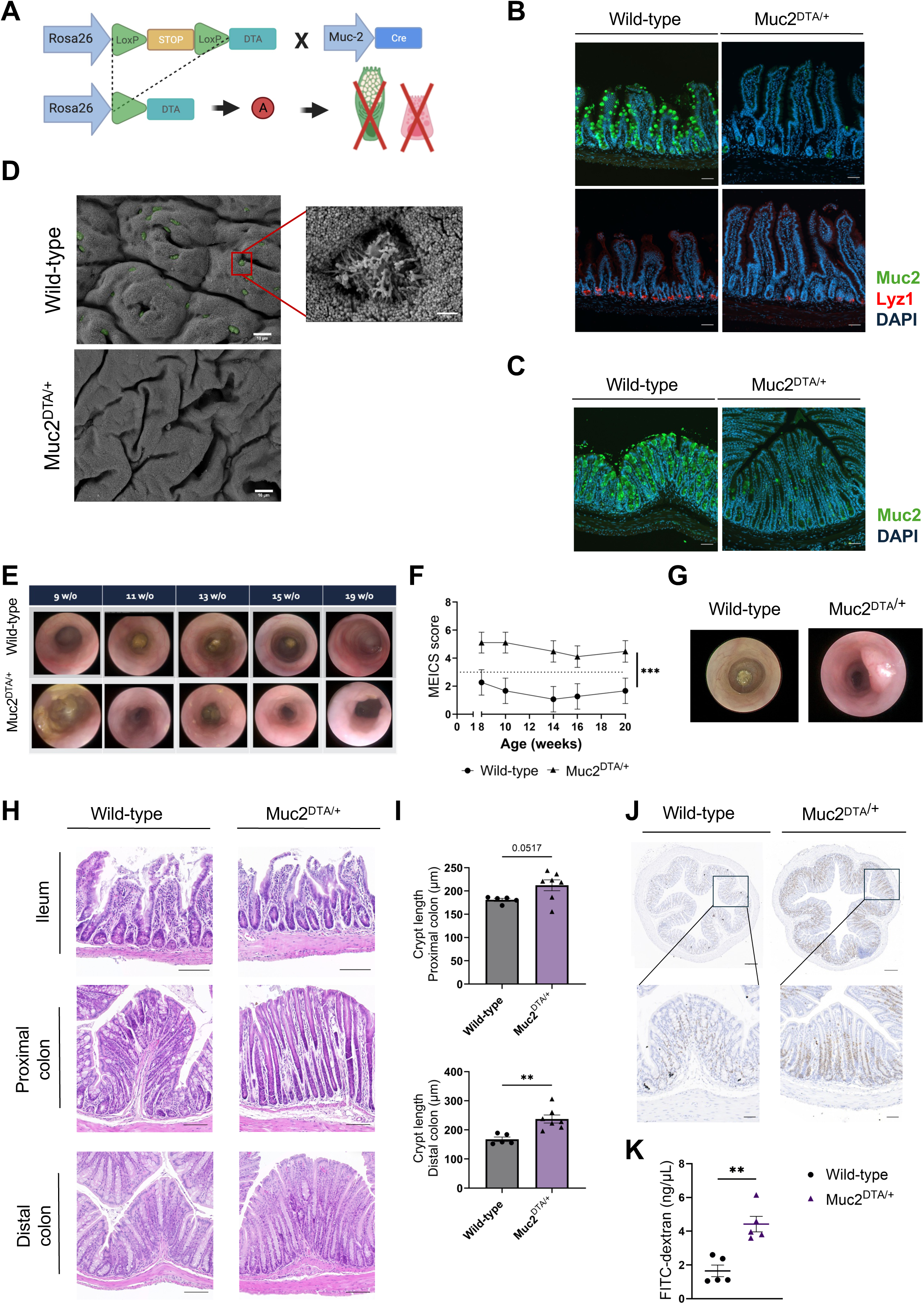
Depletion of goblet and Paneth cells in Muc2^DTA/+^ mice causes colonic pathology and barrier dysfunction. (A) Schematic figure of the Muc2^DTA^ GC depletion model. Cre-LoxP recombination drives expression of diphtheria toxin subunit A (DTA) in Muc2-expressing cells, leading to constitutive depletion of GCs and PCs. (B, C) Representative immunofluorescence images of ileal (B) and colonic (C) sections from WT and heterozygous Muc2^DTA/+^ mice stained for Muc2 (green), Lyz1 (red; Paneth cells, ileum only) and DAPI (blue). Scale bars, 50 µm. (D) Scanning electron microscopy images of the colonic surface epithelium of WT and Muc2^DTA/+^ mice. Intercrypt GCs (green) are visible in WT mice only. The boxed region is shown at higher magnification. Scale bars, 10 µm and 1 µm (magnified view). (E) Representative colonoscopy images of WT and Muc2^DTA/+^ mice showing varying degrees of intestinal pathology. (F) Murine endoscopic index of colitis severity (MEICS) scores of colonoscopy recordings over time from male WT (n = 5) and Muc2^DTA/+^ (n = 8) mice. REML model: main effect of genotype (***p < 0.001); time and time × genotype interaction, ns. (G) Colonoscopy images of a 20-week-old WT and Muc2^DTA/+^ mouse, the latter with an adenomatous polyp. (H) Representative H&E stained sections of ileum, proximal and distal colon of WT and Muc2^DTA/+^ mice. Scale bars, 100 µm. (I) Crypt length in proximal (unpaired t-test; p = 0.052) and distal colon (unpaired t-test; p = 0. 0030) of WT (n = 5) and Muc2^DTA/+^ (n = 7) mice. (J) Ki67 staining of colonic sections from WT and Muc2^DTA/+^ mice. Scale bars, 200 µm (overview) and 50 µm (magnified views). (K) Serum FITC-dextran 4 h after oral gavage in WT (n = 5) and Muc2^DTA/+^ (n = 5) mice, as a measure of intestinal permeability. Welch’s t-test; p = 0.0017.

Immunofluorescence staining confirmed efficient depletion of goblet (Muc2+) cells in ileum and colon and Paneth (Lysozyme+) cells in ileal sections from Muc2^DTA/+^ mice (Fig. 2B,C). Scanning electron microscopy (SEM) further revealed loss of intercrypt goblet cells from the colonic surface epithelium in Muc2^DTA/+^ mice (Fig. 2D), while quantitative PCR (RT-qPCR) confirmed strong reductions in the expression of the goblet cell marker *Muc2* and the Paneth-cell marker *Lyz1*, in both Muc2^DTA/+^ and Muc2^DTA/DTA^ mice (Fig. S3C).

Consistent with disruption of the protective mucus barrier, SEM analysis revealed increased bacterial adherence to the colonic epithelium of Muc2^DTA/+^ mice, including the formation of surface-associated biofilms (Fig. S3A). Homozygous Muc2^DTA/DTA^ mice, but not heterozygous Muc2^DTA/+^ mice, had a lower body weight than WT littermates (Fig. S3B).

Goblet and Paneth cell depletion in heterozygous Muc2^DTA/+^ mice was occasionally incomplete (Fig. 2B,C), resulting in residual secretory cells in a minority of crypts. In contrast, homozygous Muc2^DTA/DTA^ mice exhibited complete depletion of the secretory epithelial compartment (data not shown), but developed a markedly more severe phenotype which resulted in frequent early-life mortality. Therefore, all subsequent phenotypic analyses were performed in heterozygous Muc2^DTA/+^ mice.

Longitudinal colonoscopy demonstrated progressive development of intestinal pathology in Muc2^DTA/+^ mice, with significantly increased Murine Endoscopic Index of Colitis Severity (MEICS) scores compared with WT littermates (Fig. 2E,F) (*29*). Disease severity varied considerably between individual mice, ranging from mild pathology to the development of adenomatous polyps. Polyps were detected endoscopically in 24% of female and 36% of male Muc2^DTA/+^ mice (n = 22 males and 21 females), with the earliest lesions appearing at six weeks of age (Fig. S3D). By 20 weeks of age, large adenomatous polyps were readily detected by both endoscopy and histological analysis (Fig. 2G, Fig. S3E). Histological examination revealed pronounced epithelial hyperplasia throughout the colon, characterized by crypt elongation and occasional tubular adenomas (Fig. 2H). Immune cell infiltration was largely restricted to adenomatous lesions and was minimal in non-dysplastic mucosa. In the small intestine, only mild abnormalities were observed, including villus blunting, crypt hyperplasia and sporadic inflammatory infiltrates (Fig. S3F). Quantification of colonic crypt length confirmed a significant increase in distal colonic crypt length in Muc2^DTA/+^ mice compared to WT littermates (Fig. 2I) and Ki67 immunostaining revealed markedly expanded epithelial proliferation (Fig. 2J). Furthermore, oral FITC-dextran administration revealed significantly increased intestinal permeability in Muc2^DTA/+^ mice compared with wild-type controls, indicating impaired epithelial barrier function (Fig. 2K).

Flow cytometric analysis on lamina propria immune cells demonstrated relatively modest immune alterations despite the profound epithelial defect. Myeloid cell composition in the small intestine remained largely unchanged, whereas the colonic lamina propria exhibited increased CD45+ leukocytes, driven primarily by expansion of neutrophils together with reduced macrophage numbers (Fig. S4A). Within the T-cell compartment, ileal Th1 cells were reduced whereas colonic FoxP3+ regulatory T cells were increased (Fig. S4B). Immune alterations in lung, spleen and synovium were comparatively minor, indicating that chronic depletion of intestinal secretory epithelial cells primarily induces localized and modest intestinal pathology.

To assess whether Muc2DTA-mediated cell depletion affected extra-intestinal tissues, histological analysis was performed on 20-week-old mice. No abnormalities were detected in the lung or liver (Fig. S3G). Because the Muc2^YFP^ reporter line exhibited limited recombination in the distal stomach (Fig. S1A), gastric tissue was also examined. Despite this reporter activity, no histopathological abnormalities were observed in the stomach of Muc2^DTA/+^ mice (Fig. S3G).

Together, these findings demonstrate that constitutive depletion of goblet and Paneth cells results in chronic disruption of epithelial barrier homeostasis characterized by epithelial hyperproliferation, increased intestinal permeability, bacterial adherence and relatively mild intestinal inflammation.

To enable transient depletion of intestinal secretory cells, we additionally generated Muc2DTR mice by crossing Muc2iCre mice with Rosa26-DTR mice (Fig. S2D). Intraperitoneal administration of diphtheria toxin induced rapid and selective ablation of goblet and Paneth cells, as confirmed by Alcian Blue-Periodic Acid Schiff staining of ileal and colonic tissue 24 hours after injection (Fig. S2E). Acute loss of these secretory cells disrupted the mucus barrier, allowing bacteria to directly contact the intestinal epithelium, as demonstrated by EUB338 fluorescent in situ hybridization (Fig. S2E). Due to the rapid turnover of the intestinal epithelium, goblet and Paneth cell populations were fully restored within five days after diphtheria toxin administration, accompanied by re-establishment of the mucus barrier.

### Gut microbiota depletion mitigates intestinal pathology

We rederived Muc2^DTA^ mice under germ-free (GF) conditions by axenic embryo transfer to determine which components of the phenotype develop in complete absence of microorganisms. Effective depletion of intestinal secretory cells was maintained under GF conditions. Bulk RNA sequencing of intestinal epithelial cells demonstrated strongly reduced *Muc2* expression in colon and ileum, and reduced ileal *Defa24* expression in Muc2^DTA/+^ mice under both specific-pathogen-free (SPF) and GF conditions (Fig. S5A).

Despite persistent depletion of intestinal secretory cells, intestinal pathology was markedly attenuated in GF Muc2^DTA/+^ mice. Histological analysis of the ileum, proximal colon and distal colon showed largely preserved tissue architecture, without prominent inflammatory infiltrates or overt tissue damage in GF Muc2^DTA/+^ mice (Fig. 3A). Crypt length was larger in SPF than in GF mice in the proximal colon, but did not differ in the distal colon. Muc2^DTA/+^ mice had longer crypts than WT mice, strongly in the distal colon and modestly in the proximal colon (Fig. 3B). Colonoscopy revealed only mild macroscopic abnormalities in GF Muc2^DTA/+^ mice (Fig. 3C). MEICS scores remained significantly increased compared with GF wild-type controls but were lower than those observed in SPF Muc2^DTA/+^ mice (Fig. 3D).

**Figure 3:**
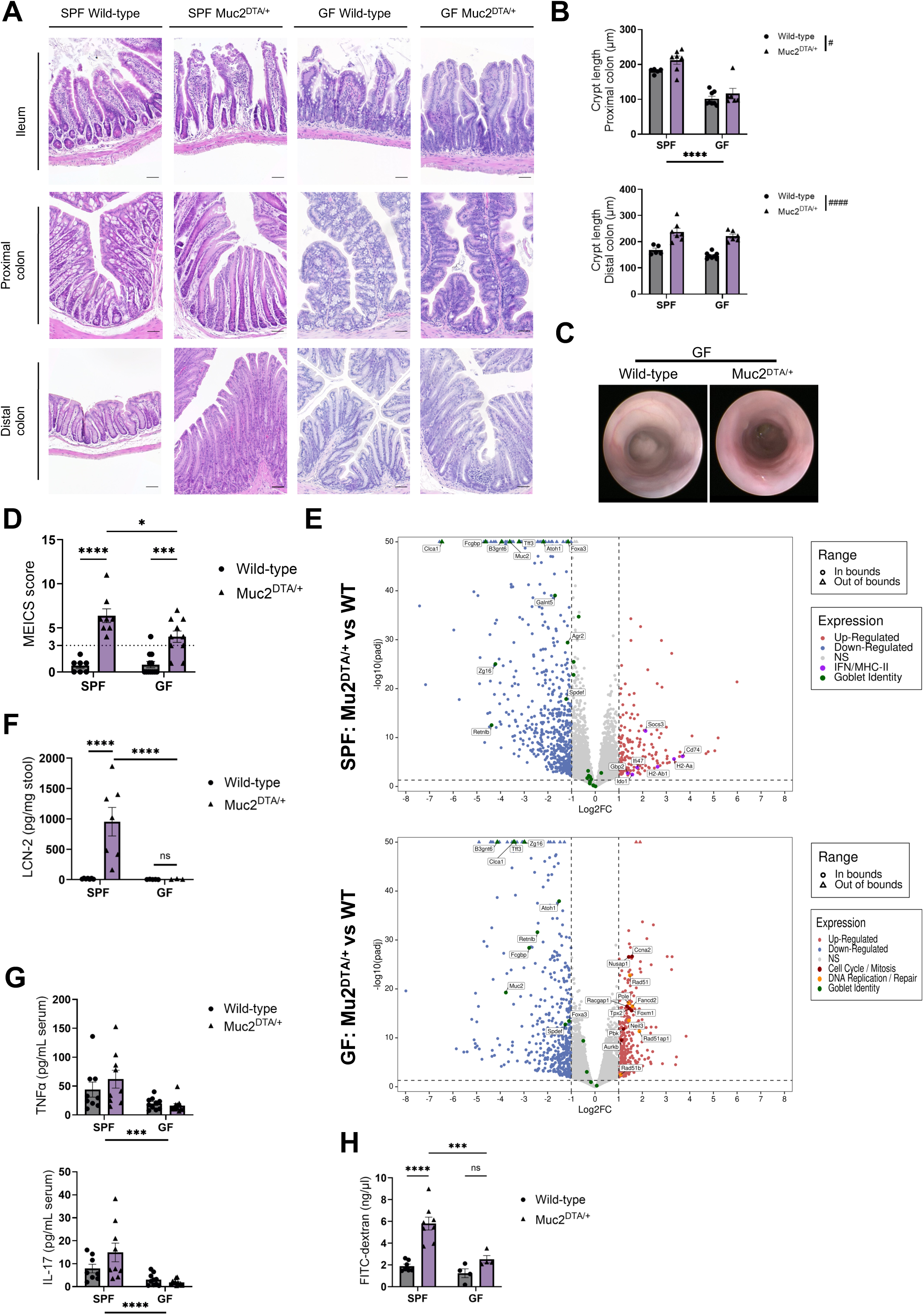
Germ-free housing of Muc2^DTA/+^ mice reduces intestinal inflammation. (A) Representative H&E-stained sections of ileum, proximal colon and distal colon from specific pathogen-free (SPF) and germ-free (GF) WT and Muc2^DTA/+^ mice. Scale bars, 50 µm. (B) Crypt length (µm) in proximal and distal colon of SPF and GF WT and Muc2^DTA/+^ mice, measured on H&E-stained sections (n = 5, 7, 8, 6). Two-way ANOVA. Proximal: main effect of housing (****p < 0.0001), main effect of genotype (#p = 0.041), interaction ns. Distal: main effect of genotype (####p < 0.0001), interaction ns. (C) Representative colonoscopy images of 20-week-old GF WT and GF Muc2^DTA/+^ mice. (D) MEICS scores from colonoscopy recordings of SPF and GF WT and Muc2^DTA/+^ mice at 20 weeks of age (n = 7, 8, 12, 10). Two-way ANOVA (genotype × housing) with Tukey’s multiple comparisons test (interaction, p = 0.034). (E) Volcano plots of RNA-sequencing data from colonic epithelial cells of Muc2^DTA/+^ versus WT mice under SPF and GF conditions, showing log2 fold change against −log10 adjusted p-value. Significantly up- and downregulated genes are shown in red and blue, respectively; non-significant genes in grey (n = 7, 8, 4, 4; thresholds: |log2FC| > 1, adj. p < 0.05). (F) Stool lipocalin-2 (LCN2), a marker of intestinal inflammation, in SPF and GF WT and Muc2^DTA/+^ mice (n = 7, 7, 5, 3). Two-way ANOVA on log10-transformed data (interaction, p = 0.0002) with Tukey’s test. (G) Bio-Plex measurement of TNFα and IL-17 in serum of SPF and GF WT and Muc2^DTA/+^ mice (n = 9, 9, 10, 11). Two-way ANOVA on log10-transformed data (TNFα) or log10(y + 1)-transformed data (IL-17). TNFα: main effect of housing (***p = 0.0001), no effect of genotype (p = 0.74), no interaction (p = 0.17). IL-17: main effect of housing (****p < 0.0001), no effect of genotype (p = 0.56), no interaction (p = 0.08). (H) Serum FITC-dextran 4 h after oral gavage in SPF and GF WT and Muc2^DTA/+^ mice (n = 7, 8, 4, 4), as a measure of intestinal permeability. Two-way ANOVA (genotype × housing) with Tukey’s multiple comparisons test (interaction, p = 0.016). Dots represent individual mice, bars show mean ± SEM and n indicates individual mice. Where four groups are shown, n is given in the order SPF WT, SPF Muc2^DTA/+^, GF WT and GF Muc2^DTA/+^. * and # denote main effects of housing and genotype, respectively, unless a post hoc comparison is stated.

To further distinguish microbiota-dependent from -independent epithelial responses, we compared the transcriptional effects of the Muc2^DTA/+^ phenotype in colonic epithelial cells under SPF and GF conditions. Analysis of genes exhibiting a significant genotype-by-microbiota interaction revealed distinct transcriptional responses. Under SPF conditions, Muc2^DTA/+^ epithelial cells showed reduced expression of genes involved in goblet cell differentiation, mucus production and mucosal barrier function, including *Atoh1, Spdef, Foxa3, Muc2, Galnt5, B3gnt6, Agr2, Fcgbp, Zg16, Clca1, Tff3 and Retnlb*. In addition, the SPF response was dominated by an inflammatory program, characterized by increased expression of MHC class II antigen-presentation genes (*Cd74, H2-Aa, H2-Ab1*) and IFN- and cytokine-responsive genes (*Socs3, Gbp2, Ifi47, Ido1*) (Fig. 3E, Fig. S5B).

In contrast, under GF conditions, the Muc2^DTA/+^ response was characterized by increased expression of genes associated with cell-cycle progression and mitosis (*Ccna2, Nusap1, Racgap1, Tpx2, Foxm1, Pbk, Aurkb*), together with genes involved in DNA replication and repair (*Rad51, Rad51b, Rad51ap1, Pole, Fancd2, Neil3*). Goblet cell-associated genes were also reduced in GF Muc2^DTA/+^ epithelium (*B3gnt6, Tff3, Zg16, Clca1, Atoh1, Retnlb, Fcgbp, Muc2, Spdef, Foxa3*) (Fig. 3E, Fig. S5B). Together, these data show a microbiota-dependent difference in the epithelial transcriptional response to goblet cell depletion. Whereas Muc2^DTA/+^ epithelial cells under SPF conditions acquired a pronounced inflammatory phenotype characterized by IFN-responsive and MHC class II antigen-presentation programs, this response was largely absent under GF conditions. These findings indicate that the microbiota is a major driver of epithelial inflammatory activation following goblet cell depletion.

Moreover, fecal LCN2, which is strongly increased in SPF Muc2^DTA/+^ mice, was not significantly elevated under GF conditions (Fig. 3F). Reduced intestinal inflammation under GF conditions was accompanied by lower systemic inflammatory cytokines. Serum TNF and IL-17 trended higher in SPF Muc2^DTA/+^ mice compared with WT, but not under GF conditions (Fig. 3G). No significant differences were observed in serum MCP-1 and IFN-γ between WT and Muc2^DTA/+^ mice in SPF and GF conditions (Fig. S5C). Intestinal barrier dysfunction in Muc2^DTA/+^ mice, assessed by FITC-dextran assay, was significantly attenuated under GF conditions, despite an increasing trend (Fig. 3H).

Consistent with findings in germ-free mice, broad-spectrum antibiotic treatment of 11-13 week old Muc2^DTA/+^ mice for six weeks ameliorated, but did not fully rescue, intestinal pathology in Muc2^DTA/+^ mice, as evidenced by lower MEICS scores and decreased crypt length compared with untreated Muc2^DTA/+^ littermates (Fig. S5D,E).

Together, our data demonstrate that the microbiota strongly amplifies intestinal inflammation and barrier dysfunction following depletion of Muc2-expressing secretory cells, while the persistence of some epithelial abnormalities under GF conditions reveals also a microbiota-independent component of the phenotype.

### Goblet cell loss promotes IgA dysregulation independent of the intestinal microbiota

Goblet cells and secretory IgA constitute complementary components of the intestinal mucosal barrier, together limiting interactions between luminal contents and the underlying mucosa. Given the disruption of intestinal barrier function and immune homeostasis following depletion of Muc2-expressing secretory cells, we next investigated whether this was accompanied by alterations in IgA responses and whether these depended on the intestinal microbiota. Strikingly, Muc2^DTA/+^ mice displayed a pronounced hyper-IgA phenotype under both SPF and GF conditions, characterized by increased fecal IgA and markedly elevated circulating total and polymeric IgA (Fig. 4A-C). This response was selective for IgA, as serum IgG and IgM levels were not significantly affected by genotype, although IgM levels differed according to microbial colonization status (Fig. S6A,B). Importantly, the persistence of hyper-IgA in GF Muc2^DTA/+^ mice contrasted with the marked attenuation of intestinal inflammation and the absence of the SPF-associated epithelial inflammatory transcriptional program under GF conditions (Fig. 3, Fig. S5B). Together, these findings demonstrate that depletion of Muc2-expressing secretory cells induces a selective mucosal and systemic hyper-IgA phenotype that does not require the intestinal microbiota and is largely uncoupled from microbiota-driven intestinal inflammation.

**Figure 4:**
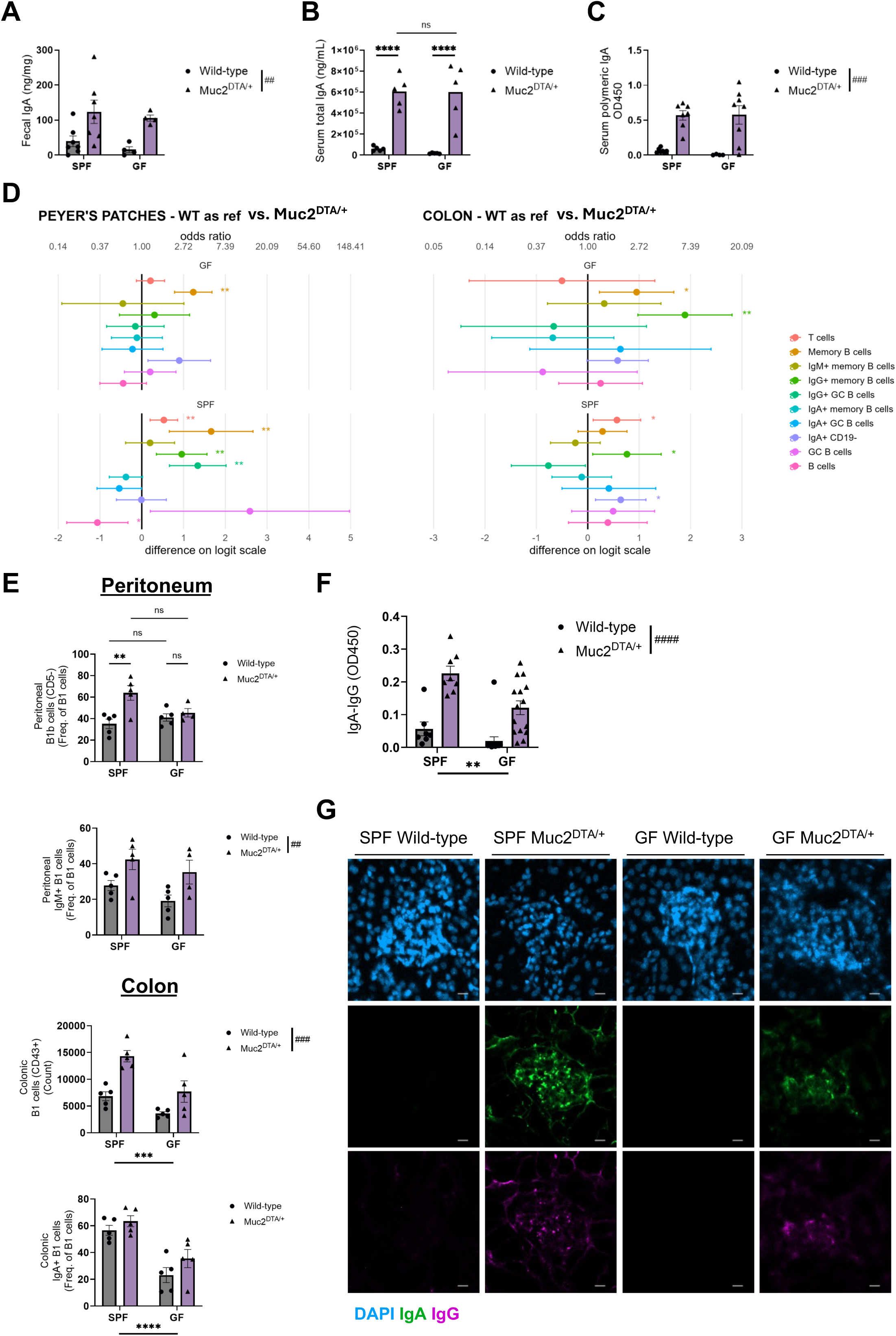
Loss of the mucus barrier dysregulates the IgA response independently of the microbiota. (A) Fecal IgA (ng/mg) in SPF and GF WT and Muc2^DTA/+^ mice (n = 7, 7, 4, 4). Two-way ANOVA: main effect of genotype (##p = 0.003), interaction ns. (B) Serum total IgA (ng/mL) in SPF and GF WT and Muc2^DTA/+^ mice (n = 5, 5, 5, 5). Two-way ANOVA (genotype × housing) with Tukey’s multiple comparisons test on log10-transformed data (interaction, p = 0.006): ****p < 0.0001; ns, not significant. (C) Serum polymeric IgA (J chain-containing IgA, OD 450) in SPF and GF WT and Muc2^DTA/+^ mice (n = 7, 7, 4, 8). GLM: main effect of genotype (###p < 0.001); housing, ns; interaction, ns. (D) Forest plots of B cell subset frequencies in Peyer’s patches and colon, showing Muc2^DTA/+^ mice relative to WT (reference) under GF (top) and SPF (bottom) conditions (n = 7, 8, 4, 4). Points show the difference on the logit scale (bottom axis; odds ratio on the top axis) with 95% CI error bars. The vertical line marks no difference (odds ratio = 1). HGLM (R). *p < 0.05, **p < 0.01. (E) Flow cytometric analysis of B1 cells in the peritoneum and colon of SPF and GF WT and Muc2^DTA/+^ mice (n = 5, 5, 5, 4 and n = 5, 5, 5, 5). Peritoneum: B1b cells (% of B1 cells), two-way ANOVA with interaction (p = 0.026); IgM+ B1 cells (% of B1 cells), main effect of genotype (##p = 0.0053). Colon: B1 cell count, main effects of housing (***p = 0.001) and genotype (###p = 0.0002), interaction ns; IgA+ B1 cells (% of B1 cells), main effect of housing (****p < 0.0001), interaction ns. (F) Serum circulating IgA-IgG complexes (OD 450) in SPF and GF WT and Muc2^DTA/+^ mice (n = 7, 8, 15, 15). Two-way ANOVA: main effects of housing (**p = 0.0013) and genotype (####p < 0.0001), interaction ns. (G) IgA and IgG staining of kidney sections from SPF and GF WT and Muc2^DTA/+^ mice. Scale bar, 5 µm. Dots represent individual mice, bars show mean ± SEM and n indicates individual mice. Where four groups are shown, n is given in the order SPF WT, SPF Muc2^DTA/+^, GF WT and GF Muc2^DTA/+^. * and # denote main effects of housing and genotype, respectively, unless a post hoc comparison is stated.

To determine how disruption of the intestinal mucus barrier affects mucosal B cell populations, we profiled B cell subsets by flow cytometry in the ileum, mesenteric lymph nodes (MLN), Peyer’s patches (PP) and colon of Muc2^DTA/+^ and WT mice housed under SPF or GF conditions. Peyer’s Patches of SPF Muc2^DTA/+^ mice (Fig. 4D), had higher proportions of T cells, memory B cells, IgG+ memory B cells and IgG+ germinal center (GC) B cells and a lower B cell fraction, than WT. In GF mice, only memory B cells were increased in PP and the IgG+ and GC changes were not detected, suggesting that the class-switched, GC-associated response in PP depends on the microbiota. In the colon (Fig. 4D), IgG+ memory B cells were increased under both GF and SPF conditions and memory B cells were also increased in GF mice, indicating that the colonic IgG+ memory B cell expansion does not require the microbiota. In contrast, higher T cell and IgA-producing plasma cell fractions (IgA+ CD19-) were detected only in SPF Muc2^DTA/+^ colon, consistent with a microbiota-dependent, T cell-associated IgA response.

Changes in the ileum and MLN (Fig. S6C) were limited compared with the PP and colon. In SPF mice, the only ileal change was a reduced IgA+ CD19-fraction in Muc2^DTA/+^ mice, opposite to the increase in the colon, whereas the MLN showed a reduced T cell fraction, opposite to the T cell increase in PP and colon. In GF mice, the ileum of Muc2^DTA/+^ showed higher T cell and IgA+ GC B cell fractions and a lower IgG+ GC B cell fraction than WT.

Peritoneal B1 cells showed partially microbiota-dependent changes in Muc2^DTA/+^ mice. In SPF mice only, B1a cells were lower and B1b cells higher as a proportion of B1 cells than in WT (Fig. 4E, Fig. S6D). In contrast, IgM+ B1 cells were higher in Muc2^DTA/+^ mice irrespective of housing, whereas IgA+ B1 cells did not differ between genotypes (Fig. 4E, Fig. S6D). In the colon, Muc2^DTA/+^ mice had more B1 cells than WT mice under both SPF and GF conditions, and B1 cell numbers were generally higher in SPF than in GF mice (Fig. 4E). This coincided with a higher frequency of B cells and higher B1a and B1b numbers (Fig. S6D), indicating an expansion of the colonic B cell compartment, including B1 cells. The proportion of IgA+ cells among B1 cells did not differ between genotypes but was much higher in SPF than in GF mice, indicating that IgA expression on colonic B1 cells depends on the microbiota (Fig. 4E). In the spleen, the frequencies of B cells, B1 cells, B1a and B1b cells and IgA+ plasma cells did not differ between WT and Muc2^DTA/+^ mice under either housing condition, indicating that the changes seen in the colon and peritoneum were not observed systemically (Fig. S6D). B1 cell frequency and B1a/B1b composition, as well as spleen size, were higher in SPF than in GF mice irrespective of genotype. We therefore examined expression of the B cell-activating cytokines *Tnfsf13b* (BAFF) and *Tnfsf13* (APRIL) in our RNA-seq dataset of intestinal epithelial cells. Expression of both genes was higher in SPF than in GF mice in ileum and colon, with little effect of genotype, apart from lower colonic *Tnfsf13* in SPF Muc2^DTA/+^ mice (Fig. S6E).

Because altered epithelial IgA transport could contribute to increased circulating IgA, we assessed expression of *pIgR*, which mediates epithelial transcytosis of dimeric IgA (*30*). In the ileum, *pIgR* expression was significantly lower in GF than in SPF Muc2^DTA/+^ mice, whereas no significant difference between WT and Muc2^DTA/+^ mice was detected under either housing condition. In the colon, *pIgR* expression was reduced in Muc2^DTA/+^ mice compared with WT, irrespective of housing, and was also lower under GF than SPF conditions (Fig. S6F). Despite these transcriptional changes, pIgR immunostaining showed a preserved colonic epithelial expression pattern in Muc2^DTA/+^ mice (Fig. S6G). Together, we show that *pIgR* expression is influenced by both genotype and colonization status. However, elevated fecal IgA in Muc2^DTA/+^ mice under both SPF and GF conditions argues against a major defect in luminal IgA transport.

Given the pronounced increase in circulating IgA in Muc2^DTA/+^ mice, we next investigated whether this was accompanied by the formation of IgA-containing immune complexes, which contribute to the pathogenesis of IgA-associated diseases such as IgA nephropathy (IgAN). Therefore, we quantified circulating IgA-IgG immune complexes in SPF and GF mice. Circulating IgA-IgG immune complexes were significantly elevated in Muc2^DTA/+^ mice under both housing conditions, although the increase was markedly attenuated in GF Muc2^DTA/+^ mice (Fig. 4F). Thus, microbial colonization is not required for the elevated mucosal and systemic IgA responses, but it amplifies IgA-IgG immune complex formation under conditions of intestinal secretory cell loss.

Elevated serum IgA and circulating IgA-containing immune complexes are linked to IgA nephropathy (*16*). We therefore examined kidney sections for IgA and IgG deposition by immunofluorescence. Glomerular IgA and IgG staining was low in wild-type mice but increased in Muc2^DTA/+^ mice, more strongly under SPF than GF conditions (Fig. 4G, Fig. S7A).

Despite the presence of IgA deposits in Muc2^DTA/+^ kidneys, histological examination of kidney sections revealed no overt pathological abnormalities (Fig. S7B). Consistent with this, RT-qPCR of kidney tissue showed no significant differences between Muc2^DTA/+^ and WT mice in the pro-inflammatory markers *Il1b* and *Tnf*. The tubular injury marker *Lcn2* trended higher in Muc2^DTA/+^ kidneys, whereas *Nphs1* (nephrin), a podocyte slit diaphragm protein, trended lower, which may point to early podocyte stress or damage (Fig. S7C). Together, these findings demonstrate that loss of Muc2-expressing secretory cells induces pronounced mucosal and systemic IgA dysregulation independently of the intestinal microbiota, accompanied by distinct microbiota-dependent and -independent changes in mucosal B cell populations. Importantly, microbial colonization amplifies the formation of circulating IgA-IgG immune complexes and glomerular immunoglobulin deposition, linking intestinal secretory cell dysfunction to systemic IgA-associated pathology.

### IgA partially protects against intestinal inflammation and shapes the mucosal microbiota following goblet cell loss

Given the pronounced increase in IgA in Muc2^DTA/+^ mice, we investigated whether this response provides compensatory mucosal protection following goblet cell depletion. Muc2^DTA/+^ mice were therefore crossed onto an IgA-deficient background. IgA deficiency did not result in overtly aggravated intestinal pathology in Muc2^DTA/+^ mice (Fig. 5A), but was associated with increased distal colonic crypt length, elevated MEICS scores and elevated fecal lipocalin-2 levels compared with IgA-competent Muc2^DTA/+^ mice (Fig. 5B-D). Serum IgM levels were increased upon IgA deficiency irrespective of genotype (Fig. 5E). Together, these findings suggest that the enhanced IgA response in Muc2^DTA/+^ mice provides partial protection against intestinal inflammation following mucus loss.

**Figure 5:**
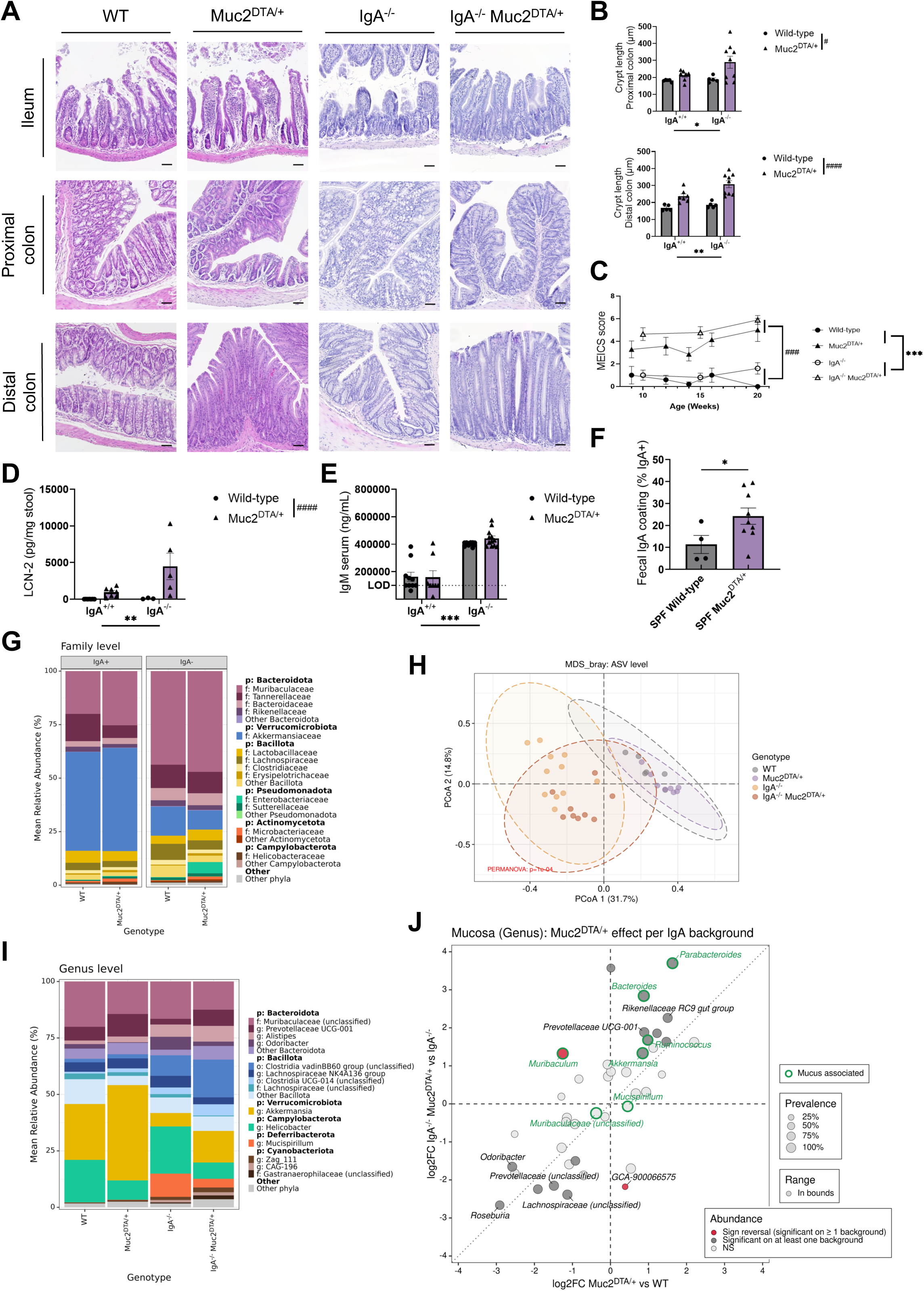
IgA deficiency exacerbates colonic pathology and shifts the mucosal microbiota in Muc2^DTA/+^ mice. (A) Representative H&E-stained sections of ileum, proximal colon and distal colon from WT, Muc2^DTA/+^, IgA^-/-^ and IgA^-/-^ Muc2^DTA/+^ mice. Scale bars, 50 µm. (B) Crypt length (µm) in proximal (top) and distal (bottom) colon of WT, Muc2^DTA/+^, IgA^-/-^ and IgA^-/-^ Muc2^DTA/+^ mice (n = 5, 7, 5, 9). Proximal: GLM main effects of genotype (#p = 0.011) and IgA availability (*p = 0.047); interaction, ns. Distal: two-way ANOVA on log10-transformed data: main effect of IgA availability (**p = 0.01) and genotype (####p < 0.0001), interaction, ns. (C) MEICS scores from colonoscopy recordings of WT, Muc2^DTA/+^, IgA^-/-^ and IgA^-/-^ Muc2^DTA/+^ mice (n = 5, 7, 5, 11). REML model: main effects of genotype (###p < 0.001) and IgA availability (***p = 0.001); interaction, ns. (D) Stool LCN2 (pg/mg stool) in WT, Muc2^DTA/+^, IgA^-/-^ and IgA^-/-^ Muc2^DTA/+^ mice (n = 7, 7, 3, 5). Two-way ANOVA on log10(y + 1)-transformed data: main effect of IgA availability (**p = 0.002) and genotype (####p < 0.0001), interaction ns (p = 0.70). (E) Serum IgM (ng/mL) in WT, Muc2^DTA/+^, IgA^-/-^ and IgA^-/-^ Muc2^DTA/+^ mice (n = 10, 8, 12, 12). GLM: main effect of IgA availability (***p < 0.001); genotype and interaction, ns. LOD, limit of detection (dotted line). (F) Percentage of IgA-coated bacteria in stool of SPF WT (n = 4) and SPF Muc2^DTA/+^ (n = 9) mice. Welch’s t-test, *p = 0.048. (G) Family-level relative abundance (%) of bacteria in the IgA-positive and IgA-negative stool fractions of SPF WT (n = 7) and Muc2^DTA/+^ (n = 8) mice, determined by metagenomic sequencing. (H) Mucosal microbiota diversity by 16S rRNA sequencing (ASV level; n = 6, 8, 11, 9). PCoA of Bray-Curtis dissimilarity. PERMANOVA, p = 0.0001. (I) Mucosal microbiota composition at genus level, as mean relative abundance (%) per genotype (n = 6, 8, 11, 9). Genera are grouped by phylum. (J) Effect of Muc2^DTA/+^ on mucosal genera by IgA background: log2 fold change of Muc2^DTA/+^ versus WT (x-axis) against IgA^-/-^ Muc2^DTA/+^ versus IgA^-/-^ (y-axis). Red points mark genera with a sign reversal (significant on at least one background, with opposite direction of effect between backgrounds), dark grey points genera significant on at least one background and light grey points genera with no significant effect (NS). Green outlines mark mucus-associated genera. Point size indicates prevalence. The dotted diagonal marks equal effects on both backgrounds. Dots represent individual mice, bars show mean ± SEM and n indicates individual mice. In B-E, n is given in the order WT, Muc2^DTA/+^, IgA^-/-^ and IgA^-/-^ Muc2^DTA/+^. * and # denote main effects of IgA availability and genotype, respectively, unless a post hoc comparison is stated.

To investigate how IgA interacts with the intestinal microbiota following goblet cell loss, we first quantified IgA coating of fecal bacteria. Muc2^DTA/+^ mice exhibited a significantly higher proportion of IgA-coated bacteria compared with wild-type littermate controls (Fig. 5F), consistent with their elevated fecal IgA levels. We next performed metagenomic profiling of IgA-coated and uncoated bacterial fractions, revealing a pronounced segregation between both fractions, with *Akkermansiaceae* particularly enriched among IgA-coated bacteria. However, the relative taxonomic composition of either fraction did not differ significantly between Muc2^DTA/+^ and wild-type mice (Fig. 5G). Thus, goblet cell depletion increases the overall proportion of IgA-coated bacteria without markedly altering the relative taxonomic distribution of IgA targeting. Nevertheless, these relative abundance profiles do not exclude increased absolute IgA coating of individual taxa in Muc2^DTA/+^ mice. We next investigated how goblet cell depletion and IgA deficiency individually and jointly influence the mucosa-associated microbiota by profiling WT, Muc2^DTA/+^, IgA^-/-^ and IgA^-/-^ Muc2^DTA/+^ mice. Microbial community composition differed between genotypes (Bray-Curtis, ASV level), separating primarily by IgA status, whereas WT and Muc2^DTA/+^ communities overlapped strongly (Fig. 5H). Muc2^DTA/+^ mice had the lowest alpha diversity, with Shannon diversity lower than in IgA^-/-^ and IgA^-/-^ Muc2^DTA/+^ mice and a trend towards lower values than WT. Observed richness was similar to WT but lower than in both IgA-deficient groups (Fig. S7D). At the genus level, Muc2^DTA/+^ mice were dominated by *Akkermansia* (42.2% vs 24.7% in WT; Fig. 5I), consistent with a less even community when goblet cells are lost. IgA deficiency lowered *Akkermansia* on both backgrounds (5.9% in IgA^-/-^, 14.0% in IgA^-/-^ Muc2^DTA/+^) and led to the appearance of *Mucispirillum*, nearly absent in WT and Muc2^DTA/+^ mice (10.3% in IgA^-/-^, 3.9% in IgA^-/-^ Muc2^DTA/+^), together with a Bacillota expansion that was most pronounced in IgA^-/-^ Muc2^DTA/+^ mice. Differential abundance analysis confirmed enrichment of *Mucispirillum* and CAG-196 and depletion of *Akkermansia* in IgA^-/-^ versus WT mice (Fig. S7E). The expansion of *Mucispirillum* in IgA-deficient mice suggests that IgA normally restrains it, whereas the lower abundance of *Akkermansia*, which was strongly IgA-coated in our IgA-seq, suggests that IgA may support its persistence in the mucosa.

Goblet cell depletion increased several mucus associated bacteria (green), including *Akkermansia*, *Bacteroides*, *Parabacteroides* and *Ruminococcus*, and decreased *Roseburia* and *Odoribacter*, with the same direction of change irrespective of IgA status (Fig. 5J). A few genera responded differently depending on IgA: *Parabacteroides* increased more strongly and *Desulfovibrio* decreased less without IgA, whereas *Muribaculum* decreased with IgA but increased without it (Fig. 5J). Together, these findings indicate that that goblet cells and IgA act as largely separate barriers shaping the mucosal microbiota, with limited interaction for some taxa.

Finally, we examined whether goblet cell depletion similarly affected the luminal microbiota by profiling fecal samples from WT and Muc2^DTA/+^ mice. In stool, goblet cell depletion did not significantly affect Shannon diversity, although observed richness tended to be lower in Muc2^DTA/+^ mice (Fig. S7F). Community composition nevertheless differed between genotypes (Bray-Curtis, Fig. S7F). At the genus level, the stool microbiota of both genotypes was dominated by Bacteroidota and Bacillota, with *Akkermansia* present at moderate and similar levels in WT and Muc2^DTA/+^ mice (Fig. S7G). Differential abundance analysis identified a limited number of genera: *Limosilactobacillus* and *Faecalibaculum* were lower in Muc2^DTA/+^ mice, whereas *Alistipes*, *Massiliomicrobiota*, unclassified Christensenellaceae and unclassified RF39 were higher (Fig. S7H). Because the luminal community is less dependent on the mucus layer than the mucosa-associated community, these modest shifts suggest that loss of goblet cells and mucus has a limited, indirect effect on the stool microbiota, possibly through altered availability of host-derived substrates.

### Goblet, not Paneth, cell loss drives systemic IgA dysregulation beyond the effects of Muc2 deficiency

To distinguish the respective contributions of goblet cell depletion, Muc2 deficiency and Paneth cell loss to the Muc2^DTA/+^ phenotype, we compared Muc2^DTA/+^ mice with Muc2-deficient mice (Muc2^-/-^), which retain goblet cells but lack Muc2 production, and with Defa24^DTA/+^ mice (*31*), in which Paneth cells are selectively depleted. Muc2-deficient mice developed more pronounced intestinal pathology than Muc2^DTA/+^ mice, with significantly increased colonic crypt length, particularly in the distal colon, and elevated MEICS scores at the age of 10 weeks old (Fig. S8A–C). Despite this more severe intestinal phenotype, serum IgA levels and circulating IgA-IgG complexes were highest in Muc2^DTA/+^ mice and comparatively lower in Muc2-deficient mice (Fig. S8E,F). In contrast, Defa24^DTA/+^ mice showed IgA and IgA-IgG complex levels comparable to wild-type mice, indicating that Paneth-cell depletion alone is insufficient to induce this hyper IgA response (Fig. S8D–F). Consistent with these findings, immunofluorescence analysis revealed glomerular IgA deposition in Muc2-/- mice, whereas no detectable deposition was observed in Defa24^DTA/+^ mice (Fig. S8G). IgG staining was weak in the glomeruli of both Muc2^-/-^ and Defa24^DTA/+^ mice.

Together, these findings identify goblet cell depletion, rather than concomitant Paneth cell loss, as the principal contributor to the pronounced systemic IgA phenotype in Muc2^DTA/+^ mice. Importantly, the stronger IgA response in Muc2^DTA/+^ mice despite more severe intestinal pathology in Muc2-/- mice indicates that systemic IgA dysregulation cannot be explained solely by Muc2 deficiency, mucus barrier disruption or the severity of intestinal inflammation. Instead, these results suggest that goblet cell loss triggers additional mechanisms of IgA dysregulation beyond the absence of the Muc2 mucus layer.

## Discussion

Our findings indicate that loss of intestinal secretory cells leads to loss of intestinal epithelial and immune homeostasis. Chronic GC depletion caused epithelial barrier dysfunction, altered mucosal microbial community structure, mucosal and systemic hyper-IgA responses, circulating IgA-IgG immune complexes and renal IgA deposition. Selective Paneth-cell depletion did not reproduce the IgA phenotype, whereas *Muc2*-deficient mice showed an intermediate phenotype. Together, our data identify loss of the goblet cells, rather than Paneth cell depletion, as the principal trigger of the disturbed IgA response. Importantly, germ-free rederivation largely separated this hyper-IgA response from the intestinal inflammatory phenotype. Stool LCN2 levels, systemic inflammatory mediators and barrier dysfunction are largely normalized under germ-free conditions, while fecal IgA, total serum IgA and polymeric IgA remained markedly elevated. Circulating IgA-IgG immune complexes were reduced but remained strongly increased and renal IgA deposits remained detectable in germ-free Muc2^DTA/+^ mice. Together, our data indicate that exaggerated IgA production is associated with goblet cell loss and is not simply a secondary consequence of dysbiosis or bacteria-epithelial contact driven mucosal stimulation.

Importantly, germ-free does not mean antigen-free and autoclaved grain-based chow contains intact dietary protein and microbial ligands, including LPS. Dietary LPS was recently shown to drive intestinal germinal center reactions and IgA diversification in germ-free mice through TLR4 (*32*), while germ-free mice on an antigen-free diet still generate polyreactive IgA plasma cells (*33*). Our data indicate that IgA dysregulation in Muc2^DTA/+^ mice does not require live microbial colonization, not that it is independent of all exogenous stimuli.

A precedent for microbiota-independent epithelial control of IgA was established by Grootjans et al., 2019 who showed that epithelial ER stress in *Xbp1*ΔIEC mice induces a T cell-independent, polyreactive IgA response through expansion and activation of peritoneal B1b cells (*34*). Notably, this B1b response persists in *Xbp1*ΔIEC mice under germ-free conditions. Our findings, however, point to a distinct pathway. Muc2^DTA/+^ mice showed a lower peritoneal B1a fraction and a higher B1b fraction under SPF conditions, but neither change was detected under GF conditions. In contrast, colonic B1a and B1b cell numbers remained higher in GF Muc2^DTA/+^ mice, although the increase was smaller than under SPF conditions. *Xbp1*ΔIEC mice also show reduced goblet cell numbers and Paneth cell dysfunction (*35*), suggesting that disruption of the secretory lineage may be a common feature of both models. In line with the B1 cell involvement reported for *Xbp1*ΔIEC mice, Muc2^DTA/+^ mice also showed B1 cell alterations, but whether B1 cells are the source of the elevated IgA and whether this IgA is T cell-independent and polyreactive, remains to be determined.

The increase in colonic IgA+ plasma cells observed under SPF conditions was also lost germ-free. Broader lymphocyte profiling reinforces tissue -or compartment-specific responses. Under SPF conditions, colonic changes included increased IgG+ memory B cells and IgA+CD19− cells, whereas Peyer’s patches showed a broader expansion of memory B-cell populations, including IgG+ and IgA+ memory cells. Several of these changes disappeared germ-free, although increased colonic memory and IgG+ memory B cells and increased Peyer’s patch memory B cells persisted. Together, we show that B cell responses vary significantly under conditions of intestinal secretory cell loss and vary according to both anatomical site and microbial status, but excessive IgA production is maintained both under colonized and germ-free conditions.

Other epithelial perturbations have linked epithelial dysfunction to systemic IgA abnormalities. Intestinal epithelial-specific loss of the epithelial polarity factor AP-1B (*Ap1m2*ΔIEC) was shown to disrupt pIgR expression and trafficking and produces elevated abnormal IgA, IgA-IgG complexes and glomerular deposition (*36*). In *Ap1m2*ΔIEC mice however, antibiotic treatment reduces abnormal IgA and renal deposition. pIgR deficiency itself causes accumulation of polymeric IgA in serum but markedly reduces luminal IgA, illustrating how impaired epithelial transport can redistribute IgA between systemic and mucosal compartments. *pIgr* expression is reduced in parts of the Muc2^DTA/+^ mice intestine, indicating that GCs significantly contribute to IgA transcytosis, and that altered IgA transport in Muc2^DTA/+^ mice may therefore contribute to systemic IgA accumulation. However, impaired pIgR transport alone cannot explain the phenotype because fecal IgA is increased, rather than reduced, and remains increased in germ-free Muc2^DTA/+^ mice. The combined increase in luminal and circulating polymeric IgA instead argues for enhanced IgA production in conditions of goblet cell loss.

Bacterial sensing by epithelial cells can promote T cell-independent IgA responses through the expression of APRIL and BAFF (*37*). Here, however, the preservation of hyper-IgA in germ-free Muc2^DTA/+^ mice points to an IgA-promoting mechanism that does not require the microbiota. Epithelial *Tnfsf13* (APRIL) and *Tnfsf13b* (BAFF) expression was higher in SPF than in GF mice but was largely unaffected by genotype, arguing against a major contribution of epithelial APRIL or BAFF expression to the elevated IgA. One possibility is that chronic goblet cell loss creates an epithelial stress that promotes IgA independently of microbial ligands. Whether chemokines, altered antigen access, specific epithelial stress mediators, or other pathways provide this microbiota-independent signal remains unclear.

MIg-seq further argues against a major shift in the microbes targeted by IgA following goblet cell depletion. IgA-coated and uncoated fractions showed distinct taxonomic profiles, with *Akkermansiaceae* particularly enriched among coated bacteria, but their relative composition did not differ significantly between wild-type and Muc2^DTA/+^ mice. Nevertheless, Muc2^DTA/+^ mice exhibited a higher proportion of IgA-coated bacteria, indicating increased overall IgA coating without major changes in taxonomic targeting. Interestingly, comparative profiling of WT, Muc2^DTA/+^, IgA^-/-^ and IgA^-/-^ Muc2^DTA/+^ mice revealed that IgA deficiency had a greater impact on mucosa-associated microbial communities than goblet cell depletion, with particularly pronounced changes in *Akkermansia* and *Mucispirillum*. The strong IgA coating of *Akkermansia*, together with its reduced abundance in IgA-deficient mice, suggests that IgA may promote its persistence rather than simply mediate bacterial exclusion. Conversely, the expansion of *Mucispirillum* in IgA-deficient mice suggests a potential role for IgA in restricting its abundance. Together, these findings highlight distinct but complementary roles of goblet cells and IgA in shaping mucosal microbial communities, while the specificity and functional consequences of the enhanced IgA response remain to be established.

Muc2^DTA/+^ mice developed glomerular IgA deposition, more pronounced under SPF than germ-free conditions, yet kidneys were histologically normal in both SPF and GF conditions. Human IgA nephropathy is driven by galactose-deficient IgA1 and mice lack the IgA1 subclass, so murine models reproduce the disease only partially, and C57BL/6 is relatively resistant to glomerular injury (*38*). The Muc2^DTA/+^ model therefore captures mostly the pre-renal phase of the gut-kidney axis. We provide evidence that a primary secretory barrier defect is sufficient to generate the mucosal and circulating IgA abnormalities that precede IgA nephropathy, but not to cause direct overt renal disease.

Together, these findings place the goblet cell/mucus compartment upstream of an IgA regulatory pathway that is separable from microbiota-driven IgA inductions. The microbiota determines much of the intestinal inflammatory phenotype and reshapes the accompanying B-cell response, but is not required for mucosal or systemic hyper-IgA. More broadly, the results support a model in which intestinal epithelial lineages may influence humoral immune responses with consequences extending beyond the intestine and may contribute to the development of IgA-driven disease such as IgAN.

## Materials & Methods

### Development of the Muc2iCre mouse line

This Muc2iCre line was created via bacterial artificial chromosome (BAC) transgenesis using a rat derived BAC (CH230-244C22) containing the *Muc2* gene locus. BAC CH230-244C22 containing the rat Muc2 locus was obtained from the Children’s Hospital Oakland Research Institute (CHORI) (https://bacpacresources.org/). Briefly, the iCre recombinase gene, along with an FRT-kan/neo-FRT selection cassette, was inserted at the start codon of the *Muc2* gene using RedET recombination. Next, bacteria containing the recombined BAC clone were subsequently transformed with a Flpe plasmid to remove the selection cassette. The recombined BAC was then digested using restriction enzymes (PvuI and SacII) to release the relevant 93 kB Muc2iCre fragment while removing the adjacent *Muc5AC* locus from the BAC. The MUC2 iCre fragment was isolated by pulse field gel electrophoresis, electro eluted, purified using an Amicon Ultra-4 (Ultracel-100K) centrifugal filter (Millipore) and injected into C57BL/6J oocytes to allow random genomic integration. Embryos were subsequently transferred into the oviducts of pseudopregnant female mice, after which founder mice were screened for the presence of the transgene by PCR and characterized for cell-specific expression. The genomic integrations site and copy number of the Muc2iCre fragment was investigated through Targeted Locus Amplification (TLA) technology and revealed integration in the mouse chromosome 3 with an estimated copy-number between 2 and 11.

### Identification of vector integration sites by Cergentis

1 transgenic mouse with the vector injection fragment MUC2-iCre sequence was analysed by Cergentis. An overview of the Targeted Locus Amplification (TLA) technology and technical details can be found in the Addendum. TLA was performed using 3 independent primer sets specific for the vector sequence (Table 1). The vector was found to be in mouse chromosome 3: 133,465,565-133,468,815. According to the RefSeq the integrity of Tet2 may have been affected. The copy number is estimated to be 2-11 copies.

**Table 1:**
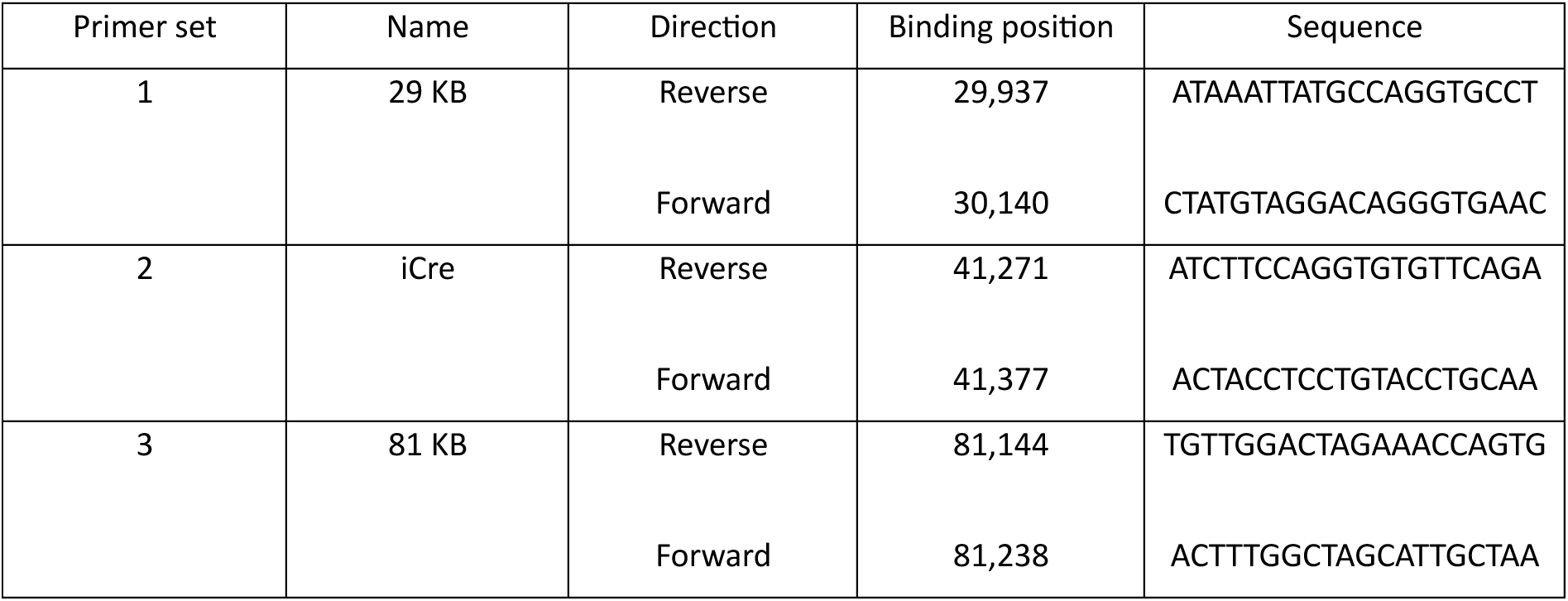
Primer sets specific for the vector Muc2iCre sequence.

### Animal experiments

Mice were housed in individually ventilated cages in a specific pathogen-free facility at the VIB Center for Inflammation Research. All experiments used mice on a C57BL/6 background. Animal experiments were approved by the Ethical Committees for Animal Experimentation of the Faculty of Medicine and Health Sciences (ECD21-76) and the Faculty of Sciences (WE14) of Ghent University (LP0014, LN1187, EC2021-006, EC2022-013, EC2024-045) and complied with national and European regulations.

#### DT injection

Muc2^DTR^ mice received diphtheria toxin (1 µg in 200 µl PBS) intraperitoneally on days −1 and 0.

#### Colonoscopy

Mice were anesthetized with isoflurane (4.5% for induction, 2.5% in oxygen for maintenance; Medini) and examined with a Karl Storz endoscope. Recorded colonoscopies were scored for colonic inflammation using the Murine Endoscopic Index of Colitis Severity (MEICS) (*29*).

#### FITC-dextran assay

Mice received FITC-dextran (FD4, 3–5 kDa; Sigma-Aldrich) by oral gavage (600 mg/kg). After 4 h, blood was collected by cardiac puncture and centrifuged (8,000 × g, 8 min) and 100 µl undiluted serum was transferred to a 96-well plate. FITC-dextran was measured fluorometrically on a FLUOstar Omega plate reader (BMG Labtech) against a standard curve and concentrations were normalized to the mean of PBS-gavaged control mice.

#### Antibiotics

Mice received vancomycin (500 mg/L), metronidazole (1 g/L), ampicillin (1 g/L) and neomycin (200 mg/L) in the drinking water ad libitum for 6 weeks, starting at 11 weeks of age.

#### Trichuris muris infection

Mice were infected by oral gavage with 200 *T. muris* eggs. At the peak of infection (day 10), mice were euthanized and colon tissue was collected for analysis.

### Histology

Tissues were fixed in 4% formaldehyde for 24 h and paraffin-embedded unless stated otherwise. Brightfield and immunofluorescence slides were imaged with a Zeiss Axioscan and Zen Blue software unless stated otherwise. For antigen retrieval, dewaxed sections were boiled for 20 min in citrate buffer (Labconsult; VECH-3300, 1:100) in a PickCell Electric Cooker and cooled for 2 h.

#### H&E (paraffin sections)

Sections were stained with hematoxylin and eosin (H&E) to evaluate inflammation. Crypt length was measured at 3 locations per sample (10 crypts per location, 30 measurements per sample) and the sample average was plotted.

#### Alcian-Blue Periodic Acid Schiff’s (AB-PAS) (paraffin sections)

Dewaxed sections were stained with 1% Alcian blue (20 min), oxidized with 1% periodic acid (Sigma; 395132; 15 min) and incubated with Schiff’s reagent (Sigma; 395-2-016; 10 min). Sections were counterstained with Mayer’s hematoxylin (30 s), dehydrated and mounted. Acid mucins stain blue and neutral mucins magenta.

#### Anti-GFP and anti-Ki67 (paraffin sections)

Colon, small intestine, kidney, lung, liver, cervix and stomach were stained for GFP and gut for Ki67. After antigen retrieval, endogenous peroxidase was blocked with 3% H₂O₂ in methanol (10 min) and sections were blocked with 5% goat serum in 1% BSA/PBS (30 min). Sections were incubated overnight at 4 °C with anti-GFP (Cell Signaling; 2956, 1:200) or anti-Ki67 (Cell Signaling; 12202, 1:1000), then for 1 h with biotinylated goat anti-rabbit IgG (DAKO; E0432, 1:500) and for 30 min with Vectastain Elite ABC-HRP (Vector; PK-7100). Signal was developed with the ImmPACT DAB Substrate Kit (Vector; SK-4105) and sections were counterstained with hematoxylin.

#### Anti-Muc2 (paraffin sections)

After antigen retrieval, sections were blocked for 30 min with 1% goat serum in PBS containing 0.5% BSA (Sigma-Aldrich; A2153) and 0.1% Tween-20 (Sigma-Aldrich; P1379). Sections were incubated overnight at 4 °C with anti-Muc2 (Abcam; ab272692, 1:200), then for 1 h with goat anti-rabbit DyLight 488 (1:1000) and Hoechst (1:1000).

#### FISH (paraffin sections)

Sections were stained as described previously (*39*) with the universal bacterial probe EUB338 (1 µg; 5’-Cy3), DAPI (1:1000), UEA-1 (Labconsult; VEC.FL-1061-5, 1:1000) and WGA (Invitrogen; W6748, 1:200). Images were acquired on a Zeiss LSM880 Airyscan (40×) and processed with Zen Blue software.

#### Anti-pIgR (paraffin sections)

After antigen retrieval with Vector antigen unmasking solution (1:100), sections were blocked for 1 h at RT in PBS with 1% donkey serum and incubated overnight at 4 °C with rabbit anti-Muc2 (Santa Cruz; H-300, 1:200) and goat anti-pIgR (R&D Systems; AF2800, 1:200). Sections were then incubated for 2 h at RT with donkey Alexa Fluor Plus 488 anti-rabbit and Alexa Fluor 568 anti-goat secondary antibodies, counterstained with DAPI (1:1000, 15 min) and mounted in PVA-DABCO.

#### Anti-IgA and anti-IgG (frozen sections)

Kidneys were fixed in 4% PFA for 1 h, cryoprotected overnight in 30% sucrose and embedded in OCT. Cryosections (10µm) were rehydrated in PBS (5 min) and blocked for 1 h with 1% donkey serum, 1% FCS, 2% BSA and 0.3% Triton X-100 in PBS. Sections were incubated overnight at 4 °C with rat anti-mouse IgA (BD; 556960, 1:100), washed in PBS, and incubated for 1 h at RT with donkey anti-rat Alexa Fluor 488 (1:500). Sections were then blocked for 1 h at RT with 1% goat serum, 1% FCS, 2% BSA and 0.3% Triton X-100 in PBS and incubated for 1 h at RT with DyLight 650-conjugated anti-mouse IgG (Abcam; ab98715, 1:200) and DAPI (1:1000). Sections were washed and mounted in ProLong Diamond Antifade Mountant (Invitrogen).

### SEM

SEM imaging was performed on wild-type (n=2) and Muc2DTA (n=2) surface intestinal epithelium. Tissues were dissected, cut longitudinally and placed on whatman filter paper in fixation solution (2% PFA, 2.5% Glutaraldehyde in 0.1M Na-Cacodylate pH7.4). After overnight fixation, the samples were washed in 0.1M Na-Cacodylate, then incubated for 30 minutes in 2% OsO4 in 0.1M Na-Cacodylate. Next, samples were washed with H2O before incubating them for 15 minutes in subsequently 50, 70, 85, 95 and 100% EtOH. After these incubation steps, samples were treated with EtOH-Aceton (1:1) solution for 15 minutes. Thereafter, samples were incubated in 100% aceton for 15 minutes and dried by Critical Point Drying technique using Leica EM CPD300. Afterwards, samples were coated with platinum (15nm Pt) using the Quorum Q150T Plus sputtercoater. Images were required using Zeiss Crossbeam 540 FIB-SEM, with a voltage of 5kV.

### RNA isolation and quantitative real-time PCR (qPCR)

Tissue was lysed and homogenized using RLT and β−mercaptoethanol and the TissueLyser II (Qiagen). Total RNA was isolated using the RNeasy Mini Kit (Qiagen; 74106), according to the manufacturer’s instructions.

The synthesis of cDNA was performed using QuantiTect Reverse Transcription Kit (Qiagen; 205311), following the manufacturer’s protocol. For qPCR, SensiFAST SYBR NO-ROX (BioLine; BIO-98005) and were used on LightCycler 480 (Roche). The reactions were performed in triplicates and the results were analyzed with qbase+ software (Biogazelle) and indicated as calibrated normalized relative quantity (CNRQ) for gene expression profiling.

As housekeeping genes, Actb, Ubc, Hprt1, Villin and Rpl13a were used.

### Single cell and bulk RNA sequencing

Mice were euthanized by cervical dislocation and the colon and ileum were dissected and flushed with cold PBS. The tissue was cut into small pieces and incubated in 12.5 mL 20 mM EDTA in PBS for 90 min at 4 °C on a rotating mixer. The supernatant (fraction 1) was collected by vigorous shaking and centrifuged (400 g, 5 min) and the pellet was resuspended in PBS with 2% FCS. The tissue was incubated twice more in 12.5 mL 20 mM EDTA in PBS for 30 min at 4 °C and fractions 2 and 3 were collected in the same way. Fractions were pooled and digested for 10 min at 37 °C with shaking in 10 mL prewarmed 0.8 mg/mL Dispase II in HBSS. The digestion was stopped with 10 mL 10% FCS, and cells were filtered through a 100 µm filter. Debris was removed by 30% Percoll (Cytiva; 17-0891-02) in PBS and cells were stained with DAPI (1:1000). Live (DAPI⁻) cells were sorted as follows: for single-cell RNA sequencing, 100,000 cells from Muc2^RFP/+^ ileum and colon (80% RFP⁺ and 20% RFP⁻); for bulk and single-cell RNA sequencing, 100,000 YFP⁺ cells from Muc2^YFP/+^ colon.

### Bulk RNA sequencing

#### Muc2^YFP^

Sorted YFP+ and YFP-cells from the colon lamina propria were analysed with bulk RNA sequencing using a QuantSeq mRNA Library Prep and sequencing on the NextSeq500 SR-75 by NXTGNT (Faculty of Pharmaceutical Sciences, Ghent University). Raw FASTQ files were processed with the nf-core/rnaseq pipeline (version 3.14.0) (*40*, *41*). Default parameters were used, with the following modifications:

UMI Handling:

- --with_umi,
- --umitools_extract_method “regex”,
- --umitools_bc_pattern “^(?P<umi_1>.{6})(?P<discard_1>.{4}).*”

Alignment Parameters:

- --alignIntronMax 1000000,
- --alignIntronMin 20,
- --alignMatesGapMax 1000000,
- --alignSJoverhangMin 8,
- --outFilterMismatchNmax 999,
- --outFilterMultimapNmax 20,
- --outFilterType BySJout,
- --outFilterMismatchNoverLmax 0.1,
- --clip3pAdapterSeq AAAAAAAA

#### Muc2^DTA^

Poly(A)-enriched, strand-specific (dUTP) RNA-seq libraries were prepared and sequenced by Novogene (Cambridge, UK) using their in-house library prep kit. Sequencing was performed on an Illumina NovaSeq X Plus to generate 150 bp paired-end reads.

Raw sequencing data were processed using nf-core/rnaseq v3.21.0 (*42*) from the nf-core collection of workflows (*41*), utilising reproducible software environments provided by Bioconda (*43*) and Biocontainers (*44*). The pipeline was executed using Nextflow v25.04.6 (*40*). Transcripts were mapped to the mouse reference genome GRCm39 (GCA_000001635.9) using Ensembl annotation release 112. Sequencing and alignment quality metrics were assessed using the quality-control tools implemented within the nf-core/rnaseq workflow using default settings.

Salmon transcript abundance estimates generated by the pipeline were imported into R (v4.6.0) (*45*) using tximport (v1.40.0) (*46*) and summarised to gene-level counts for downstream analyses. The experimental design and sample relationships were assessed using principal component analysis (PCA) performed on variance-stabilized count data generated using DESeq2 (v1.52.0) (*47*). Differential gene expression analysis was conducted using DESeq2. Genes with an adjusted P-value ≤ 0.05 and an absolute log2 fold change ≥ 1 were considered differentially expressed. To identify genes with a genotype-by-housing interaction, a model including genotype, housing and their interaction was fitted in DESeq2, and genes with an adjusted P-value < 0.05 and an absolute interaction log2 fold change ≥ 0.5 were considered to show a significant interaction.

#### Muc2^YFP^ and Muc2^DTA^

Gene set enrichment analysis (GSEA) and over-representation analysis (ORA) were performed using the clusterProfiler package (v4.20.0) (*48*, *49*). Gene set annotations were retrieved from the msigdbr package (v26.1.0) (*50*) for Mus musculus. The following mouse MSigDB (v2025.1.Mm) (*51–54*) collections were included: Gene Ontology terms: Biological Processes, Cellular Components and Molecular Functions (M5); canonical pathways from Reactome (CP:REACTOME), WikiPathways (CP:WIKIPATHWAYS) and BioCarta (CP:BIOCARTA), and chemical and genetic perturbations (CGP) (M2); Hallmark gene sets (MH); transcription factor target gene sets from the GTRD subcollection (M3) (*55*); and immunologic signature gene sets (M7).

ORA was performed separately for up- and downregulated genes, using all tested genes as background. For GSEA, differential gene expression (DGE) results were shrunk using the ashr package (v2.2.63) (*56*) to reduce variance associated with lowly expressed genes. A preranked gene list was constructed by ordering genes based on their shrunken log2 fold change. For both ORA and GSEA, only gene sets containing between 15 and 500 genes (*57*) were included to ensure statistical robustness and biological interpretability. Significantly enriched pathways were defined by a Benjamini–Hochberg adjusted P-value < 0.05 and a q-value < 0.2 for ORA, and an adjusted P-value < 0.05 for GSEA.

Transcription factor activity was inferred with the decoupleR package (v2.17.0) (*58*), using a univariate linear model (ULM) and the mouse CollecTRI regulatory network (*59*). Ligand activity predictions were performed using nichenetr (v2.2.1.1) (*60*) with the NicheNet v2 mouse prior model (*61*), and multinichenetr (v2.1.0) (*62*). Figures were generated using ggplot2 (v4.0.3) (*63*).

### Single cell RNA sequencing

#### Muc2^YFP^

Sequencing libraries were prepared and loaded onto an Illumina NovaSeq flow cell at the VIB Nucleomics Core, following the manufacturer’s recommended protocols provided by 10x Genomics. The Cell Ranger software suite (version 6.0.0, 10x Genomics (*64*)) was used to process the raw sequencing data, generating FASTQ files. These files were then aligned to the mouse reference genome Gencode Release M24 (GRCm38.p6) (*65*) to produce gene expression matrices. Subsequent processing steps, including barcode demultiplexing, unique molecular identifier (UMI) filtering and gene counting, were performed using Cell Ranger (v6.0.0, 10x Genomics (*64*)) and Seurat (v5.1.0, (*66*)). For downstream analyses, the filtered feature-barcode matrix produced by Cell Ranger (v6.0.0, 10x Genomics, (*64*)) was used. Additional quality control filtering was applied to retain barcodes with mitochondrial content below 30%, a minimum of 500 counts and more than 300 detected features. Following initial clustering, a small subset of tuft and immune cells was excluded to focus the analysis exclusively on goblet cells. Gene signature scoring in single-cell datasets was performed using UCell (*67*). The specific marker genes used for these analyses are provided in the Supplementary Table 1. Cell-type proportions were calculated using get_abundance_info function (*62*).

#### Muc2^RFP^

Sequencing libraries were prepared and loaded onto an Illumina NovaSeq flow cell at the VIB Nucleomics Core, following the manufacturer’s recommended protocols provided by 10x Genomics. The Cell Ranger software suite (version 6.0.0, 10x Genomics, (*64*)) was used to process the raw sequencing data, generating FASTQ files. These files were then aligned to the mouse reference genome Gencode Release M24 (GRCm38.p6) (*65*) to produce gene expression matrices. Subsequent processing steps, including barcode demultiplexing, unique molecular identifier (UMI) filtering and gene counting, were performed using Cell Ranger (v6.0.0, 10x Genomics, (*64*)) and Seurat (v5.1.0, (*66*)). For downstream analyses, we utilized the filtered feature-barcode matrix produced by Cell Ranger (v6.0.0, 10x Genomics, (*64*)). Gene signature scoring in single-cell datasets was performed using UCell (*67*). The specific marker genes used for these analyses are provided in Supplementary Table 1.

### DNA extraction of stools

DNA was extracted from fecal samples using the RNeasy PowerMicrobiome Kit (Qiagen; 26000-50-2), according to manufacturer’s instructions.

### 16S rRNA sequencing

Full-length (V1–V9, ∼1.5 kbp) 16S rRNA gene sequences were obtained using PacBio HiFi sequencing. Data was processed using nf-core/ampliseq v2.15.0 (*68*, *69*) of the nf-core collection of workflows (*41*), utilising reproducible software environments from the Bioconda (*43*) and Biocontainers (*44*) projects. The pipeline was executed with Nextflow v25.04.6 (*40*) using Singularity containers and default parameters, with the following modifications to accommodate the PacBio sequencing data and specific primers: Forward Primer (27F): AGRGTTYGATYMTGGCTCAG, Reverse Primer (1492R): RGYTACCTTGTTACGACTT, the --pacbio option to specify single-end PacBio reads, and --sample_inference pseudo to enable pseudo-pooled sample inference. Read quality was assessed with FastQC v0.12.1 (*70*) and MultiQC v1.29 (*71*). Primers were removed with Cutadapt v4.6 (*72*), and reads without primers were discarded. DADA2 v1.30.0 (*73*) was used to quality-filter reads (maximum 2 expected errors, minimum length 50 bp), denoise them with the PacBio error model (*74*) and remove chimeras. Mitochondrial and chloroplast ASVs were removed. Taxonomic classification was performed with the DADA2 naïve Bayesian classifier (minimum bootstrap confidence of 50 out of 100) against the SILVA 138.2 prokaryotic SSU database (*75*, *76*). Species were assigned by exact sequence matching against the same database.

Data were analyzed in R v4.6.2 (*45*) with Bioconductor v3.23. The pipeline output was imported with phyloseq v1.56.0 (*77*) and TreeSummarizedExperiment v2.20.0 (*78*) and analyzed with mia v1.20.0 (*79*). Figures were made with miaViz v1.20.0 (*80*), scater v1.40.2 (*81*), ggplot2 v4.0.3 (*63*) and ggsignif v0.6.4 (*82*). Mucosa and stool samples were analyzed separately. ASVs were kept if they reached at least 0.1% relative abundance in at least 10% of the samples. Alpha diversity (observed richness, Shannon, inverse Simpson, Gini–Simpson and Pielou evenness) was calculated on unfiltered ASV counts rarefied to the smallest library size, and groups were compared with Wilcoxon rank-sum tests. Beta diversity was calculated with Bray–Curtis and Aitchison distances (*83*), visualized by PCoA and tested by PERMANOVA (adonis2, vegan v2.7.5, 9999 permutations) (*84*, *85*). Sex (mucosa) or cage (stool) was included as a covariate. Differential abundance was tested with MaAsLin2 v1.26.0 (*86*) at the family, genus and species level. Taxa with at least 10% prevalence were included, and counts were TSS-normalized and log2-transformed before fitting linear models with the same covariates. P-values were adjusted with the Benjamini–Hochberg method (*87*), and q ≤ 0.25 was considered significant.

### Absolute IgA coating

IgA coating of fecal bacteria was assessed by flow cytometry as previously described (*9*). Fresh fecal pellets were weighed and homogenized in sterile HBSS (1 mL per 100 mg) by vortexing and debris was removed by centrifugation (400 × g, 5 min). The bacterial supernatant was diluted in HBSS to ∼10⁶ bacteria per well (assuming ∼10⁹ CFU/g stool) in a V-bottom plate and washed twice with HBSS. Bacteria were stained for 30 min at 4 °C in the dark with PE-conjugated anti-mouse IgA (eBioscience; clone mA-6E1, 12-4204-82; 1:250) and SYTO9 (Invitrogen; 1:1000) in 100 µL, then washed twice and resuspended in HBSS. Samples were acquired on a NovoCyte flow cytometer and analyzed in FlowJo. Unstained, PE-only and SYTO9-only controls, together with germ-free and IgA-knockout fecal samples, were included in each run. IgA-coated bacteria were defined as the SYTO9⁺PE⁺ population (Fig. S9B).

### Magnetic separation of IgA-coated fecal bacteria

IgA-coated bacteria were isolated as previously described (*88*). Fecal pellets stored at −20 °C were resuspended in 500 µl ice-cold PBS, homogenized and shaken (20 min, 1,100 rpm., 4 °C). After adding 2× volume of PBS, debris was sedimented (100 × g, 3 min, 4 °C). The supernatant was passed through a 70 µm sieve, bacteria were pelleted (10,000 × g, 5 min, 4 °C) and resuspended in 1 ml PBS. Bacterial density was estimated from OD600 (OD600 = 1 was assumed to equal 5 × 10⁸ bacteria/mL). For each sample, 10⁹ bacteria were blocked in 5% goat serum in PBS (20 min, on ice), washed and stained with 4 µg FITC-conjugated goat anti-mouse IgA (Abcam; ab97234) in 100 µl PBS (30 min, 4 °C, 800 rpm.). Stained bacteria were washed, incubated with anti-FITC microbeads (Miltenyi; 10 µl per 10⁹ bacteria; 15 min, 4 °C) in 5% goat serum/PBS and resuspended in 5 ml of the same buffer. They were then passed over an LD column (Miltenyi; 130-042-901) and the flow-through was collected as the IgA⁻ fraction. The retained IgA⁺ fraction was eluted and passed over an LS column (Miltenyi; 130-042-401). The LS flow-through was pooled with the IgA⁻ fraction and the LS eluate was pooled with the IgA⁺ fraction. Fractions were pelleted by ultracentrifugation and the pellets were stored at −20 °C until DNA isolation.

### Shotgun metagenomics

Shotgun metagenomic libraries were prepared and sequenced by Novogene on an Illumina NovaSeq X Plus platform, generating 2 × 150 bp paired-end reads. Data was processed using nf-core/taxprofiler v1.2.5 (*89*) of the nf-core collection of workflows (*41*), utilising reproducible software environments from the Bioconda (*43*) and Biocontainers (*44*) projects. The pipeline was executed with Nextflow v25.04.6 (*40*) using Singularity containers and default parameters, with short-read quality control, low-complexity filtering, host read removal, lane merging and Bracken enabled.

Read quality was assessed with FastQC v0.12.1 (*70*) and MultiQC v1.32 (*71*). Reads were trimmed with fastp v0.24.0 (*90*), and low-complexity reads were removed with BBDuk v39.18 (*91*). Host reads were removed by mapping to the mouse genome (GRCm39) with Bowtie2 v2.5.4 (*92*) and SAMtools v1.21 (*93*). Reads were classified with Kraken2 v2.1.5 (confidence 0.1) (*94*) against the PlusPF database (October 2025) (*95*). Species abundances were estimated with Bracken v3.0.1 (read length 150 bp) (*96*). Profiles were merged with TAXPASTA v0.7.0 (*97*) using the NCBI Taxonomy (*98*).

Data were analysed in R v4.6.2 (*45*) with mia v1.20.0 (*79*) and TreeSummarizedExperiment v2.20.0 (*78*). Only bacterial species with at least 10 reads in at least 5% of the samples were kept. Counts were summed at the family level and converted to relative abundance. Mean relative abundances per genotype and IgA fraction were shown as a stacked barplot with ggplot2 v4.0.3 (*63*). Families were grouped and colored by phylum, showing the six most abundant phyla and up to four families per phylum (mean relative abundance ≥ 0.5%).

### ELISA

Stool samples were weighed, homogenized in 500 µl sterile PBS and centrifuged (30 min, full speed, 4 °C) and supernatants were stored at −20 °C (short term) or −80 °C (long term). Blood was collected postmortem by cardiac puncture and centrifuged (8,000 × g, 8 min) and serum was stored at −20 °C. Unless stated otherwise, plates were developed with TMB substrate (Invitrogen; 12750000), the reaction was stopped with H₂SO₄ and absorbance was read at 450 nm with wavelength correction (570 nm; 650 nm for total Ig ELISAs). Incubations were at RT.

#### Fecal IgA

Fecal IgA was quantified with a mouse IgA ELISA kit (Invitrogen; 88-50450) according to the manufacturer’s instructions. Concentrations were interpolated from the standard curve and normalized to stool weight.

#### Fecal lipocalin-2

Fecal lipocalin-2 was quantified with the Mouse Lipocalin-2/NGAL DuoSet ELISA (R&D Systems; DY1857) according to the manufacturer’s instructions, using stool supernatants at 1:10 and 1:100 dilutions in reagent diluent (1% BSA in PBS). Concentrations were interpolated from the standard curve and normalized to stool weight.

#### Total serum IgA, IgM and IgG

Plates were coated overnight at 4 °C with anti-mouse IgA (BD; 556969, 1:500), anti-mouse IgM (BD; 553435, 1:500) or anti-mouse IgG Fab (Abcam; ab197774, 1:1,000) and blocked with 1% casein for 2 h. Serum diluted in assay buffer (0.1% casein; 1:400 for IgA, 1:32,000 for IgM and IgG) was incubated for 2 h, followed by HRP-conjugated goat anti-mouse IgA (SouthernBiotech; 1040-05, 1:5,000), IgM (SouthernBiotech; 1021-05, 1:5,000) or IgG-Fc (Abcam; ab98741, 1:40,000) for 1 h. Concentrations were calculated from standard curves.

#### IgA-IgG complexes

Plates were coated overnight at 4 °C with anti-mouse IgA (BioLegend; clone RMA-1, 407001; 1 µg/mL) and blocked with 2% BSA in PBS for 1 h. Serum diluted in 2% BSA/PBS was incubated for 2 h and captured IgG was detected with HRP-conjugated donkey anti-mouse IgG (R&D Systems; HAF018; 1:5,000) for 2 h. Results are reported as OD450.

#### J chain-containing IgA

Assays were performed as previously described (*99*). Plates were coated overnight at 4 °C with anti-J chain antibody (Proteintech; 13688-1-AP, 1:3,000) and blocked with 1% BSA in PBS for 1 h. Serum diluted in PBST was incubated for 90 min, and captured IgA was detected with HRP-conjugated goat anti-mouse IgA (Abcam; ab97235, 1:2,000) for 1 h. Results are reported as OD450.

#### Serum cytokines

TNF-α, IL-6, IL-1β, MCP-1, IFN-γ and IL-17 were measured in serum using the Bio-Plex multiplex immunoassay system (Bio-Rad) according to the manufacturer’s instructions.

### Flow cytometry

Mice were euthanized by cervical dislocation and tissues were processed as follows. Cells (1 × 10⁶/sample) were stained for surface and, where indicated, intracellular markers of T cells and myeloid cells in separate panels (Table 3). For intracellular staining, cells were fixed and permeabilized after surface staining with the Foxp3 Transcription Factor Staining Buffer Set (eBioscience; 00-5523-00). Samples were acquired on a five-laser BD LSRFortessa, except lung samples (BD FACSymphony A5, five lasers) and B1 cells (BD FACSDiscover A8 Cell Analyzer). Data were analyzed with FlowJo v10.8.1 (BD Biosciences). Gating strategies are shown in Supplementary Figs. S9A, S10 and S11.

**Table 2:**
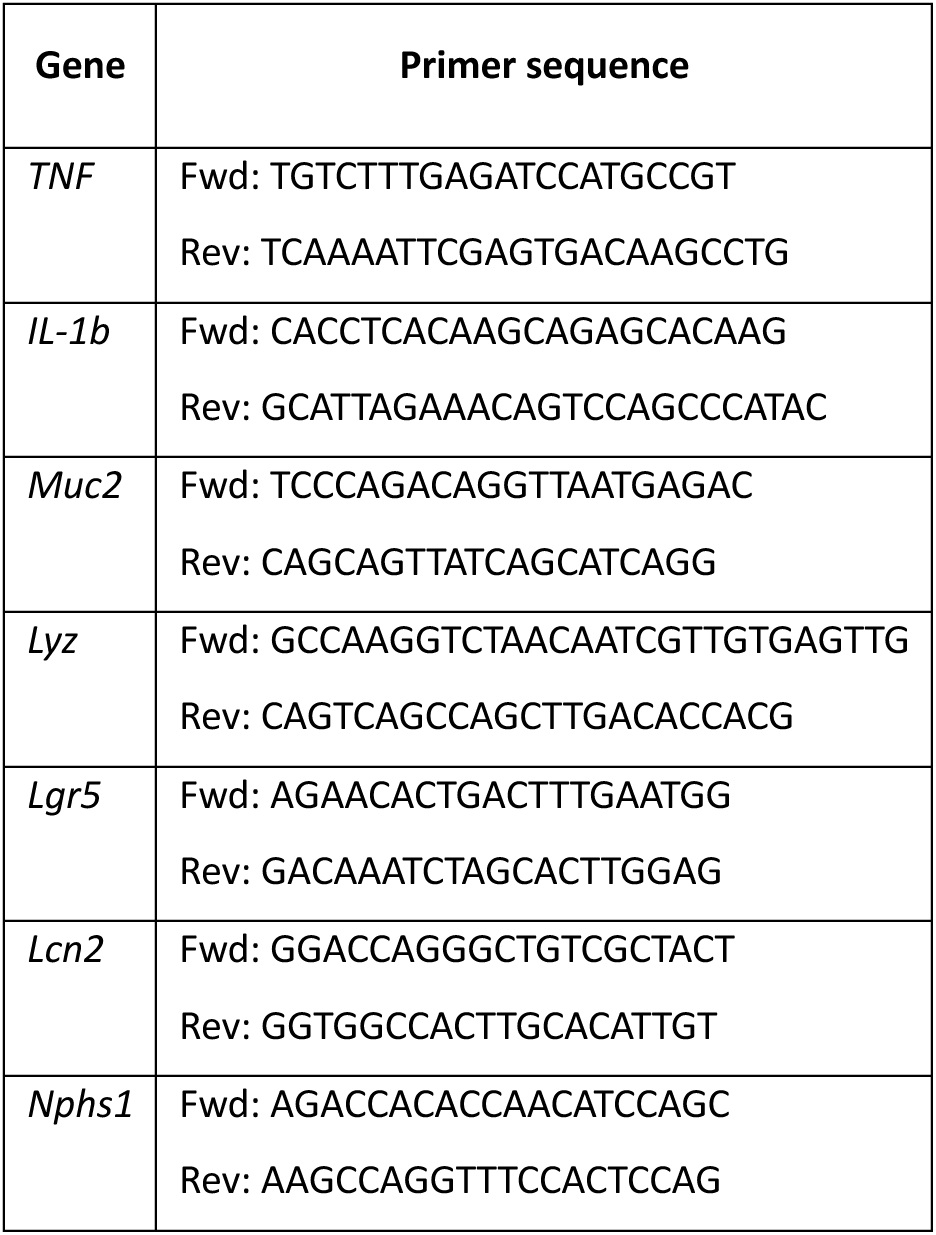
Primer sequences used for qPCR analysis of duodenal, jejunal, ileal, colonic and kidney tissue.

**Table 3:** Antibodies used for flow cytometry.

| **Small intestine, large intestine, synovium, spleen** |  |  |
|---|---|---|
| Myeloid panel |  |  |
| Fixable Viability Dye eFluor 506 (live/dead) | eBioscience; 65-0866-14 | 1:300 |
| CD19 | eBioscience; 15-0193-82 | 1:400 |
| CD3 | eBioscience; 15-0031-82 | 1:200 |
| NK1.1 | BioLegend; 108716 | 1:200 |
| anti-CD45 Alexa Fluor 700 | eBioscience 56-0451-82 | 1:800 |
| anti-Ly6G PercpCy5.5 | BD; 560602 | 1:200 |
| anti-Ly6C-APC | eBIoscience; 17-5932-80 | 1:200 |
| anti-Siglec F BUV395 | BD; 740280 | 1:200 |
| anti-CD11b BV506 | BD; 563015 | 1:600 |
| anti-CD64 BV711 | BioLegend; 139311 | 1:100 |
| anti-F4/80 Biotin | eBioscience; 13-4801-82 | 1:100 |
| Streptavidin BV421 | BioLegend; 405226 | 1:1000 |
| anti-CD11c PE-eFluor 610 | eBioscience; 61-0114-82 | 1:300 |
| anti MHC class II APC-eFLuor 780 | eBioscience; 47-5321-80 | 1:800 |
| anti-XCR1 BV650 | Biolegend, 148220 | 1:400 |
| anti-SIRPa PECy7 | Biolegend, 144007 | 1:100 |
| T cell panel |  |  |
| 7-AAD (live/dead) | BD; 559925 | 1:1000 |
| anti-CD3 APC | eBioscience; 17-0031-83 | 1:100 |
| antiCD4 APC Cy-7 | BD; 552051 | 1:200 |
| anti-CD8 V500 | BD; 560776 | 1:100 |
| anti-Foxp3 Alexa Fluor 488 | eBioscience; 53-5773-82 | 1:100 |
| anti-RORct BV421 | BD; 562894 | 1:100 |
| anti-Tbet PE-Cy7 | Invitrogen; 25-5825-82 | 1:100 |
| anti-GATA3 PE | Invitrogen; 12- 9966-42 | 1:100 |
| **Lung** |  |  |
| Myeloid panel |  |  |
| eFluor506 (live/dead) | eBioscience; 65-0866-14 | 1:200 |
| anti-CD26 FITC | BioLegend; 137805 | 1:100 |
| anti-XCR1 BV421 | BioLegend; 148216 | 1:400 |
| anti-Ly6G BV650 | BD; 740554 | 1:400 |
| anti-CD64 BV711 | BioLegend; 139311 | 1:100 |
| anti-Siglec F BV786 | BD; 740956 | 1:400 |
| anti-CD45 APC | BioLegend; 103112 | 1:200 |
| anti-Ly6C AF700 | BD; 561237 | 1:300 |
| anti-MHCII APC-Cy7 | eBioscience; 47-5321-82 | 1:500 |
| anti-CD101 PE | eBioscience; 12-1011-82 | 1:800 |
| anti-CD11c PE-TR | eBioscience; 61-0114-82 | 1:300 |
| anti-CD3 PE-Cy5 | BioLegend; 100310 | 1:200 |
| anti-CD19 PE-Cy5 | eBioscience; 15-0193-83 | 1:400 |
| anti-CD206 PE-Cy7 | BioLegend; 141720 | 1:200 |
| anti-CD11b BUV395 | BD; 563553 | 1:200 |
| anti-MerTK Biotin (UV496) Streptavidin | Bio-Techne; BAF591 | 1:100 |
| anti-CD172a BUV661 | BD; 741593 | 1:200 |
| anti-CD43 BUV737 | BD; 612840 | 1:200 |
| T cell panel |  |  |
| APC-Cy7 (live/dead) | Invitrogen; 65-0865-14 | 1:400 |
| anti-CD16/anti-CD32 | Bioceros; 2.4G2 | 1:200 |
| anti-CD8 FITC | BioLegend; 100706 | 1:50 |
| anti-IFNγ PerCP-Cy5.5 | BioLegend; 505822 | 1:200 |
| anti-IL-13 BV421 | eBioscience; 48-7133-82 | 1:150 |
| anti-CD62L BV510 | BD; 563117 | 1:100 |
| anti-IL-17 BV605 | BioLegend; 506927 | 1:150 |
| anti-CD44 BV650 | BioLegend; 103049 | 1:200 |
| anti-Siglec F BV786 | BD; 740956 | 1:400 |
| anti-GATA3 APC | eBioscience; 50-9966-42 | 1:250 |
| anti-CD3 AF700 | BioLegend; 100216 | 1:100 |
| anti-RORγt PE | eBioscience; 12-6988-82 | 1:50 |
| anti-CD4 PE-eFluor610 | eBioscience; 61-0042-82 | 1:400 |
| anti-NK1.1 PE-Cy5 | BioLegend; 108716 | 1:250 |
| anti-TCRγδ | eBioscience; 15-5711-81 | 1:250 |
| anti-B220 PE-Cy5 | BioLegend; 103210 | 1:400 |
| anti-Ter119 PE-Cy5 | BioLegend; 116210 | 1:200 |
| anti-CD19 PE-Cy5 | eBioscience | 1:400 |
| anti-CD11b PE-Cy5 | BioLegend; 101210 | 1:800 |
| anti-T-bet PE-Cy7 | eBioscience; 25-5825-82 | 1:50 |
| anti-CD90 BUV496 | BD; 741047 | 1:200 |
| anti-ST2 BUV661 | BD; 750671 | 1:200 |
| anti-CD127 BUV737 | BD; 612841 | 1:200 |
| anti-CD45 BUV805 | BD; 568336 | 1:1000 |
| **Mesenteric lymph nodes, Peyer's patches, colon, ileum** |  |  |
| B cell panel |  |  |
| APC-Cy7 (live/dead) | Invitrogen; 65-0865-14 | 1:400 |
| anti-CD16/anti-CD32 | Bioceros; 2.4G2 | 1:200 |
| anti-IgA AF488 | Abcam; ab9701 | 1:400 |
| anti-TACI Pacific Blue | BD; 742840 | 1:200 |
| anti-IgM biotin | BD; 553406 | 1:400 |
| Streptavidin BV605 | BD; 563260 | 1:400 |
| anti-CD138 BV711 | BioLegend; 142519 | 1:100 |
| anti-CD19 BV786 | BD; 563333 | 1:200 |
| anti-GL7 APC | BioLegend; 144606 | 1:200 |
| anti-IgG1 AF700 | BioLegend; 406632 | 1:100 |
| anti-CD80 PE-Cy5 | BioLegend; 104712 | 1:100 |
| anti-CD95 PE-Cy7 | BD; 557653 | 1:200 |
| anti-CD273 BUV379 | BD; 565102 | 1:200 |
| anti-CD3e BUV737 | BD; 612771 | 1:200 |
| **Peritoneal fluid, spleen, colon** |  |  |
| B1 cell panel |  |  |
| APC-Cy7 (live/dead) | Invitrogen; 65-0865-14 | 1:400 |
| anti-CD16/anti-CD32 | Bioceros; 2.4G2 | 1:200 |
| anti-FcRL5 AF488 | R&D Systems; FAB6757G | 1:200 |
| anti-IgM PerCP-Cy5.5 | BD; 550881 | 1:200 |
| anti-CD21:CD35 eFluor 450 | eBioscience; 48-0212-82 | 1:200 |
| anti-B220 BV605 | BioLegend; 103244 | 1:200 |
| anti-CD5 BV786 | BD; 740842 | 1:100 |
| anti-Blimp-1 AF647 | BioLegend; 150003 | 1:400 |
| anti-CD19 APC-Cy7 | BD; 557655 | 1:200 |
| anti-IgA PE | eBioscience; 12-4204-82 | 1:100 |
| anti-CD93 biotin | eBioscience; 13-5892-85 | 1:50 |
| Streptavidin PE-CF594 | BD; 562284 | 1:2000 |
| anti-CD23 PE-Cy7 | BioLegend; 101614 | 1:100 |
| anti-CD43 BUV737 | BD; 612840 | 1:200 |

#### Intestine

Lamina propria cells from the small and large intestine were isolated as previously described (*100*).

#### Synovium

Knee synovium (patella and patellar tendon) and foot synovium (tibiotalar joint) were collected in RPMI. Synovia from two mice were pooled per sample. Tissue was digested for 30 min at 500 rpm with type VIII collagenase (0.75 mg/mL; Sigma; C2139) and dispase (1 mg/mL; Gibco; 17105041) in warm RPMI, passed through a 100 µm strainer and washed with cold 2% FBS in PBS. Cells were pelleted (393 × g, 7 min, 4 °C), resuspended in 0.5 ml 2% FBS/PBS and counted.

#### Spleen

Spleens were mashed through a 100 µm strainer in RPMI with GlutaMAX and 10% FBS and cells were pelleted (1,500 rpm, 5 min, 4 °C). Red blood cells were lysed with 3 ml ACK buffer (Lonza; 10548-E) for 3 min at RT and lysis was stopped with 30 ml PBS. Cells were passed through a 40 µm strainer, pelleted, resuspended in 10 ml PBS and counted.

#### Lung

Lungs were minced and digested for 30 min in a shaking water bath at 37 °C in RPMI (2 ml per lung) containing Liberase TM (Roche; R0366, 1:50) and DNase (Roche; R0066, 1:1000). Cells were passed through a 70 µm filter, pelleted (400 × g, 5 min, 4 °C) and treated with osmotic lysis buffer (2–3 ml, 3 min). Cells were washed, stained in 96-well plates and resuspended in 123count eBeads (Invitrogen; R2184, 1:10) to obtain absolute cell counts.

#### Peritoneal fluid

Peritoneal cells were collected by lavage with 10 ml PBS and processed as described for the spleen (ACK lysis, 40 µm strainer, counting).

### Statistics

Statistical analysis and graphing were performed in GraphPad Prism v9.2.0. Data were analyzed by unpaired t-test or analysis of variance (ANOVA) followed by Tukey’s multiple comparisons test, after confirming normality of residuals (QQ plots) and homogeneity of variance (residual plots). Data are presented as mean ± SEM, and p < 0.05 was considered statistically significant (ns, p > 0.05; *p ≤ 0.05; **p ≤ 0.01; ***p ≤ 0.001; ****p ≤ 0.0001).

#### Statistical analysis of non-normal data

A log-linear model as implemented in Genstat v24 (*101*), was fitted to the data. For Figure 4, the model included genotype, housing and their interaction term genotype.housing. For Figure 5, the model included genotype, IgA and their interaction term genotype.IgA. Observations under the detection limit were generated as a pseudo-random sample from the beta-distribution with parameters α= 1 and β=5, with the detection limit set as upper limit. Upon non-significance of the interaction term, the interaction was excluded from the fitted model. The variance dispersion parameter for responses was estimated. A Wald statistic for dropping terms from a regression was used to assess the significance of the interaction and the main effects.

#### Repeated measurements analysis of body weight and MEICS score data

Body weight and MEICS score data were analyzed as repeated measurements using the method of residual maximum likelihood (REML) as implemented in Genstat v24. Briefly, a linear mixed model (random terms underlined) of the following form: y = µ + genotype + time + genotype.time + id.time, was fitted to the data. Genotype, time and their two-way interaction term represent the fixed effects; the terms id.time represent the random term. Id.time is the residual error term with dependent errors because the repeated measurements are taken of the same individual id, causing correlations among observations. The correlation model AR1 (= autoregressive of order 1 with unequally spaced time points) was giving the best fitting model based on the Akaike’s information criterion coefficient. This model allows the correlations between measurements to decrease as the length of time between them increases. The significance of the fixed terms was assessed using an approximate F-test or a Wald-test as implemented in Genstat v24.

#### Forest plots T cell and myeloid cell panel

Immune cell counts were analysed separately for each organ using the conjugate hierarchical generalized linear model (HGLM) implemented in the R package hglm. Cell type, genotype and their interaction were included as fixed effects, with mouse ID as a random effect. Fixed effects were modelled using a Poisson distribution and random effects using a Gamma distribution, both with a log link. The response variance-dispersion parameter was estimated. Genotype effects on the log-transformed scale were evaluated using t-statistics and presented in forest plots on both the transformed and fold-change scales.

#### Forest plots B cell panel

Immune cell proportions were analysed separately for each organ and housing condition, using the conjugate hierarchical generalized linear model (HGLM) implemented in the R package hglm. Cell type, genotype and their interaction were included as fixed effects, with mouse ID as a random effect. Fixed effects were modelled using a Binomial distribution and random effects using a Beta distribution, both with a logit link. The response variance-dispersion parameter was estimated. Genotype effects on the logit-transformed scale were evaluated using t-statistics and presented in forest plots on both the transformed and odds ratio scales.

## Acknowledgements

We thank the Germ-Free and Gnotobiotic Mouse Facility Ghent (GFMF), Vanessa Andries for coordinating the germ-free and gnotobiotic mouse experiments and Korneel Barbry for animal care; Leander Vandereecken for animal care at the VIB-UGent Center for Inflammation Research; Gert Van Isterdael and Julie Van Duyse of the VIB Flow Core Ghent, Amanda Goncalves and Jérémy Verbeke of the VIB BioImaging Core Ghent and Kelly Lemaire for experimental support; and Marnik Vuylsteke for statistical consultancy. Research performed together with S.V. was supported by Ghent University (bof/baf/3y/2026/01/002). Research performed in the lab of B.N.L. was funded by an Excellence of Science (EOS) consortium grant (3G0H1222; BENEFICIARIES consortium), the Fund Alphonse and Jean Forton and the Belgian Cystic Fibrosis Association through the King Baudouin Foundation (2020-J1810150-218018; EXT/ONZ/000256), and the Scientific Research Fund (FWO) Flanders (G024423N). B.N.L. is supported by an ERC Advanced grant (789384) and ERC Proof of Concept grant (CRYSTAL-SHINE project 101189517), and by a Special Research Fund of Ghent University Methusalem grant (01M01521). B.N.L. is funded by the European Union. Views and opinions expressed are however those of the author(s) only and do not necessarily reflect those of the European Union or the European Research Council Executive Agency. Neither the European Union nor the granting authority can be held responsible for them. This work is supported by ERC grant ASTHMACRYSTALCLEAR(789384) and CRYSTAL-SHINE (101189517). Research in the Host-Microbiota-Interaction lab headed by L.V. is supported by FWO (G0APB25N, 1220226N, 3G0H2511, G0AE225N, S009626N, 1114626N), Foundation against Cancer (365V05625) and Ghent University (bof/baf/4y/2024/01/269, 01G03122).

## References

1. Knoop KA, Newberry RD, Goblet cells: multifaceted players in immunity at mucosal surfaces. Mucosal Immunology, doi: 10.1038/s41385-018-0039-y (2018).

2. Birchenough GMH, Nystrom EEL, Johansson MEV, Hansson GC, A sentinel goblet cell guards the colonic crypt by triggering Nlrp6-dependent Muc2 secretion. Science (1979), doi: 10.1126/science.aaf7419 (2016).

3. Nyström EEL, Martinez-Abad B, Arike L, Birchenough GMH, Nonnecke EB, Castillo PA, et al, An intercrypt subpopulation of goblet cells is essential for colonic mucus barrier function. Science (1979) 372 (2021).

4. J. R. McDole, L. W. Wheeler, K. G. McDonald, B. Wang, V. Konjufca, K. A. Knoop, R. D. Newberry, M. J. Miller, Goblet cells deliver luminal antigen to CD103+ dendritic cells in the small intestine. Nature 483, 345–349 (2012).

5. Wallaeys C, Garcia-Gonzalez N, Libert C, Paneth cells as the cornerstones of intestinal and organismal health: a primer. EMBO Mol Med 15 (2023).

6. K. E. Huus, C. Petersen, B. B. Finlay, Diversity and dynamism of IgA−microbiota interactions. Nature Reviews Immunology, doi: 10.1038/s41577-021-00506-1 (2021).

7. O. Pabst, A. Izcue, Secretory IgA: controlling the gut microbiota. Nat Rev Gastroenterol Hepatol 19 (2022).

8. O. Pabst, E. Slack, IgA and the intestinal microbiota: the importance of being specific. Mucosal Immunology, doi: 10.1038/s41385-019-0227-4 (2020).

9. Donaldson GP, Ladinsky MS, Yu KB, Sanders JG, Yoo BB, Chou WC, et al, Gut microbiota utilize immunoglobulin a for mucosal colonization. Science (1979) 360 (2018).

10. S. Van Der Post, K. S. Jabbar, G. Birchenough, L. Arike, N. Akhtar, H. Sjovall, Structural weakening of the colonic mucus barrier is an early event in ulcerative colitis pathogenesis. Gut, doi: 10.1136/gutjnl-2018-317571 (2019).

11. L. Erkert, L. M. Haag, C. Becker, Paneth cells as orchestrators of epithelial barrier defense and emerging therapeutic targets in inflammatory bowel disease. Semin Immunopathol 48 (2026).

12. G. Bamias, K. Kitsou, J. Rivera-Nieves, The Underappreciated Role of Secretory IgA in IBD. Inflammatory Bowel Diseases, doi: 10.1093/ibd/izad024 (2023).

13. S. A. Doumas, C. Tsironis, A. A. Bolaji, P. Garantziotis, E. Frangou, Glomerulonephritis and inflammatory bowel disease: A tale of gut-kidney axis dysfunction. Autoimmunity Reviews, doi: 10.1016/j.autrev.2023.103327 (2023).

14. M. Xiao, Y. Ran, J. Shao, Z. Lei, Y. Chen, Y. Li, Causal association between inflammatory bowel disease and IgA nephropathy: A bidirectional two-sample Mendelian randomization study. Front Genet 13 (2022).

15. Xu LL, Zhou XJ, Zhang H, An Update on the Genetics of IgA Nephropathy. Journal of Clinical Medicine, doi: 10.3390/jcm13010123 (2024).

16. Stamellou E, Seikrit C, Tang SCW, Boor P, Tesař V, Floege J, et al, IgA nephropathy. Nature Reviews Disease Primers 2023 9:1 9, 1–21 (2023).

17. Coppo R, Steps forward in the treatment of IgA nephropathy. Nat Rev Nephrol 21, 218–9 (2025).

18. C. Seikrit, O. Pabst, The immune landscape of IgA induction in the gut. Seminars in Immunopathology, doi: 10.1007/s00281-021-00879-4 (2021).

19. Seikrit C, Schimpf JI, Wied S, Stamellou E, Izcue A, Pabst O, et al, Intestinal permeability in patients with IgA nephropathy and other glomerular diseases: an observational study. J Nephrol 36 (2023).

20. K. Yao, L. Zheng, W. Chen, Y. Xie, C. Liao, T. Zhou, Characteristics, pathogenic and therapeutic role of gut microbiota in immunoglobulin A nephropathy. Front Immunol 16 (2025).

21. P. J. Gleeson, N. Benech, J. Chemouny, E. Metallinou, L. Berthelot, J. Da Silva, The gut microbiota posttranslationally modifies IgA1 in autoimmune glomerulonephritis. Sci Transl Med 16 (2024).

22. Y. Wang, L. Tian, L. Sun, W. Zhou, W. Zhi, J. Qing, Gut Microbes in Immunoglobulin A Nephropathy and Their Potential Therapeutic Applications. Frontiers in Medicine, doi: 10.3389/fmed.2022.823267 (2022).

23. X. Yuan, J. Qing, W. Zhi, F. Wu, Y. Yan, Y. Li, Gut and respiratory microbiota landscapes in IgA nephropathy: a cross-sectional study. Ren Fail 46 (2024).

24. Shimshek DR, Kim J, Hübner MR, Spergel DJ, Buchholz F, Casanova E, et al, Codon-improved Cre recombinase (iCre) expression in the mouse. Genesis 32, 19–26 (2002).

25. Hasnain SZ, Thornton DJ, Grencis RK, Changes in the mucosal barrier during acute and chronic Trichuris muris infection. Parasite Immunol 33, 45–55 (2011).

26. J. E. Klementowicz, M. A. Travis, R. K. Grencis, Trichuris muris: A model of gastrointestinal parasite infection. Semin Immunopathol 34, 815–828 (2012).

27. D. M. Oram, R. K. Holmes, “Diphtheria toxin” in The Comprehensive Sourcebook of Bacterial Protein Toxins (2006), pp. 245–256.

28. M. Saito, T. Iwawaki, C. Taya, H. Yonekawa, M. Noda, Y. Inui, Diphtheria toxin receptor–mediated conditional and targeted cell ablation in transgenic mice. Nature Biotechnology 19, 746–750 (2001).

29. Becker C, Fantini MC, Wirtz S, Nikolaev A, Kiesslich R, Lehr HA, et al, In vivo imaging of colitis and colon cancer development in mice using high resolution chromoendoscopy. Gut 54, 950–4 (2005).

30. Wei H, Wang JY, Role of Polymeric Immunoglobulin Receptor in IgA and IgM Transcytosis. Int J Mol Sci 22, 2284 (2021).

31. N. Garcia-Gonzalez, C. Wallaeys, W. Toussaint, T. Hochepied, S. Dewaele, S. De Beul, R. Van Loke, J. Vandewalle, S. Timmermans, R. S. Blumberg, C. Libert, Generation of a new Paneth cell–specific Cre-recombinase transgenic mouse line. Front. Immunol. 16, 1576995 (2025).

32. C. Mooser, F. Ronchi, J. P. Limenitakis, C. Kalbermatter, S. Christensen, M. G. De Agüero, T. Fuhrer, U. Sauer, A. J. Macpherson, S. C. Ganal-Vonarburg, Diet-derived LPS determines intestinal IgA induction and repertoire characteristics independently of the microbiota. Immunity 58, 1778–1793.e7 (2025).

33. J. J. Bunker, S. A. Erickson, T. M. Flynn, C. Henry, J. C. Koval, M. Meisel, B. Jabri, D. A. Antonopoulos, P. C. Wilson, A. Bendelac, Natural polyreactive IgA antibodies coat the intestinal microbiota. Science 358, eaan6619 (2017).

34. Joep Grootjans, Epithelial endoplasmic reticulum stress orchestrates a protective IgA response. Science 363, 993–998 (2019).

35. A. Kaser, A.-H. Lee, A. Franke, J. N. Glickman, S. Zeissig, H. Tilg, E. E. S. Nieuwenhuis, D. E. Higgins, S. Schreiber, L. H. Glimcher, R. S. Blumberg, XBP1 Links ER Stress to Intestinal Inflammation and Confers Genetic Risk for Human Inflammatory Bowel Disease. Cell 134, 743–756 (2008).

36. Y. Kinashi, K. Tanaka, S. Kimura, M. Hirota, S. Komiyama, T. Shindo, A. Hashiguchi, D. Takahashi, S. Shibata, S.-I. Karaki, H. Ohno, K. Hase, Intestinal epithelium dysfunctions cause IgA deposition in the kidney glomeruli of intestine-specific Ap1m2-deficient mice. eBioMedicine 106, 105256 (2024).

37. E. K. Grasset, A. Chorny, S. Casas-Recasens, C. Gutzeit, G. Bongers, I. Thomsen, L. Chen, Z. He, D. B. Matthews, M. A. Oropallo, P. Veeramreddy, M. Uzzan, A. Mortha, J. Carrillo, B. S. Reis, M. Ramanujam, J. Sintes, G. Magri, P. J. Maglione, C. Cunningham-Rundles, R. J. Bram, J. Faith, S. Mehandru, O. Pabst, A. Cerutti, Gut T cell–independent IgA responses to commensal bacteria require engagement of the TACI receptor on B cells. Sci. Immunol. 5, eaat7117 (2020).

38. T. Kano, H. Suzuki, Y. Makita, Y. Nihei, Y. Fukao, M. Nakayama, M. Lee, R. Aoki, K. Yamada, M. Muto, Y. Suzuki, Lessons from IgA Nephropathy Models. IJMS 25, 11484 (2024).

39. Johansson MEV, Hansson GC, Preservation of mucus in histological sections, immunostaining of mucins in fixed tissue, and localization of bacteria with FISH. Methods Mol Biol 842, 229–35 (2012).

40. DI Tommaso P, Chatzou M, Floden EW, Barja PP, Palumbo E, Notredame C, Nextflow enables reproducible computational workflows. Nat Biotechnol 35, 316–9 (2017).

41. Ewels PA, Peltzer A, Fillinger S, Patel H, Alneberg J, Wilm A, et al, The nf-core framework for community-curated bioinformatics pipelines. Nat Biotechnol 38, 276–8 (2020).

42. Harshil Patel, Jonathan Manning, Phil Ewels, Maxime U Garcia, Alexander Peltzer, Rickard Hammarén, Olga Botvinnik, Adam Talbot, nf-core bot, Friederike Hanssen, Gregor Sturm, EladH1, Matthias Zepper, Gary Burnett, Denis Moreno, Ezra Greenberg, Pranathi Vemuri, Mahesh Binzer-Panchal, silviamorins, Lorena Pantano, Robert Syme, Gavin Kelly, Lorenzo Sola, Felix Krueger, Matthias Hörtenhuber, James A. Fellows Yates, emmanuel-tan, Gabriel Lichtenstein, Edmund Miller, Jose Espinosa-Carrasco, nf-core/rnaseq: nf-core/rnaseq v3.27.0 - Vanadium Vulture, version 3.27.0, Zenodo (2026); 10.5281/ZENODO.1400710.

43. The Bioconda Team, B. Grüning, R. Dale, A. Sjödin, B. A. Chapman, J. Rowe, C. H. Tomkins-Tinch, R. Valieris, J. Köster, Bioconda: sustainable and comprehensive software distribution for the life sciences. Nat Methods 15, 475–476 (2018).

44. F. Da Veiga Leprevost, B. A. Grüning, S. Alves Aflitos, H. L. Röst, J. Uszkoreit, H. Barsnes, M. Vaudel, P. Moreno, L. Gatto, J. Weber, M. Bai, R. C. Jimenez, T. Sachsenberg, J. Pfeuffer, R. Vera Alvarez, J. Griss, A. I. Nesvizhskii, Y. Perez-Riverol, BioContainers: an open-source and community-driven framework for software standardization. Bioinformatics 33, 2580–2582 (2017).

45. R Core Team, R: A language and environment for statistical computing, R Foundation for Statistical Computing (2026); https://www.R-project.org/.

46. C. Soneson, M. I. Love, M. D. Robinson, Differential analyses for RNA-seq: transcript-level estimates improve gene-level inferences. F1000Res 4, 1521 (2015).

47. M. I. Love, W. Huber, S. Anders, Moderated estimation of fold change and dispersion for RNA-seq data with DESeq2. Genome Biol 15, 550 (2014).

48. S. Xu, E. Hu, Y. Cai, Z. Xie, X. Luo, L. Zhan, W. Tang, Q. Wang, B. Liu, R. Wang, W. Xie, T. Wu, L. Xie, G. Yu, Using clusterProfiler to characterize multiomics data. Nat Protoc 19, 3292–3320 (2024).

49. G. Korotkevich, V. Sukhov, N. Budin, B. Shpak, M. N. Artyomov, A. Sergushichev, Fast gene set enrichment analysis. [Preprint] (2016). 10.1101/060012.

50. I. Dolgalev, msigdbr: MSigDB Gene Sets for Multiple Organisms in a Tidy Data Format, (2018); 10.32614/CRAN.package.msigdbr.

51. A. Subramanian, P. Tamayo, V. K. Mootha, S. Mukherjee, B. L. Ebert, M. A. Gillette, A. Paulovich, S. L. Pomeroy, T. R. Golub, E. S. Lander, J. P. Mesirov, Gene set enrichment analysis: A knowledge-based approach for interpreting genome-wide expression profiles. Proc. Natl. Acad. Sci. U.S.A. 102, 15545–15550 (2005).

52. A. Liberzon, A. Subramanian, R. Pinchback, H. Thorvaldsdóttir, P. Tamayo, J. P. Mesirov, Molecular signatures database (MSigDB) 3.0. Bioinformatics 27, 1739–1740 (2011).

53. A. Liberzon, C. Birger, H. Thorvaldsdóttir, M. Ghandi, J. P. Mesirov, P. Tamayo, The Molecular Signatures Database Hallmark Gene Set Collection. Cell Systems 1, 417–425 (2015).

54. A. S. Castanza, J. M. Recla, D. Eby, H. Thorvaldsdóttir, C. J. Bult, J. P. Mesirov, Extending support for mouse data in the Molecular Signatures Database (MSigDB). Nat Methods 20, 1619–1620 (2023).

55. S. Kolmykov, I. Yevshin, M. Kulyashov, R. Sharipov, Y. Kondrakhin, V. J. Makeev, I. V. Kulakovskiy, A. Kel, F. Kolpakov, GTRD: an integrated view of transcription regulation. Nucleic Acids Research 49, D104–D111 (2021).

56. M. Stephens, P. Carbonetto, D. Gerard, M. Lu, L. Sun, J. Willwerscheid, N. Xiao, ashr: Methods for Adaptive Shrinkage, using Empirical Bayes, (2016); 10.32614/CRAN.package.ashr.

57. Broad Institute and UC San Diego, GSEA User Guide. https://docs.gsea-msigdb.org/#GSEA/GSEA_User_Guide/.

58. P. Badia-i-Mompel, J. Vélez Santiago, J. Braunger, C. Geiss, D. Dimitrov, S. Müller-Dott, P. Taus, A. Dugourd, C. H. Holland, R. O. Ramirez Flores, J. Saez-Rodriguez, decoupleR: ensemble of computational methods to infer biological activities from omics data. Bioinformatics Advances 2, vbac016 (2022).

59. S. Müller-Dott, E. Tsirvouli, M. Vazquez, R. O. Ramirez Flores, P. Badia-i-Mompel, R. Fallegger, D. Türei, A. Lægreid, J. Saez-Rodriguez, Expanding the coverage of regulons from high-confidence prior knowledge for accurate estimation of transcription factor activities. Nucleic Acids Research 51, 10934–10949 (2023).

60. R. Browaeys, W. Saelens, Y. Saeys, NicheNet: modeling intercellular communication by linking ligands to target genes. Nat Methods 17, 159–162 (2020).

61. R. Browaeys, NicheNet-v2: final networks and ligand-target matrices, Zenodo (2022); 10.5281/ZENODO.7074290.

62. Browaeys R, Gilis J, Sang-Aram C, De Bleser P, Hoste L, Tavernier S, et al, MultiNicheNet: a flexible framework for differential cell-cell communication analysis from multi-sample multi-condition single-cell transcriptomics data. bioRxiv 2023 (2023).

63. H. Wickham, Ggplot2 (Springer International Publishing, Cham, 2016; http://link.springer.com/10.1007/978-3-319-24277-4)*Use R!*

64. Zheng GXY, Terry JM, Belgrader P, Ryvkin P, Bent ZW, Wilson R, et al, Massively parallel digital transcriptional profiling of single cells. Nature Communications 2017 8:1 8, 1–12 (2017).

65. GENCODE - Mouse Release M24 (2024).

66. Hao Y, Stuart T, Kowalski MH, Choudhary S, Hoffman P, Hartman A, et al, Dictionary learning for integrative, multimodal and scalable single-cell analysis. Nature Biotechnology 2023 42:2 42, 293–304 (2023).

67. Andreatta M, Carmona SJ, UCell: Robust and scalable single-cell gene signature scoring. Comput Struct Biotechnol J 19, 3796–8 (2021).

68. Daniel Straub, Daniel Lundin, Jeanette Tångrot, Alexander Peltzer, nf-core bot, Matthew, Pieter Provoost, emnilsson, Sateesh_Peri, Adam Bennett, Tom Brekke, 4less, John Sundh, DiegoBrambilla, Sofoklis Keisaris, Lokeshwaran Manoharan, Till Englert, Asaf Peer, Aaron O’Brien, Maxime U Garcia, Nicolas Henry, Thomas Weber, Chandini, Matthias Hörtenhuber, Sam Minot, Thom Griffioen, Zack Mudge, Dan Clayton, Gisela Gabernet, Venkat Malladi, nf-core/ampliseq: Ampliseq Version 3.1.0, version 3.1.0, Zenodo (2026); 10.5281/ZENODO.1493841.

69. Straub D, Blackwell N, Langarica-Fuentes A, Peltzer A, Nahnsen S, Kleindienst S, Interpretations of Environmental Microbial Community Studies Are Biased by the Selected 16S rRNA (Gene) Amplicon Sequencing Pipeline. Front Microbiol 11 (2020).

70. Andrews, S., FastQC: a quality control tool for high throughput sequence data, version Version: 0.12.1, Babraham Bioinformatics (2023); https://www.bioinformatics.babraham.ac.uk/projects/fastqc/.

71. P. Ewels, M. Magnusson, S. Lundin, M. Käller, MultiQC: summarize analysis results for multiple tools and samples in a single report. Bioinformatics 32, 3047–3048 (2016).

72. M. Martin, Cutadapt removes adapter sequences from high-throughput sequencing reads. EMBnet j. 17, 10 (2011).

73. B. J. Callahan, P. J. McMurdie, M. J. Rosen, A. W. Han, A. J. A. Johnson, S. P. Holmes, DADA2: High-resolution sample inference from Illumina amplicon data. Nat Methods 13, 581–583 (2016).

74. B. J. Callahan, J. Wong, C. Heiner, S. Oh, C. M. Theriot, A. S. Gulati, S. K. McGill, M. K. Dougherty, High-throughput amplicon sequencing of the full-length 16S rRNA gene with single-nucleotide resolution. Nucleic Acids Research 47, e103–e103 (2019).

75. C. Quast, E. Pruesse, P. Yilmaz, J. Gerken, T. Schweer, P. Yarza, J. Peplies, F. O. Glöckner, The SILVA ribosomal RNA gene database project: improved data processing and web-based tools. Nucleic Acids Research 41, D590–D596 (2012).

76. B. Callahan, Silva taxonomic training data formatted for DADA2 (Silva version 138.2), Zenodo (2024); 10.5281/ZENODO.14169026.

77. P. J. McMurdie, S. Holmes, phyloseq: An R Package for Reproducible Interactive Analysis and Graphics of Microbiome Census Data. PLoS ONE 8, e61217 (2013).

78. R. Huang, C. Soneson, F. G. M. Ernst, K. C. Rue-Albrecht, G. Yu, S. C. Hicks, M. D. Robinson, TreeSummarizedExperiment: a S4 class for data with hierarchical structure. F1000Res 9, 1246 (2021).

79. T. Borman, L. Lahti, S. Shetty, F. M. Ernst, Orchestrating Microbiome Analysis with Bioconductor (2026; https://bioconductor.org/books/release/OMA/).

80. T. Borman, F. G. M. Ernst, L. Lahti, miaViz: Microbiome Analysis Plotting and Visualization, version 1.20.0 (2026); https://bioconductor.org/packages/miaViz.

81. D. J. McCarthy, K. R. Campbell, A. T. L. Lun, Q. F. Wills, Scater: pre-processing, quality control, normalization and visualization of single-cell RNA-seq data in R. Bioinformatics 33, 1179–1186 (2017).

82. C. Ahlmann-Eltze, I. Patil, ggsignif: Significance Brackets for “ggplot2,” (2017); 10.32614/CRAN.package.ggsignif.

83. G. B. Gloor, J. M. Macklaim, V. Pawlowsky-Glahn, J. J. Egozcue, Microbiome Datasets Are Compositional: And This Is Not Optional. Front. Microbiol. 8, 2224 (2017).

84. J. Oksanen, G. L. Simpson, F. G. Blanchet, R. Kindt, P. Legendre, P. R. Minchin, R. B. O’Hara, P. Solymos, M. H. H. Stevens, E. Szoecs, H. Wagner, M. Bedward, B. Bolker, D. Borcard, G. Carvalho, M. De Caceres, S. Durand, H. B. A. Evangelista, G. Hannigan, M. O. Hill, L. Lahti, C. Martino, M.-H. Ouellette, E. Ribeiro Cunha, T. Smith, A. Stier, C. J. F. Ter Braak, J. Weedon, vegan: Community Ecology Package, (2001); 10.32614/CRAN.package.vegan.

85. M. J. Anderson, A new method for non-parametric multivariate analysis of variance. Austral Ecology 26, 32–46 (2001).

86. H. Mallick, A. Rahnavard, L. J. McIver, S. Ma, Y. Zhang, L. H. Nguyen, T. L. Tickle, G. Weingart, B. Ren, E. H. Schwager, S. Chatterjee, K. N. Thompson, J. E. Wilkinson, A. Subramanian, Y. Lu, L. Waldron, J. N. Paulson, E. A. Franzosa, H. C. Bravo, C. Huttenhower, Multivariable association discovery in population-scale meta-omics studies. PLoS Comput Biol 17, e1009442 (2021).

87. Y. Benjamini, Y. Hochberg, Controlling the False Discovery Rate: A Practical and Powerful Approach to Multiple Testing. Journal of the Royal Statistical Society Series B: Statistical Methodology 57, 289–300 (1995).

88. A. Steimle, M. Neumann, E. T. Grant, S. Willieme, A. De Sciscio, A. Parrish, M. Ollert, E. Miyauchi, T. Soga, S. Fukuda, H. Ohno, M. S. Desai, Gut microbial factors predict disease severity in a mouse model of multiple sclerosis. Nat Microbiol 9, 2244–2261 (2024).

89. S. Stamouli, M. E. Beber, T. Normark, T. A. Christensen, L. Andersson-Li, M. Borry, M. Jamy, nf-core community, J. A. Fellows Yates, nf-core/taxprofiler: highly parallelised and flexible pipeline for metagenomic taxonomic classification and profiling. [Preprint] (2023). 10.1101/2023.10.20.563221.

90. S. Chen, Y. Zhou, Y. Chen, J. Gu, fastp: an ultra-fast all-in-one FASTQ preprocessor. Bioinformatics 34, i884–i890 (2018).

91. B. Bushnell, BBMap; http://sourceforge.net/projects/bbmap/.

92. B. Langmead, S. L. Salzberg, Fast gapped-read alignment with Bowtie 2. Nat Methods 9, 357–359 (2012).

93. P. Danecek, J. K. Bonfield, J. Liddle, J. Marshall, V. Ohan, M. O. Pollard, A. Whitwham, T. Keane, S. A. McCarthy, R. M. Davies, H. Li, Twelve years of SAMtools and BCFtools. GigaScience 10, giab008 (2021).

94. D. E. Wood, J. Lu, B. Langmead, Improved metagenomic analysis with Kraken 2. Genome Biol 20, 257 (2019).

95. B. Langmead, Kraken 2/Bracken refseq indexes; https://benlangmead.github.io/aws-indexes/k2.

96. J. Lu, F. P. Breitwieser, P. Thielen, S. L. Salzberg, Bracken: estimating species abundance in metagenomics data. PeerJ Computer Science 3, e104 (2017).

97. M. E. Beber, M. Borry, S. Stamouli, J. A. F. Yates, TAXPASTA: TAXonomic Profile Aggregation andSTAndardisation. JOSS 8, 5627 (2023).

98. C. L. Schoch, S. Ciufo, M. Domrachev, C. L. Hotton, S. Kannan, R. Khovanskaya, D. Leipe, R. Mcveigh, K. O’Neill, B. Robbertse, S. Sharma, V. Soussov, J. P. Sullivan, L. Sun, S. Turner, I. Karsch-Mizrachi, NCBI Taxonomy: a comprehensive update on curation, resources and tools. Database 2020, baaa062 (2020).

99. S. Prakash, N. J. Steers, Y. Li, E. Sanchez-Rodriguez, M. Verbitsky, I. Robbins, J. Simpson, S. Pathak, M. Raska, C. Reily, A. Ng, J. Liang, N. DeMaria, A. Katiraei, K. O. Stevens, C. Fischman, S. Shapiro, S. Kodali, J. McCutchan, H. Park, D. Eliby, M. Delsante, L. Allegri, E. Fiaccadori, M. Bodria, M. Marasa, E. Raveche, B. A. Julian, A.-C. Uhlemann, K. Kiryluk, H. Zhang, V. D. D’Agati, S. Sanna-Cherchi, J. Novak, A. G. Gharavi, Loss of GalNAc-T14 links O-glycosylation defects to alterations in B cell homing in IgA nephropathy. Journal of Clinical Investigation 135, e181164 (2025).

100. Bain CC, Mowat AMI, CD200 receptor and macrophage function in the intestine. Immunobiology 217, 643–51 (2012).

101. VSN International, Genstat, version 24th Edition, VSN International (2022); https://vsni.co.uk/software/genstat.

