## Supplementary data for "Intestinal goblet cell loss induces microbiota-independent mucosal and systemic IgA dysregulation"

#### Supplementary Figures

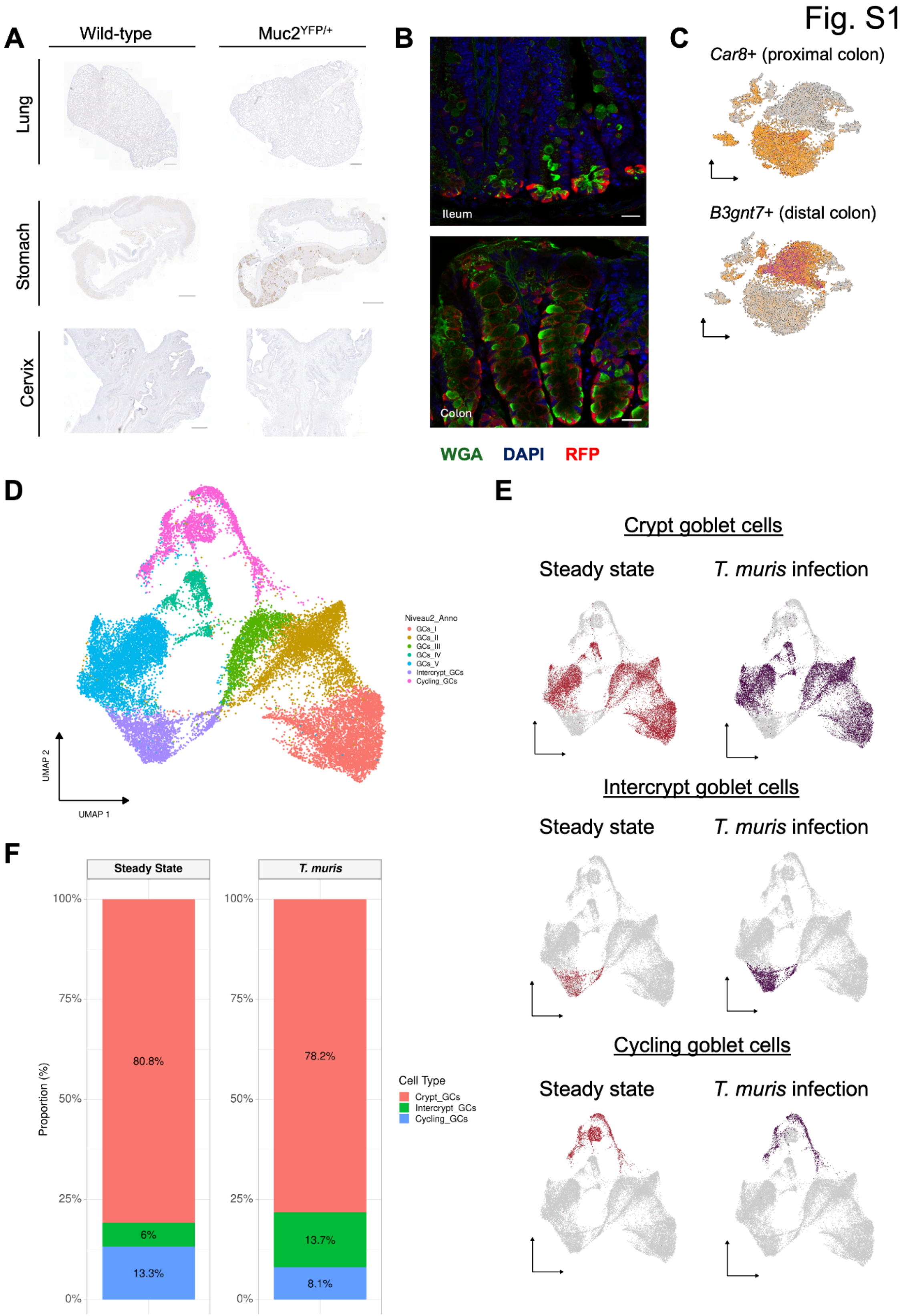

**Figure S1: Intestinal reporter expression of the Muc2iCre line and colonic goblet cell subpopulations during Trichuris muris infection** (A) Anti-GFP immunohistochemistry on sections of lung, stomach and cervix from Muc2$^{YFP/+}$ mice. Scale bars, 500 µm (lung, cervix) and 1000 µm (stomach). (B) Immunofluorescence staining for RFP (red) on sections of ileum and colon from Muc2$^{RFP/+}$ mice. Wheat germ agglutinin (WGA, green) and DAPI (blue) were used as counterstains. Scale bars, 20 µm. (C) Feature plots of *Car8* (proximal colon marker) and *B3gnt7* (distal colon marker) in sorted RFP+ colonic cells from Muc2$^{RFP/+}$ mice, showing the underlying proximal-distal clustering. (D) UMAP of scRNA-seq data from colonic samples at steady state (n = 1) and after Trichuris muris infection (n = 1), merged. Unsupervised Louvain clustering identified three GC subpopulations: crypt, intercrypt and cycling GCs. (E) UMAP plots of scRNA-seq data showing changes in GC subpopulations at steady state and after Trichuris muris infection (steady state n = 1, infected n = 1). (F) Relative abundance of GC subtypes at steady state and after T. muris infection (steady state n = 1, infected n = 1).

Fig. S2

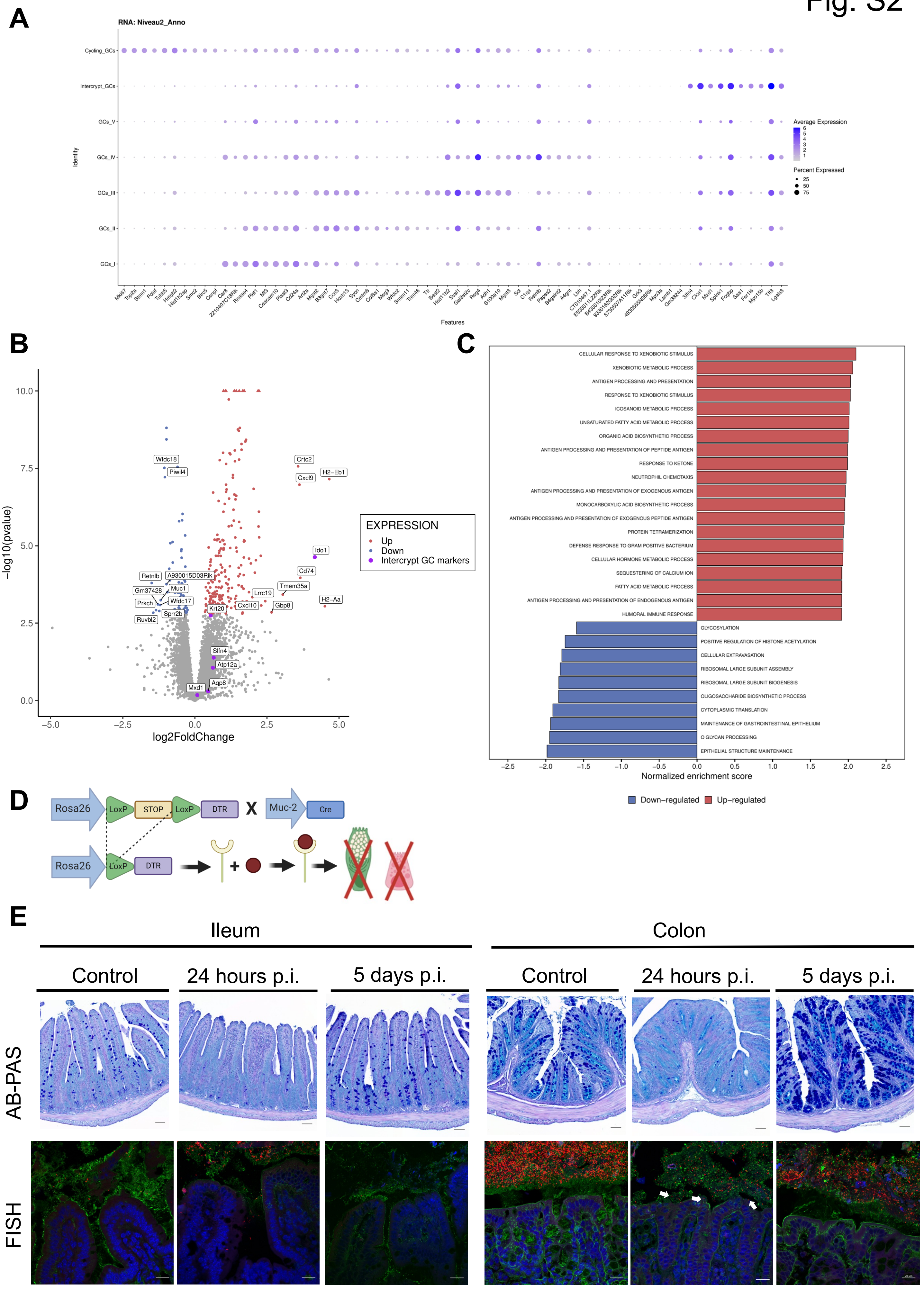
A
RNA: Niveau2_Anno
Cycling_GCs
Intercrypt_GCs
GCs_V
GCs_IV
GCs_III
GCs_II
GCs_I
Identity
Features
Average Expression
Percent Expressed
25
50
75
B
−log10(pvalue)
log2FoldChange
EXPRESSION
Up
Down
Intercrypt GC markers
Wfdc18
Piwil4
Crtc2
H2−Eb1
Cxcl9
Ido1
Cd74
Tmem35a
H2−Aa
Retnlb
A930015D03Rik
Gm37428
Muc1
Prkch
Wfdc17
Sprr2b
Ruvbl2
Krt20
Lrrc19
Cxcl10
Gbp8
Slfn4
Atp12a
Aqp8
Mxd1
C
CELLULAR RESPONSE TO XENOBIOTIC STIMULUS
XENOBIOTIC METABOLIC PROCESS
ANTIGEN PROCESSING AND PRESENTATION
RESPONSE TO XENOBIOTIC STIMULUS
ICOSANOID METABOLIC PROCESS
UNSATURATED FATTY ACID METABOLIC PROCESS
ORGANIC ACID BIOSYNTHETIC PROCESS
ANTIGEN PROCESSING AND PRESENTATION OF PEPTIDE ANTIGEN
RESPONSE TO KETONE
NEUTROPHIL CHEMOTAXIS
ANTIGEN PROCESSING AND PRESENTATION OF EXOGENOUS ANTIGEN
MONOCARBOXYLIC ACID BIOSYNTHETIC PROCESS
ANTIGEN PROCESSING AND PRESENTATION OF EXOGENOUS PEPTIDE ANTIGEN
PROTEIN TETRAMERIZATION
DEFENSE RESPONSE TO GRAM POSITIVE BACTERIUM
CELLULAR HORMONE METABOLIC PROCESS
SEQUESTERING OF CALCIUM ION
FATTY ACID METABOLIC PROCESS
ANTIGEN PROCESSING AND PRESENTATION OF ENDOGENOUS ANTIGEN
HUMORAL IMMUNE RESPONSE
GLYCOSYLATION
POSITIVE REGULATION OF HISTONE ACETYLATION
CELLULAR EXTRAVASATION
RIBOSOMAL LARGE SUBUNIT ASSEMBLY
RIBOSOMAL LARGE SUBUNIT BIOGENESIS
OLIGOSACCHARIDE BIOSYNTHETIC PROCESS
CYTOPLASMIC TRANSLATION
MAINTENANCE OF GASTROINTESTINAL EPITHELIUM
O GLYCAN PROCESSING
EPITHELIAL STRUCTURE MAINTENANCE
Normalized enrichment score
Down−regulated
Up−regulated
D
Rosa26
LoxP
STOP
LoxP
DTR
X
Muc-2
Cre
Rosa26
LoxP
DTR
E
Ileum
Colon
Control
24 hours p.i.
5 days p.i.
Control
24 hours p.i.
5 days p.i.
AB-PAS
FISH

**Figure S2: Goblet cell transcriptional responses to Trichuris muris infection and bacterial adherence after acute goblet and Paneth cell depletion** (A) Dot plot of selected GC marker genes across the crypt, intercrypt and cycling GC subpopulations. Dot size indicates the proportion of cells expressing the gene; color intensity indicates average expression level. (B) Volcano plot of differentially expressed genes in YFP+ GCs from *T. muris*-infected versus uninfected mice. The top 10 upregulated (red) and top 10 downregulated (blue) genes are highlighted; marker genes of the intercrypt GC subpopulation are shown in purple (steady state n = 3, infected n = 5). (C) Gene set enrichment analysis (GSEA) of YFP+ GCs from T. muris-infected versus uninfected mice, showing the top enriched biological processes (steady state n = 3, infected n = 5). (D) Schematic of the Muc2$^{DTR}$ depletion model. Cre-LoxP recombination drives expression of the diphtheria toxin receptor (DTR) in PCs and GCs, resulting in cell death upon diphtheria toxin (DT) injection. (E) Ileal and colonic sections of Muc2$^{DTR}$ mice at steady state and 24 h and 5 days after DT injection, stained with Alcian blue-periodic acid-Schiff (AB-PAS; upper panels) or by fluorescence in situ hybridization with the pan-bacterial probe EUB338 (red; lower panels), with DAPI (blue) and UEA1 and WGA (green). White arrows indicate bacteria adhering to the epithelium. Scale bars, 50 µm (AB-PAS) and 20 µm (FISH).

Fig. S3

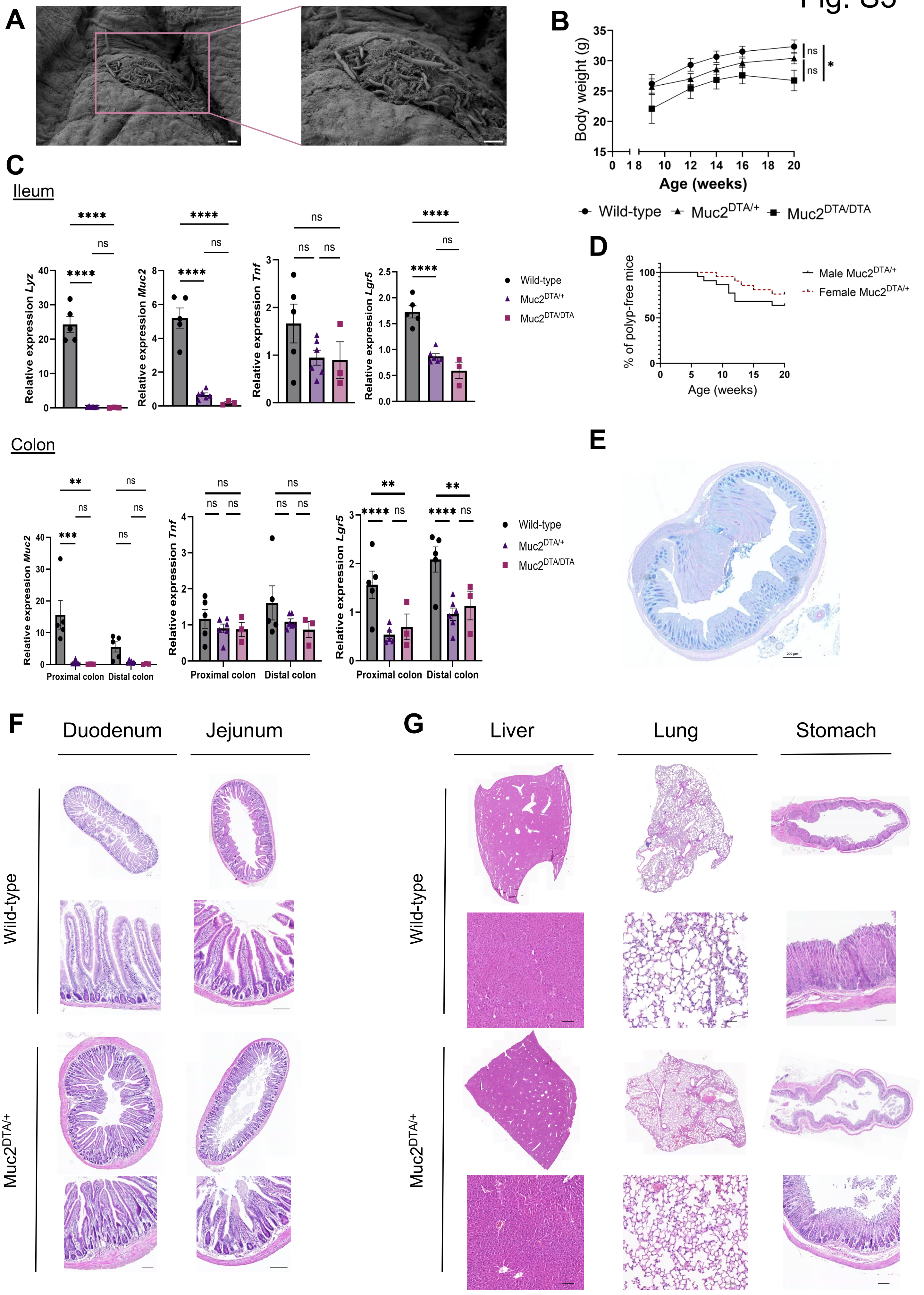
A
B
Body weight (g)
Age (weeks)
ns
*
Wild-type
Muc2DTA/+
Muc2DTA/DTA
C
Ileum
Relative expression Lyz
Relative expression Muc2
Relative expression Tnf
Relative expression Lgr5
****
***
**
ns
Colon
Proximal colon
Distal colon
D
% of polyp-free mice
Age (weeks)
Male Muc2DTA/+
Female Muc2DTA/+
E
200 µm
F
Duodenum
Jejunum
G
Liver
Lung
Stomach

**Figure S3: Phenotyping of goblet and Paneth cell-depleted Muc2$^{DTA/+}$ mice** (A) Scanning electron microscopy (SEM) images of the colonic surface epithelium of Muc2$^{DTA/+}$ mice showing bacterial biofilm. Scale bars, 2 µm. (B) Body weight of male WT (n = 5), Muc2$^{DTA/+}$ (n = 8) and Muc2$^{DTA/DTA}$ (n = 3) mice over time. REML model: main effects of time (###p < 0.001) and genotype (*p = 0.026); time × genotype interaction, ns. (C) RT-qPCR analysis of *Muc2*, *Lyz*, *Lgr5* and *Tnf* expression in ileal and colonic tissue of WT (n = 5), Muc2$^{DTA/+}$ (n = 6) and Muc2$^{DTA/DTA}$ (n = 3) mice. Ordinary one-way ANOVA with Tukey's multiple comparisons test. (D) Percentage of male and female Muc2$^{DTA/+}$ mice with polyps at different ages, as detected by colonoscopy. (E) AB-PAS-stained overview section of a colonic polyp from an 20-week-old Muc2$^{DTA/+}$ mouse. Scale bar, 200 µm. (F) Representative H&E-stained sections of duodenum and jejunum from WT and Muc2$^{DTA/+}$ mice. Scale bars, 100 µm. (G) Representative H&E-stained sections of lung, liver and stomach from WT and Muc2$^{DTA/+}$ mice. Scale bars, 100 µm (liver, stomach) and 50 µm (lung). Data are mean ± SEM and n indicates individual mice.

Fig. S4

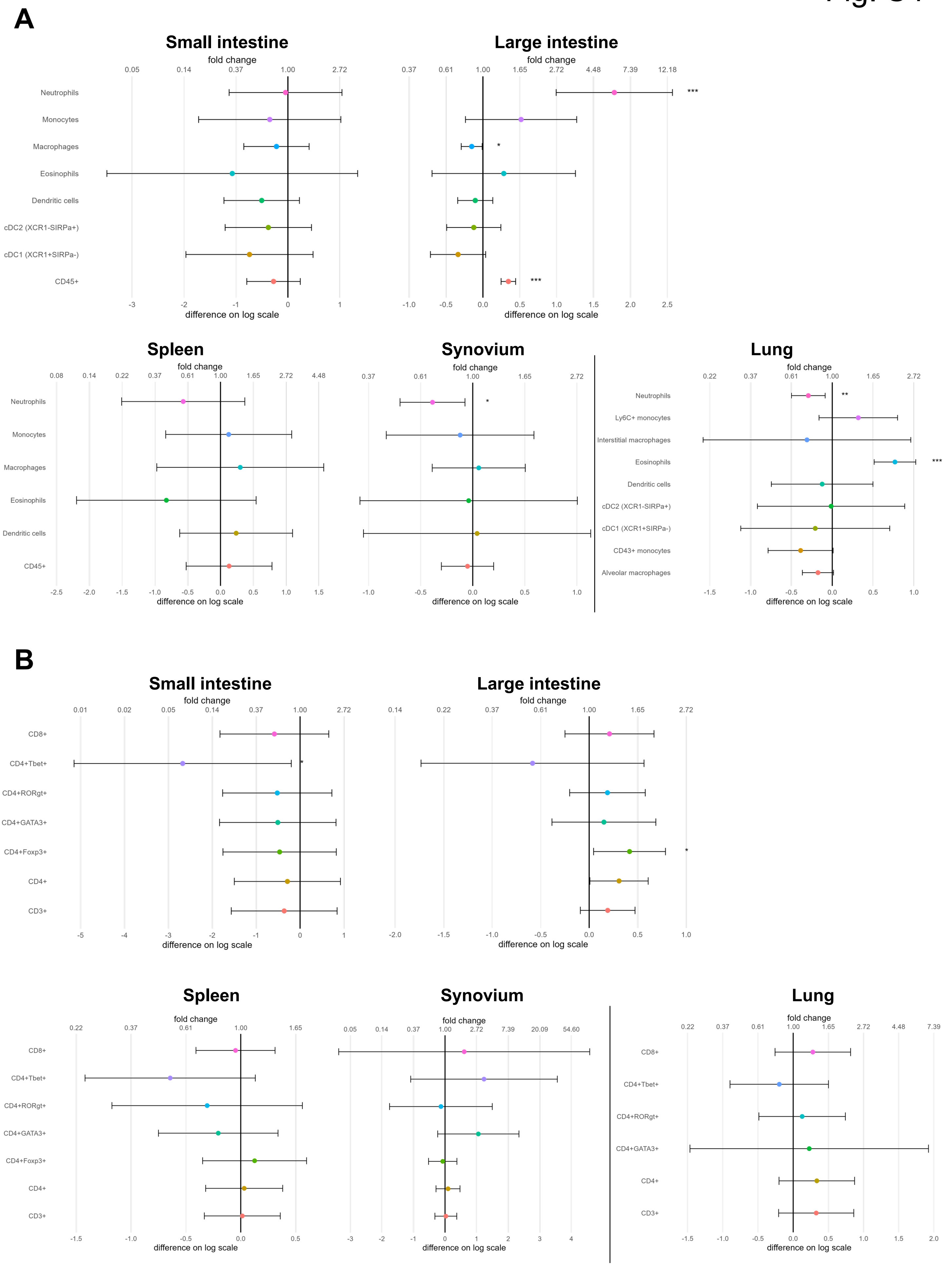
A
Small intestine
fold change
0.05 0.14 0.37 1.00 2.72
Neutrophils
Monocytes
Macrophages
Eosinophils
Dendritic cells
cDC2 (XCR1-SIRPa+)
cDC1 (XCR1+SIRPa-)
CD45+
-3 -2 -1 0 1
difference on log scale
Large intestine
fold change
0.37 0.61 1.00 1.65 2.72 4.48 7.39 12.18
***
*
***
-1.0 -0.5 0.0 0.5 1.0 1.5 2.0 2.5
difference on log scale
Spleen
fold change
0.08 0.14 0.22 0.37 0.61 1.00 1.65 2.72 4.48
Neutrophils
Monocytes
Macrophages
Eosinophils
Dendritic cells
CD45+
-2.5 -2.0 -1.5 -1.0 -0.5 0.0 0.5 1.0 1.5
difference on log scale
Synovium
fold change
0.37 0.61 1.00 1.65 2.72
*
-1.0 -0.5 0.0 0.5 1.0
difference on log scale
Lung
fold change
0.22 0.37 0.61 1.00 1.65 2.72
Neutrophils
Ly6C+ monocytes
Interstitial macrophages
Eosinophils
Dendritic cells
cDC2 (XCR1-SIRPa+)
cDC1 (XCR1+SIRPa-)
CD43+ monocytes
Alveolar macrophages
**
***
-1.5 -1.0 -0.5 0.0 0.5 1.0
difference on log scale
B
Small intestine
fold change
0.01 0.02 0.05 0.14 0.37 1.00 2.72
CD8+
CD4+Tbet+
CD4+RORgt+
CD4+GATA3+
CD4+Foxp3+
CD4+
CD3+
*
-5 -4 -3 -2 -1 0 1
difference on log scale
Large intestine
fold change
0.14 0.22 0.37 0.61 1.00 1.65 2.72
*
-2.0 -1.5 -1.0 -0.5 0.0 0.5 1.0
difference on log scale
Spleen
fold change
0.22 0.37 0.61 1.00 1.65
CD8+
CD4+Tbet+
CD4+RORgt+
CD4+GATA3+
CD4+Foxp3+
CD4+
CD3+
-1.5 -1.0 -0.5 0.0 0.5
difference on log scale
Synovium
fold change
0.05 0.14 0.37 1.00 2.72 7.39 20.09 54.60
-3 -2 -1 0 1 2 3 4
difference on log scale
Lung
fold change
0.22 0.37 0.61 1.00 1.65 2.72 4.48 7.39
CD8+
CD4+Tbet+
CD4+RORgt+
CD4+GATA3+
CD4+
CD3+
-1.5 -1.0 -0.5 0.0 0.5 1.0 1.5 2.0
difference on log scale

**Figure S4: Immune profiling of intestinal and extra-intestinal tissues in Muc2$^{DTA/+}$ mice** (A) Forest plots of myeloid cell populations, analyzed by flow cytometry, in small intestine, large intestine, synovial fluid, spleen and lung of 15-week-old Muc2$^{DTA/+}$ mice relative to WT mice (n = 5 per genotype). Points show the estimated difference of Muc2$^{DTA/+}$ relative to WT on the log scale (fold change on the top axis), with 95% confidence intervals, estimated for each cell type using a hierarchical generalized linear model (Poisson distribution with log link for fixed effects, Gamma-distributed random effect of mouse ID). Genotype effects were evaluated using t-statistics. (B) Forest plots of T cell populations, analyzed by flow cytometry, in the same tissues of 15-week-old Muc2$^{DTA/+}$ mice relative to WT mice (n = 5 per genotype). Plots are as in (A). HGLM; R. *$p < 0.05$, **$p < 0.01$.

Fig. S5

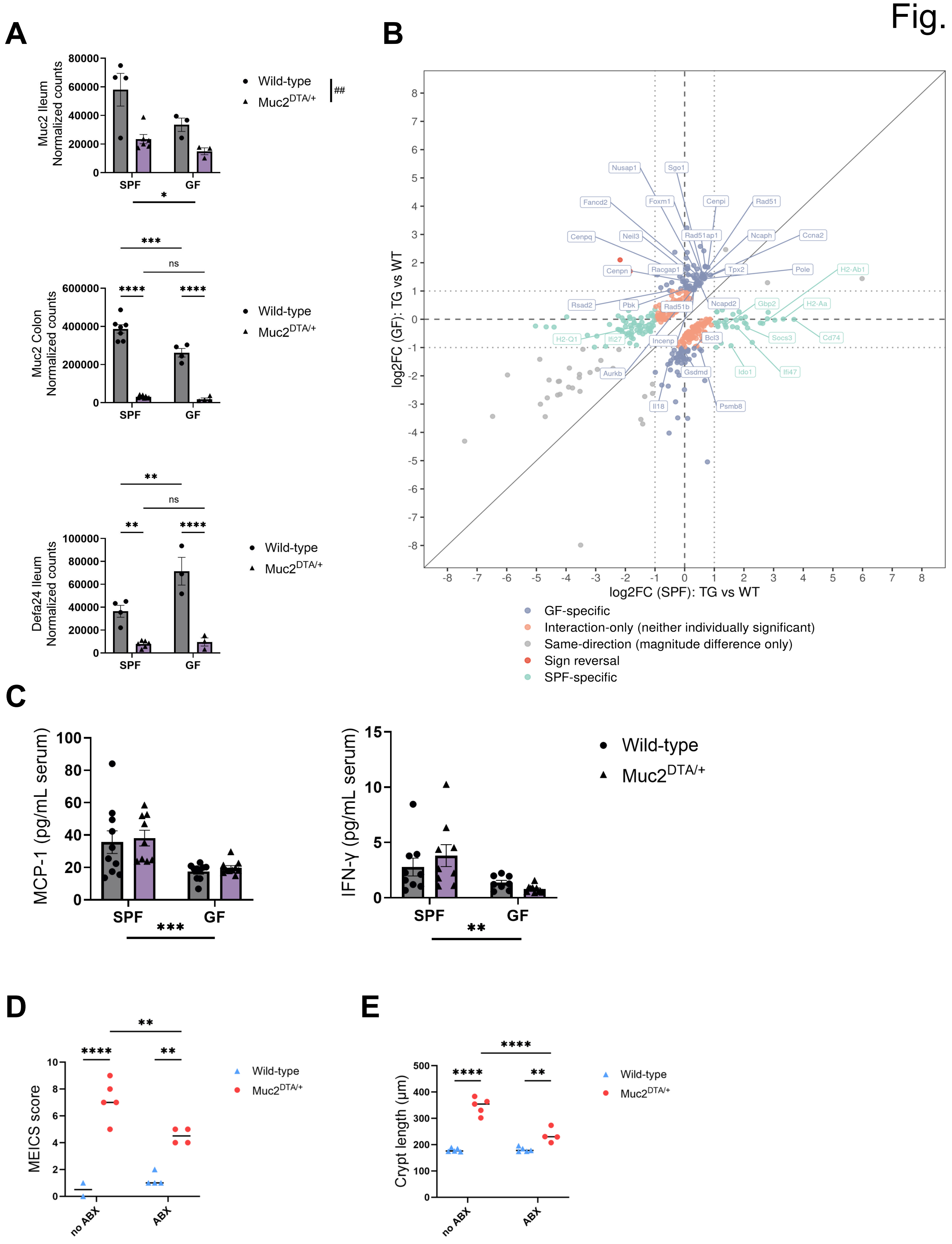
A
Muc2 Ileum Normalized counts
SPF
GF
Wild-type
Muc2DTA/+
##
Muc2 Colon Normalized counts
Defa24 Ileum Normalized counts
ns
B
log2FC (GF): TG vs WT
log2FC (SPF): TG vs WT
Nusap1
Sgo1
Fancd2
Foxm1
Cenpi
Rad51
Cenpq
Neil3
Rad51ap1
Ncaph
Ccna2
Cenpn
Racgap1
Tpx2
Pole
H2-Ab1
Rsad2
Pbk
Rad51b
Ncapd2
Gbp2
H2-Aa
H2-Q1
Ifi27
Incenp
Bcl3
Socs3
Cd74
Aurkb
Gsdmd
Ido1
Ifi47
Il18
Psmb8
GF-specific
Interaction-only (neither individually significant)
Same-direction (magnitude difference only)
Sign reversal
SPF-specific
C
MCP-1 (pg/mL serum)
IFN-γ (pg/mL serum)
D
MEICS score
no ABX
ABX
E
Crypt length (µm)

**Figure S5: Antibiotic treatment alleviates colonic pathology in Muc2$^{DTA/+}$ mice and genotype effects on the colonic epithelial transcriptome differ between SPF and GF conditions** (A) Normalized *Muc2* and *Defa24* transcript counts from bulk RNA sequencing of colonic and ileal tissue from WT and Muc2$^{DTA/+}$ mice kept under SPF or GF conditions. Two-way ANOVA. Muc2, ileum: main effect of housing (*p = 0.032) and genotype (##p < 0.01), interaction ns. Muc2, colon: genotype × housing with Tukey's multiple comparisons test (interaction, p = 0.03). Defa24, ileum: genotype × housing with Tukey's multiple comparisons test (interaction, p = 0.011). (B) Concordance of genotype effects between SPF and GF colonic epithelial cells. Scatter plot of log2 fold change (Muc2$^{DTA/+}$ versus WT) from bulk RNA sequencing under SPF (x-axis) and GF (y-axis) conditions. Shown are the 427 genes with a significant genotype × microbiota interaction (|interaction log2FC| ≥ 0.5, adjusted p ≤ 0.05). Genes are categorized as GF-specific (purple), SPF-specific (green), sign-reversed between conditions (red), same direction with different effect sizes (grey) or interaction-only, without a significant genotype effect in either condition alone (orange). Dashed lines mark log2FC = 0; dotted lines mark log2FC = ±1. (C) Bio-Plex measurement of MCP-1 and IFN-γ in serum of SPF and GF WT and Muc2$^{DTA/+}$ mice (n = 9, 9, 10 and 11 for SPF WT, SPF Muc2$^{DTA/+}$, GF WT and GF Muc2$^{DTA/+}$). Two-way ANOVA. MCP-1: main effect of housing (***p < 0.001), interaction ns. IFN-γ: main effect of housing (**p < 0.01), interaction ns. (D) MEICS scores of untreated and antibiotic (ABX)-treated WT and Muc2$^{DTA/+}$ mice at 20 weeks of age (n = 2, 5, 4 and 4 for WT, Muc2$^{DTA/+}$, WT + ABX and Muc2$^{DTA/+}$ + ABX). Two-way ANOVA (genotype × treatment) with Tukey's multiple comparisons test (interaction, p = 0.009). (E) Colonic crypt length in untreated and ABX-treated WT and Muc2$^{DTA/+}$ mice at 20 weeks of age (n = 5, 5, 5 and 4 for WT, Muc2$^{DTA/+}$, WT + ABX and Muc2$^{DTA/+}$ + ABX). Two-way ANOVA (genotype × treatment) with Tukey's multiple comparisons test (interaction, p < 0.0001). Dots represent individual mice, bars show mean ± SEM and n indicates individual mice. * and # denote main effects of housing and genotype, respectively.

#### Fig. S6

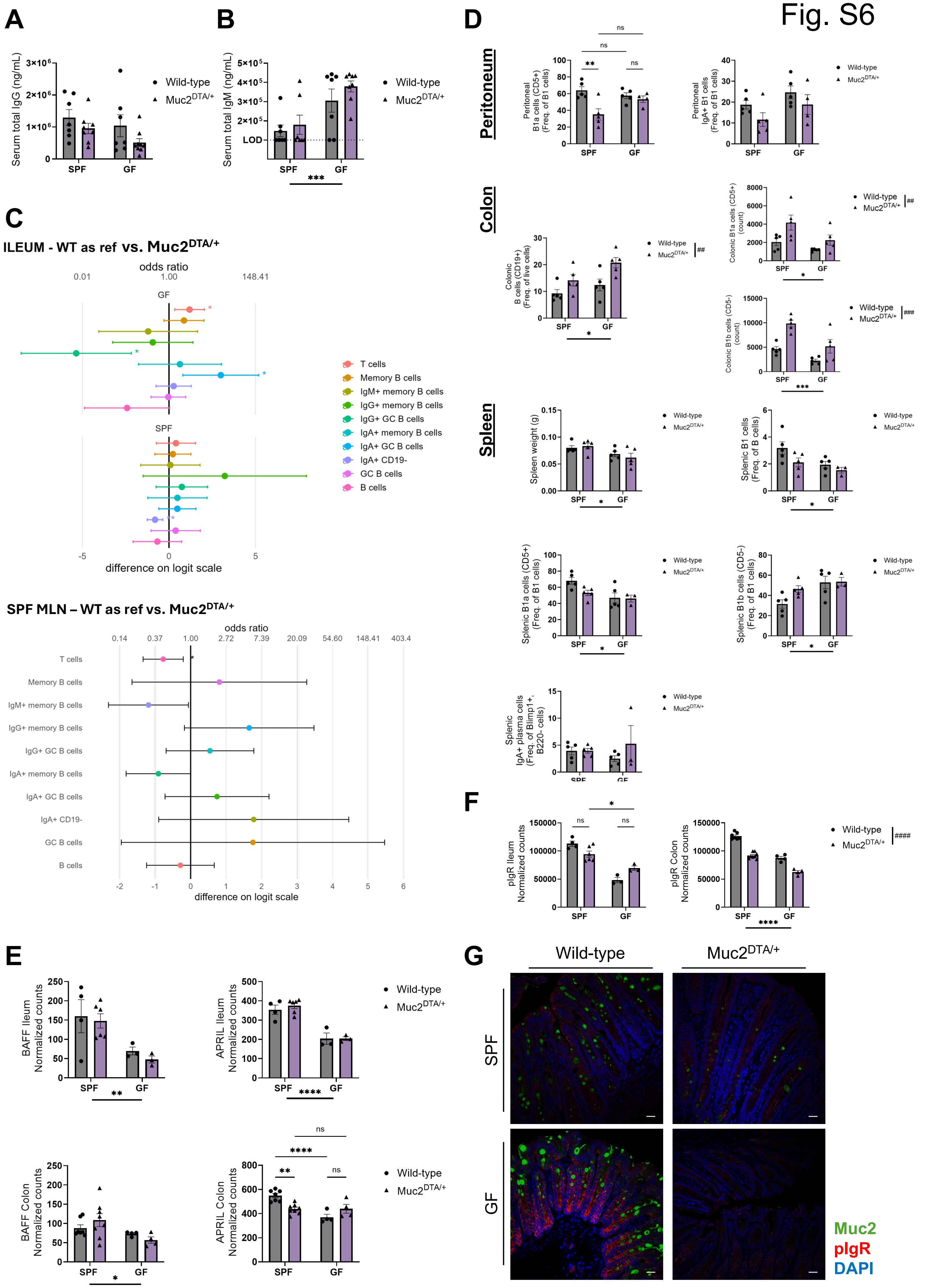

A
Serum total IgG (ng/mL)
Wild-type
Muc2DTA/+
SPF
GF
B
Serum total IgM (ng/mL)
LOD
***
C
ILEUM - WT as ref vs. Muc2DTA/+
odds ratio
difference on logit scale
T cells
Memory B cells
IgM+ memory B cells
IgG+ memory B cells
IgG+ GC B cells
IgA+ memory B cells
IgA+ GC B cells
IgA+ CD19-
GC B cells
B cells
SPF MLN – WT as ref vs. Muc2DTA/+
D
Peritoneum
Colon
Spleen
Peritoneal B1a cells (CD5+) (Freq. of B1 cells)
Peritoneal IgA+ B1 cells (Freq. of B1 cells)
Colonic B cells (CD19+) (Freq. of live cells)
Colonic B1a cells (CD5+) (count)
Colonic B1b cells (CD5-) (count)
Spleen weight (g)
Splenic B1 cells (Freq. of B cells)
Splenic B1a cells (CD5+) (Freq. of B1 cells)
Splenic B1b cells (CD5-) (Freq. of B1 cells)
Splenic IgA+ plasma cells (Freq. of Blimp1+, B220- cells)
E
BAFF Ileum Normalized counts
APRIL Ileum Normalized counts
BAFF Colon Normalized counts
APRIL Colon Normalized counts
F
pIgR Ileum Normalized counts
pIgR Colon Normalized counts
G
Wild-type
Muc2DTA/+
SPF
GF
Muc2
pIgR
DAPI

**Figure S6: Additional characterization of the B cell and IgA response in SPF and GF Muc2$^{DTA/+}$ mice** (A) Serum IgG in SPF and GF WT and Muc2$^{DTA/+}$ mice (n = 7, 8, 7 and 9). Two-way ANOVA: no main effects, interaction ns. (B) Serum IgM in SPF and GF WT and Muc2$^{DTA/+}$ mice (n = 7, 7, 7 and 8). GLM: main effect of housing (***$p < 0.001$); genotype and interaction, ns. LOD, limit of detection, 100,000. (C) Forest plots of B cell populations in ileum and mesenteric lymph nodes of Muc2$^{DTA/+}$ mice relative to WT, under SPF and GF conditions (n = 7, 8, 4 and 4). Points show the difference on the logit scale with 95% CI error bars. HGLM; R. *$p < 0.05$, **$p < 0.01$. (D) B1 cell populations in the peritoneum, colon and spleen of SPF and GF WT and Muc2$^{DTA/+}$ mice (n = 5 per group, except peritoneal GF Muc2$^{DTA/+}$, n = 4). Two-way ANOVA. Peritoneum: B1a cells, interaction ($p = 0.0006$); IgA+ cells, all effects ns. Colon: B cells, main effects of housing (*$p = 0.024$) and genotype (##$p = 0.004$), interaction ns; B1a cells, housing (*$p = 0.02$) and genotype (##$p = 0.009$), interaction ns; B1b cells, housing (***$p = 0.0006$) and genotype (###$p = 0.0002$), interaction ns. Spleen: weight, main effect of housing (*$p = 0.013$); B1 cells, housing (*$p = 0.026$); B1a cells, housing (*$p = 0.012$); B1b cells, housing (*$p = 0.011$); IgA+ plasma cells, all effects ns; interaction ns throughout. (E) *Tnfsf13b* (BAFF) and *Tnfsf13* (APRIL) expression in ileum and colon, from bulk RNA sequencing of epithelial cells from SPF and GF WT and Muc2$^{DTA/+}$ mice. Two-way ANOVA. BAFF: main effect of housing in ileum (**$p = 0.0047$) and colon (*$p = 0.0281$), interaction ns. APRIL: main effect of housing in ileum (****$p < 0.0001$), interaction ns; in colon, interaction ($p = 0.026$). (F) Normalized *pIgR* transcript counts from bulk RNA sequencing of ileal and colonic tissue from SPF and GF WT and Muc2$^{DTA/+}$ mice. Two-way ANOVA. Ileum: genotype × housing with Tukey's multiple comparisons test (interaction, $p = 0.004$). Colon: main effects of housing (****$p < 0.0001$) and genotype (####$p < 0.0001$), interaction ns. (G) Immunostaining for pIgR in colonic sections of SPF and GF WT and Muc2$^{DTA/+}$ mice. Scale bar, 20 µm. Dots represent individual mice, bars show mean ± SEM and n indicates individual mice. Where four groups are shown, n is given in the order SPF WT, SPF Muc2$^{DTA/+}$, GF WT and GF Muc2$^{DTA/+}$. * and # denote main effects of housing and genotype, respectively.

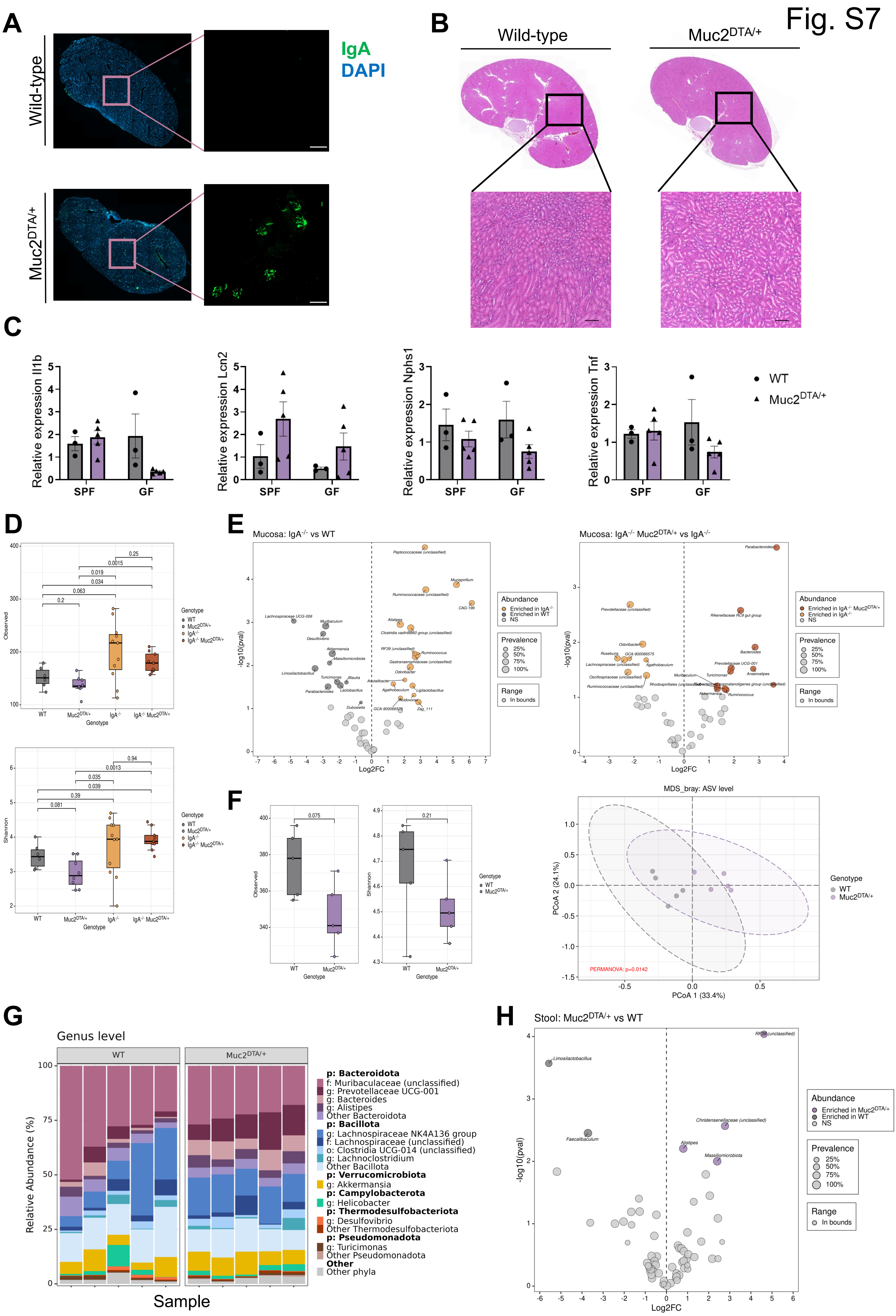

Fig. S7
A
IgA
DAPI
Wild-type
Muc2DTA/+
B
Wild-type
Muc2DTA/+
C
Relative expression Il1b
Relative expression Lcn2
Relative expression Nphs1
Relative expression Tnf
SPF
GF
WT
Muc2DTA/+
D
Genotype
Observed
Shannon
E
Mucosa: IgA-/- vs WT
Mucosa: IgA-/- Muc2DTA/+ vs IgA-/-
Log2FC
-log10(pval)
Abundance
Prevalence
Range
F
MDS_bray: ASV level
PCoA 1 (33.4%)
PCoA 2 (24.1%)
PERMANOVA: p=0.0142
G
Genus level
Relative Abundance (%)
Sample
p: Bacteroidota
f: Muribaculaceae (unclassified)
g: Prevotellaceae UCG-001
g: Bacteroides
g: Alistipes
Other Bacteroidota
p: Bacillota
g: Lachnospiraceae NK4A136 group
f: Lachnospiraceae (unclassified)
o: Clostridia UCG-014 (unclassified)
g: Lachnoclostridium
Other Bacillota
p: Verrucomicrobiota
g: Akkermansia
p: Campylobacterota
g: Helicobacter
p: Thermodesulfobacteriota
g: Desulfovibrio
Other Thermodesulfobacteriota
p: Pseudomonadota
g: Turicimonas
Other Pseudomonadota
Other
Other phyla
H
Stool: Muc2DTA/+ vs WT

**Figure S7: Kidney and gut microbiota analysis of Muc2$^{DTA/+}$ mice** (A) Representative immunofluorescence image of a whole kidney section stained for IgA (green) and DAPI (blue). (B) Representative H&E-stained kidney sections of WT and Muc2$^{DTA/+}$ mice showing no histological signs of kidney disease. Scale bars, 100 µm. (C) RT-qPCR analysis of *Il1b*, *Lcn2, Nphs1* and *Tnf* expression in kidney tissue of SPF and GF WT and Muc2$^{DTA/+}$ mice (n = 3, 5, 3, 5 for SPF WT, SPF Muc2$^{DTA/+}$, GF WT and GF Muc2$^{DTA/+}$.). *Il1b*: GLM (Poisson, log link, estimated dispersion), genotype × housing interaction p = 0.011 (Wald t-test). Two-way ANOVA: *Tnf*, *Lcn2* and *Nphs1*, all effects ns. (D) Mucosal microbiota diversity by 16S rRNA sequencing (ASV level; n = 6, 8, 11, 9). Alpha diversity: observed ASVs (above) and Shannon index (below). Boxes show median and interquartile range; dots are individual mice; numbers above brackets are pairwise p-values. (E) Differential abundance of mucosal genera between 1) IgA$^{-/-}$ and WT mice and 2) IgA$^{-/-}$ Muc2$^{DTA/+}$ and IgA$^{-/-}$ mice, shown as volcano plots of log2 fold change against -log10 p-value. Colors indicate enrichment; point size indicates prevalence (n = 6, 8, 11, 9 for WT, Muc2$^{DTA/+}$, IgA$^{-/-}$, IgA$^{-/-}$ Muc2$^{DTA/+}$). (F) Stool microbiota of WT and Muc2$^{DTA/+}$ mice (n = 5 per group) by 16S rRNA sequencing at the ASV level. 1) Alpha diversity: observed ASVs (left) and Shannon index (right); boxes show median and interquartile range, dots are individual mice. 2) PCoA of Bray-Curtis dissimilarity. PERMANOVA, p = 0.0142. (G) Genus-level relative abundance (%) in stool of WT and Muc2$^{DTA/+}$ mice (n = 5 per group); each bar is one mouse. Genera are grouped by phylum. (H) Volcano plot of differential abundance of stool genera, Muc2$^{DTA/+}$ versus WT (n = 5 per group). Colors indicate enrichment; point size indicates prevalence. n indicates individual mice.

Fig. S8

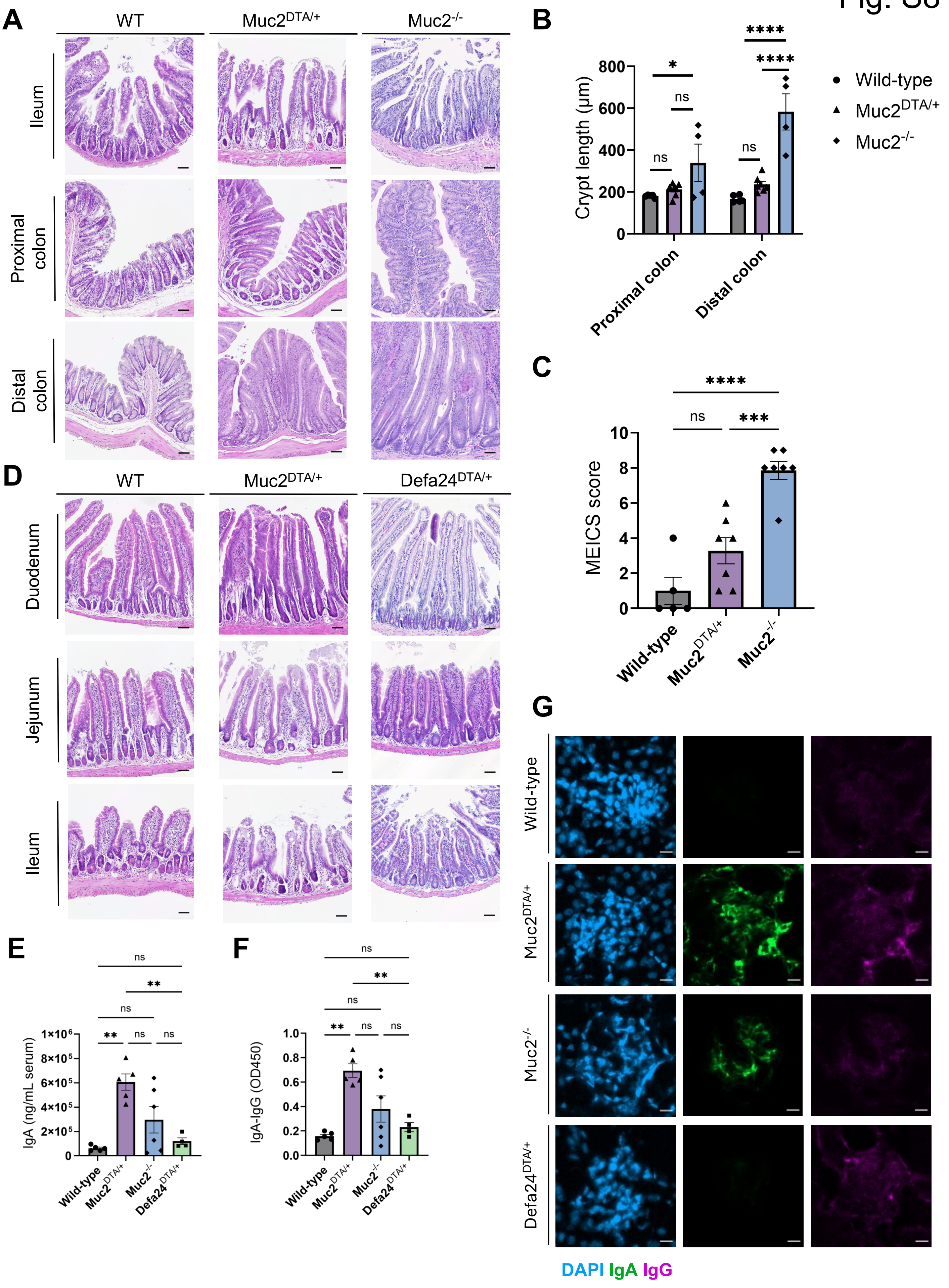

A
WT
Muc2DTA/+
Muc2-/-
Ileum
Proximal colon
Distal colon
B
Crypt length (μm)
800
600
400
200
0
*
ns
****
Proximal colon
Distal colon
Wild-type
Muc2DTA/+
Muc2-/-
C
MEICS score
10
8
6
4
2
0
****
***
ns
Wild-type
Muc2DTA/+
Muc2-/-
D
WT
Muc2DTA/+
Defa24DTA/+
Duodenum
Jejunum
Ileum
E
IgA (ng/mL serum)
1×10⁶
8×10⁵
6×10⁵
4×10⁵
2×10⁵
0
ns
**
Wild-type
Muc2DTA/+
Muc2-/-
Defa24DTA/+
F
IgA-IgG (OD450)
1.0
0.8
0.6
0.4
0.2
0.0
ns
**
Wild-type
Muc2DTA/+
Muc2-/-
Defa24DTA/+
G
Wild-type
Muc2DTA/+
Muc2-/-
Defa24DTA/+
DAPI IgA IgG

**Figure S8: IgA dysregulation is strongest with goblet cell depletion, intermediate with Muc2 deficiency and absent with Paneth cell depletion** (A) Representative H&E-stained sections of ileum and colon from WT, $Muc2^{DTA/+}$ and $Muc2^{-/-}$ mice. Scale bars, 50 µm. (B) Crypt length (µm) in proximal and distal colon of WT (n = 5), $Muc2^{DTA/+}$ (n = 7) and $Muc2^{-/-}$ (n = 4) mice. Bars show mean ± SEM of untransformed values. Two-way ANOVA with Tukey's multiple comparisons test on log10-transformed data (interaction, $p = 0.02$). (C) MEICS scores from colonoscopy recordings of WT (n = 5), $Muc2^{DTA/+}$ (n = 7) and $Muc2^{-/-}$ (n = 7) mice at 10 weeks of age. Ordinary one-way ANOVA with Tukey's multiple comparisons test: ***$p = 0.0003$, ****$p < 0.0001$; ns, not significant. (D) Representative H&E-stained sections of duodenum, jejunum and ileum from WT, $Muc2^{DTA/+}$ and $Defa24^{DTA/+}$ mice. Scale bars, 50 µm. (E) Serum IgA (ng/mL) in WT (n = 5), $Muc2^{DTA/+}$ (n = 5), $Muc2^{-/-}$ (n = 6) and $Defa24^{DTA/+}$ (n = 4) mice. Welch's ANOVA with Dunnett's T3 multiple comparisons test, because variances were unequal (Brown-Forsythe test, $p = 0.0004$). **$p < 0.01$; ns, not significant. (F) Serum circulating IgA-IgG complexes (OD 450) in WT (n = 5), $Muc2^{DTA/+}$ (n = 5), $Muc2^{-/-}$ (n = 6) and $Defa24^{DTA/+}$ (n = 4) mice. Welch's ANOVA with Dunnett's T3 multiple comparisons test, because variances were unequal (Brown-Forsythe test, $p = 0.0018$). **$p < 0.01$; ns, not significant. (G) IgA and IgG staining of kidney sections from $Muc2^{-/-}$ and $Defa24^{DTA/+}$ mice. Scale bar, 10 µm. Dots represent individual mice, bars show mean ± SEM and n indicates individual mice.

Fig. S9

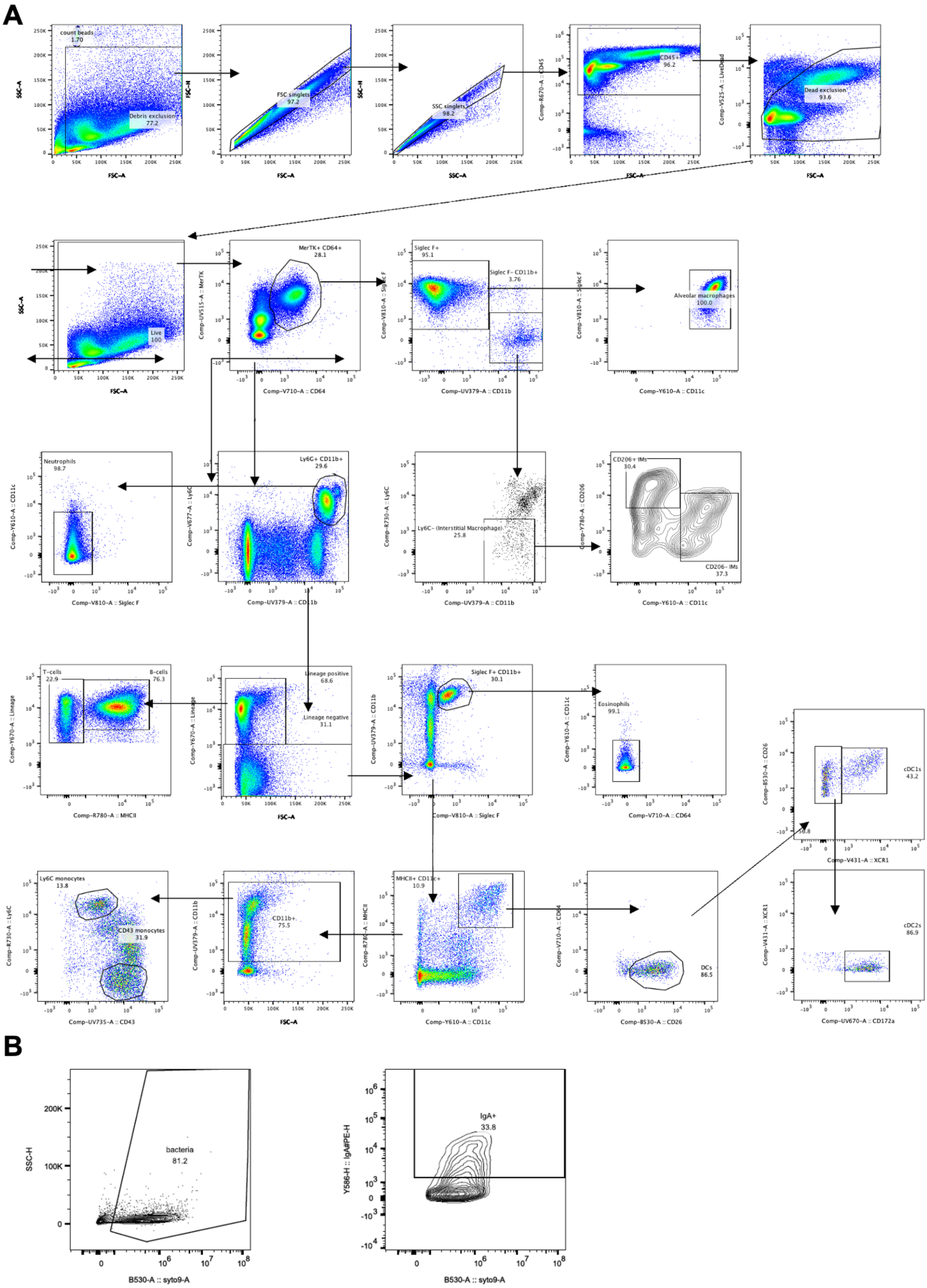

**Figure S9: Gating strategy for myeloid cells and for the analysis of IgA-coated bacteria**

Fig. S10

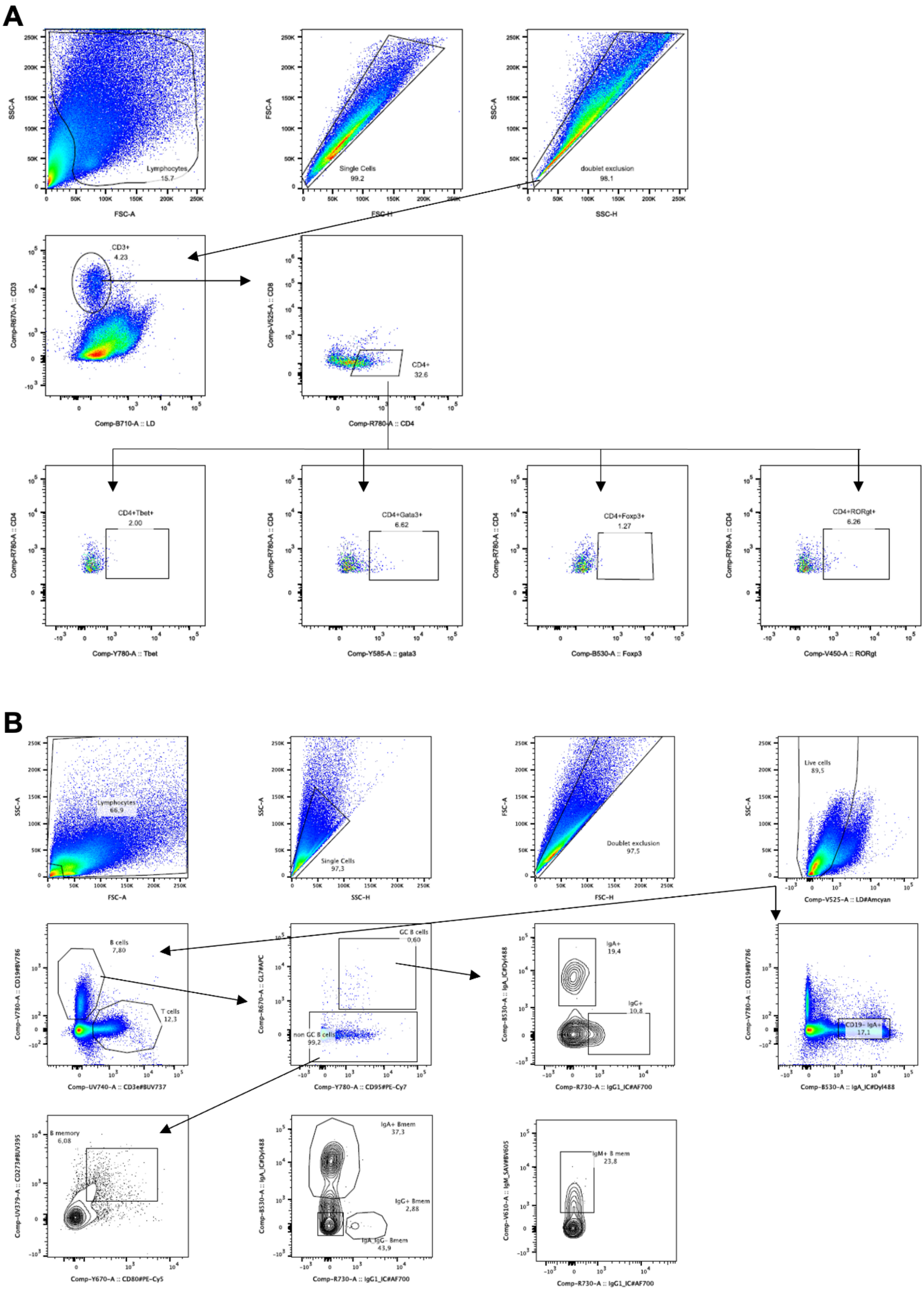

**Figure S10: Gating strategy for T cells and B cells**

Fig. S11

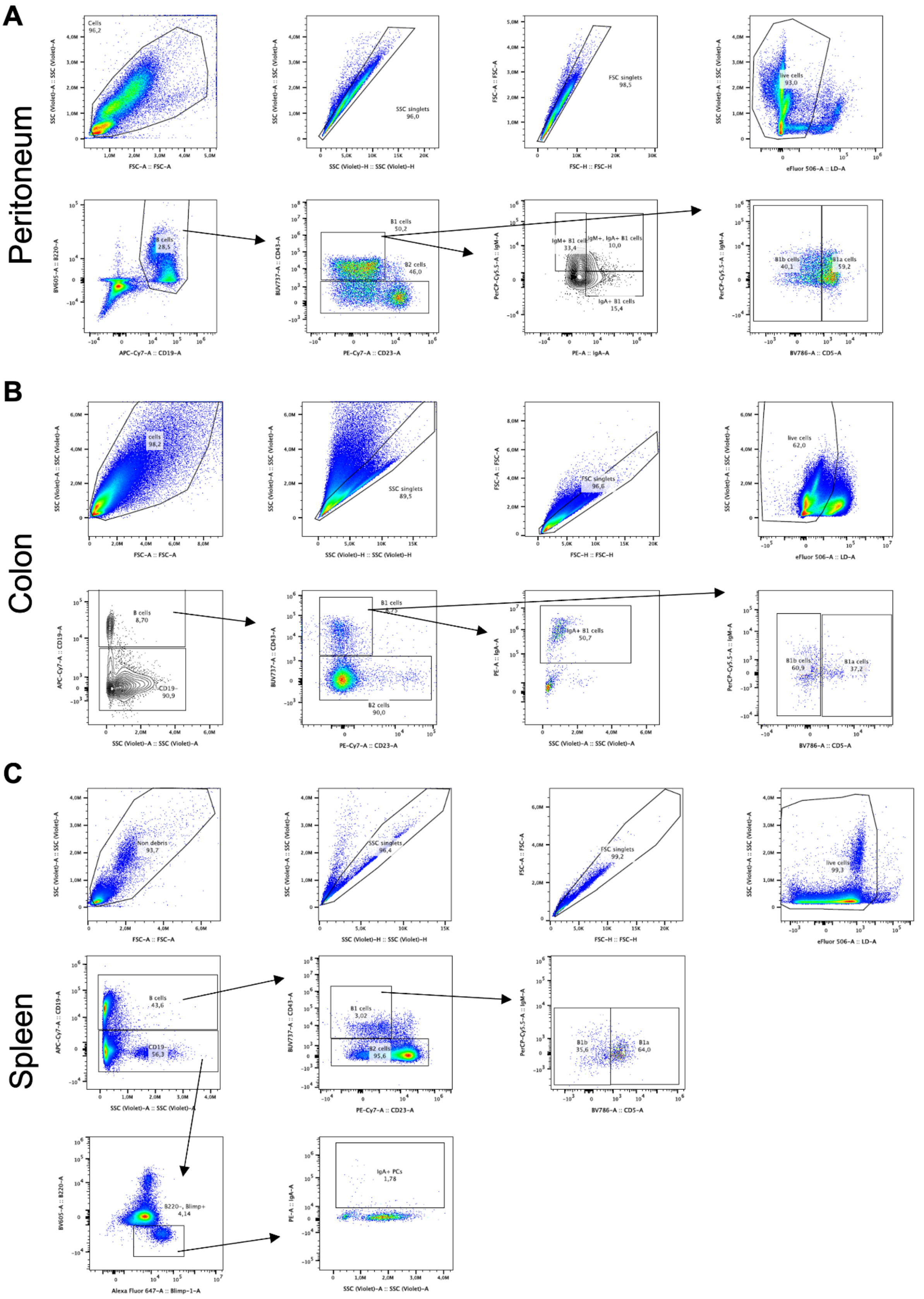

**Figure S11: Gating strategy for B1 cells**

#### Supplementary Tables

***Supplementary Table 1:*** Marker gene sets used for cell type annotation of epithelial cells in scRNA-seq data

| Goblet | Paneth | Tuft | Enteroendocrine | Enterocyte | Proximal | Distal | Intercrypt |
|---|---|---|---|---|---|---|---|
| Agr2 | Gm15284 | Alox5ap | Chgb | Gsta1 | Car8 | B3gnt7 | Slfn4 |
| Fcgbp | AY761184 | Lrmp | Gfra3 | Rbp2 | | | Mxd1 |
| Tff3 | Defa17 | Hck | Cck | Adh6a | | | Fer1l6 |
| Clca1 | Gm14851 | Avil | Vwa5b2 | Apoa4 | | | Cyp2d34 |
| Zg16 | Defa22 | Rgs13 | Neurod1 | Reg3a | | | Krt20 |
| Tpsg1 | Defa-rs1 | Ltc4s | Fev | Creb3l3 | | | Cldn4 |
| Muc2 | Defa3 | Trpm5 | Aplp1 | Cyp3a13 | | | Cyp2d12 |
| Guca2a | Defa24 | Dclk1 | Scgn | Cyp2d26 | | | |
| S100a6 | Defa26 | Spib | Neurog3 | Ms4a10 | | | |
| AW112010 | Defa21 | Fyb | Resp18 | Ace | | | |
| Klk1 | Lyz1 | Ptpn6 | Trp53i11 | Aldh1a1 | | | |
| Tmsb10 | Gm15292 | Matk | Bex2 | Rdh7 | | | |
| | Mptx2 | Snrnp25 | Rph3al | H2-Q2 | | | |
| | Ang4 | Sh2d7 | Scg5 | Hsd17b6 | | | |
| | | Ly6g6f | Pcsk1 | Gstm3 | | | |
| | | Kctd12 | Isl1 | Gda | | | |
| | | 1810046K07Rik | Maged1 | Apoc3 | | | |
| | | Hpgds | Fabp5 | Gpd1 | | | |
| | | Tuba1a | Celf3 | Fabp1 | | | |
| | | Pik3r5 | Pcsk1n | Slc5a1 | | | |
| | | Vav1 | Fam183b | Mme | | | |
| | | Tspan6 | Prnp | Cox7a1 | | | |
| | | Skap2 | Tac1 | Gsta4 | | | |
| | | Pygl | Gpx3 | Lct | | | |
| | | Ccdc109b | Cplx2 | Khk | | | |
| | | Ccdc28b | Nkx2-2 | Mttp | | | |
| | | Plcg2 | Olfm1 | Xdh | | | |
| | | Ly6g6d | Vim | Sult1b1 | | | |
| | | Alox5 | Rimbp2 | Treh | | | |
| | | Pou2f3 | Anxa6 | Lpgat1 | | | |
| | | Gng13 | Scg3 | Dhrs1 | | | |
| | | Bmx | Ngfrap1 | Cyp2c66 | | | |
| | | Ptpn18 | Insm1 | Ephx2 | | | |
| | | Nebl | Gng4 | Cyp2c65 | | | |

| | | | | |
|---|---|---|---|---|
| | | Limd2 | Pax6 | Cyp3a25 |
| | | Pea15a | Cnot6l | Slc2a2 |
| | | Tmem176a | Cacna2d1 | Ugdh |
| | | Smpx | Tox3 | Gstm6 |
| | | Itpr2 | Slc39a2 | Retsat |
| | | Il13ra1 | Riiad1 | Ppap2a |
| | | Siglecf | | Acsl5 |
| | | Ffar3 | | Cyb5r3 |
| | | Rac2 | | Cyb5b |
| | | Hmx2 | | Ckmt1 |
| | | Bpgm | | Aldob |
| | | Inpp5j | | Ckb |
| | | Ptgs1 | | Scp2 |
| | | Aldh2 | | Prap1 |
| | | Pik3cg | | Tmigd1 |
| | | Cd24a | | Fabp6 |
| | | Ethe1 | | Slc51b |
| | | Inpp5d | | Slc51a |
| | | Krt23 | | Mep1a |
| | | Gprc5c | | Fam151a |
| | | Reep5 | | Naaladl1 |
| | | Csk | | Slc34a2 |
| | | Bcl2l14 | | Plb1 |
| | | Tmem141 | | Nudt4 |
| | | Coprs | | Dpep1 |
| | | Tmem176b | | Pmp22 |
| | | 1110007C09Rik | | Xpnpep2 |
| | | Ildr1 | | Muc3 |
| | | Galk1 | | Neu1 |
| | | Zfp428 | | Clec2h |
| | | Rgs2 | | Phgr1 |
| | | Inpp5b | | 2200002D01Rik |
| | | Gnai2 | | Prss30 |
| | | Pla2g4a | | Cubn |
| | | Acot7 | | Plec |
| | | Rbm38 | | Fgf15 |
| | | Gga2 | | Crip1 |
| | | Myo1b | | Krt20 |
| | | Adh1 | | Dhcr24 |

| | | | | |
|---|---|---|---|---|
| | | Bub3 | | Myo15b |
| | | Sec14l1 | | Amn |
| | | Asah1 | | Enpep |
| | | Ppp3ca | | Anpep |
| | | Agt | | Slc7a9 |
| | | Gimap1 | | Ocm |
| | | Krt18 | | Anxa2 |
| | | Pim3 | | Aoc1 |
| | | 2210016L21Rik | | Ceacam20 |
| | | Tmem9 | | Arf6 |
| | | Lima1 | | Abcb1a |
| | | Fam221a | | Xpnpep1 |
| | | Nt5c3 | | Vnn1 |
| | | Atp2a3 | | Cndp2 |
| | | Mlip | | Nostrin |
| | | Vdac3 | | Slc13a1 |
| | | Ccdc23 | | Aspa |
| | | Tmem45b | | Maf |
| | | Cd47 | | Myh14 |
| | | Lect2 | | |
| | | Pla2g16 | | |
| | | Mocs2 | | |
| | | Arpc5 | | |
| | | Ndufaf3 | | |

**Addendum**

### Report

#### Prepared for:

Company name: VIB Nucleomics Core
Customer name: Dr. Lars Vereecke
Quote number: 2020VIB0401
Project number: 1435
Version: 1
Date: July 2020

Cergentis B.V. Yalelaan 62 3584 CM Utrecht The Netherlands +31 30 760 1636 www.cergentis.com

### Goal

In this study, 1 transgenic mouse with the vector injection fragment Muc2-iCre sequence was analysed.
The aim of this analysis was to:

1. Sequence the vector;
   1) Determine the presence of sequence variants and their allele frequency.
   2) Determine the presence of vector-vector breakpoints that represent concatemerization of multiple copies of the vector and/or structural rearrangements in a single vector sequence.
2. Identify vector integration site(s) and breakpoint sequences between the vector and genome.
3. Assess the presence of structural variants surrounding the vector integration site(s).
4. Estimate the copy number of the vector.

An overview of the TLA technology and technical details of the performed analyses is provided in the manual "Introduction to the terminology and methods used in TLA analyses".

### Summary

| Sample | Integration site(s) | Sequence and structural variants in vector | Structural variants at the integration site | Notes |
|---|---|---|---|---|
| Muc2iCre | chr3:133,465,565-133,468,815. | 26 sequence variants, 2 structural variants | | Complex integration event |

### Conclusion

In this sample a complex integration event was found on chromosome 3. According to the RefSeq the integrity of Tet2 may have been affected. Furthermore 26 sequence variants and 2 structural variants were observed.

#### TLA, sequencing and data mapping

Viable frozen mouse spleen cells were used and processed according to Cergentis' TLA protocol (de Vree et al. Nat Biotechnol. Oct 2014). An overview of the TLA technology and technical details of the performed analyses is provided in the manual "Introduction to the terminology and methods used in TLA analyses".

TLA was performed with 3 independent primer sets specific for the vector sequence (Table 1).

**Table 1**: Primers used in TLA analysis

| Primer set | Name/View point | Direction | Binding position | Sequence |
|---|---|---|---|---|
| 1 | 29 KB | RV | 29,937 | ATAAATTATGCCAGGTGCCT |
| | | FW | 30,140 | CTATGTAGGACAGGGTGAAC |
| 2 | iCre | RV | 41,271 | ATCTTCCAGGTGTGTTCAGA |
| | | FW | 41,377 | ACTACCTCCTGTACCTGCAA |
| 3 | 81 KB | RV | 81,144 | TGTTGGACTAGAAACCAGTG |
| | | FW | 81,238 | ACTTTGGCTAGCATTGCTAA |

The NGS reads were aligned to the vector sequence and host genome. The mouse mm10 genome was used as host reference genome sequence.

### Results Muc2iCre

#### Vector integrity

Figure 1 depicts the NGS coverage across the vector sequence using primer sets 1-3.

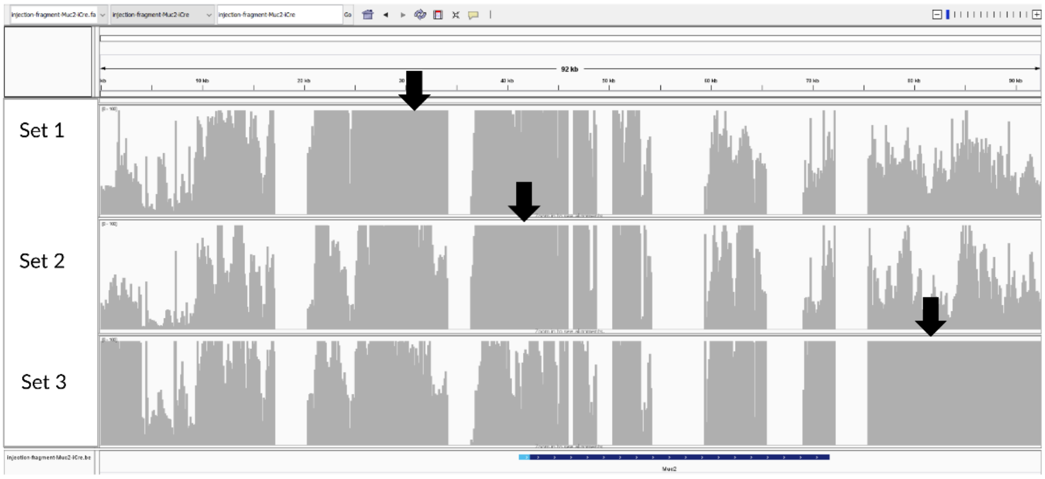

**Figure 1:** NGS sequencing coverage (in grey) across the vector. Black arrows indicate the primer location. The blue arrows indicate the locations of the identified vector-genome breakpoint sequences (described below). The vector map is shown on the bottom. Y-axes are limited to 100x.

Coverage is observed across the vector sequence Vector: 1-92,973, except for 14,548-14,620; 17,038-20,382, 34,079-36,444; 45,867-46,501; 48,670-50,367; 54,112-59,408; 65,472-69,140, 72,221-75,525 due to the gaps (NNNNNN) in the provided reference sequence. No coverage is observed across the Vector: 32,635-32,640 due a deletion described below in sequence variants. Local dips in coverage are due to GC rich regions that are less efficiently sequenced.

The gaps (NNNNNN in provided reference sequence) in coverage in the vector are bordered by the following reads:
Vector:14,548 (tail) fused to unknown
TCGGGGCCAACTGTATGTCTACATGTGTGTGTATTTGAGGGTGTATGTGTGTGTATGTGCGTGTATGTGTGCTTGTG
TGTGTATGTGTGCGTGTTCGTGTGCGTGTGTGCTCATG

Vector: 17,034 (tail) fused to unknown
AAAAAAAGTGGTCAGGGAAATGCAAATCAAAATGACCCTGAGATTCTACCTTATATCAGTCGGAATGGCTAAGATC
AAAAACTCAGGTGACAGCACATG

Chr7: 141668983 (tail, downstream of endogenous Muc2) fused to Vector: 20,382 (head)
GCTCTAGACAGAAAGGAGTAAGGATGGGACTTTTTGCATATTTTCACTTGGGACCTTCTTTAGGTGCTCCTGGTCCT
TGTGAGGGAAGCTGCACATCTGGTTTCCAGAGGATGGGCAGCGAGGTGCACCCCTAAGGAGTTATGAATGAGGG

Vector: 34,079 (tail) fused to unknown
CATGAACACAGAGGAAAACATTCTAAATGTTGACAAATGAAATCCAAAGTGATTAAGATGTGACATG

Chr 7: 141,685,202 (tail, downstream of endogenous Muc2) fused to vector: 36,445 (head)
GCTAAGCATAGAAGCGCTATATAATACAGAAATTCTCTCTGTTGGTTAGAAACAGAAGAATCCAGAGCAGAGTCTT
AACGATATTTGTACACTGATGATGAAGACAGCAGGAAAGACGACAACTGCGCTGTGGGAAACAACCCTAGTACCA

Vector: 45,867 (tail) fused to chr7: 141,694,177 (head, endogenous Muc2)
CAGGGAGGGTTGGGACCCTCCCAAGAGCCCTGAGGGCTGTGAGATGAGAACCTTCATTGACAGGTCAGGGCTCCA
GACACCTGTGGGTGGAGGTCTGGCTGGAAGCCTCACCTTGGGGGAGACTGGGACCTCCTGAG

Chr7: 141,694,531 (tail, endogenous Muc2) fused to vector: 46,501
ACCAATGACTGTGAGCAGTGGTGAGTCTCAGAGTAAAGAGGTTGGGGGTCAGACAGGCTGGGGTGTTACCGAGTC
TTGACGCTGTGCCCTGCTTCTGGGCAGTGTCTGCAATGCTGGTCGCTGGATGTGTAAAGACCTGCCTTGCCCTGAG

Vector: 48,670 (tail) fused to chr 7: 141,696,790 (head, endogenous Muc2)
GAGTGCTTCCCCACTGACTGATTCCTACCTAAGATGATGATTTAGGCTTGGGCTGGGAGCCCTGAGCTCAGCAGTC
CAGGGAGAGGGGCCAGTTCCTTGTAGCTGCCTCCCGGCTTCTGCCAGAAGCAGCAGATTTCCCTGAGTTTTTAGA

Unknown fused to vector: 59,408 (head)
CAGACAGGGACTCCACCCTCAACACCAACCACCACAGGGCCCACAACACCAACATCTACTACCACCACACAGACC
ACAACTCCAACACCCAACATAGAGACCTCCACTCCTACATCAACCACTACTCAGACACCAAGTCCAACCCCTACAA

Vector: 65,472 (tail) fused to chr7: 141,749,326 (head, endogenous Muc2)
ATGGCCTTCCTTATTCTCGTCCCTTCCCTGCAGCTGTGACGTGTCCAGCTCACAGGCAGTATCAGGCCTGTGGCCCT
TCAGAGGAACCCACCTGCCAGTCCAGGTATGCACCCAAGATCCCTGCTGCTTCCCTAGTCCTGCTCATG

Chr7:141,752,183 (tail, endogenous Muc2) fused to vector: 69,140 (head)
GCTCAGGATAGCCTCTGATGGCTTTCCCCACTCTTTCCTCCAAGTGCCCAAGGGTGTGTGCGTGCACGAGAATGCT
GAATACCAGGTGAGTTCTGAGCTGGGAGGGGATGGGGGTGGGGGACGGGTGGCGTTGCCCTTGCCTGAAGATGG

Chr7: 141,758,933(tail, endogenous Muc2) fused to vector: 75,525 (head)
CATGGGCACTGGGAAATGTAGCCACAGGATGAGTGTATGATCAGTCCTATTAGAGGGGACCTGAGACCAGTTTAG
GCCAGGTTGTCAGAGGGAAGATAGAGCCTTCCTCTTGGACTATCAAGT

Due to a CATG site near the unknown region in 14,620, 50,367, 54,110 a fusion read could not be identified. Due to a repetitive sequence before the unknown region a fusion read at 72,207 could not be identified.

Sequence variants and structural variants were called in the covered regions.

**Sequence variants**
Detected sequence variants are presented in table 2. Sequence variants near or at 100% mutation frequency are most likely represent deviations present in the provided reference sequence of the vector before its introduction into the sample.

**Table 2:** Identified sequence variants

| | | | | Primer set 1 | | Primer set 2 | |
|---|---|---|---|---|---|---|---|
| Region | Position | Reference | Mutation | Coverage | % | Coverage | % |
| Not annotated | 1,215 | G | -4GATC | 20 | 85 | 42 | 71 |
| Not annotated | 3,590 | C | A | 31 | 61 | 15 | 33 |
| Not annotated | 7,227 | C | +1A | 21 | 48 | 20 | 65 |
| Not annotated | 7,238 | G | A | 31 | 32 | 25 | 52 |
| Not annotated | 7,246 | A | G | 34 | 29 | 27 | 48 |
| Not annotated | 9,162 | T | +2AG | 47 | 89 | 20 | 70 |
| Not annotated | 14,538 | A | T | 96 | 86 | 11 | 82 |
| Not annotated | 16,879 | G | A | 116 | 28 | 75 | 20 |
| Not annotated | 17,035 | N | G | 69 | 100 | 30 | 100 |
| Not annotated | 17,036 | N | G | 69 | 100 | 30 | 100 |
| Not annotated | 17,037 | N | A | 69 | 100 | 30 | 100 |
| Not annotated | 17,038 | N | A | 69 | 100 | 28 | 100 |
| Not annotated | 21,036 | A | G | 105 | 30 | 42 | 60 |
| Not annotated | 32,634 | T | +6TTGAAA | 145 | 74 | 57 | 68 |
| Not annotated | 32,635 | C | -5GCCCC | 107 | 100 | 39 | 100 |
| Not annotated | 32,778 | N | T | 119 | 99 | 53 | 100 |
| Not annotated | 36,444 | N | C | 18 | 100 | 49 | 100 |
| Not annotated | 40,536 | A | +3CTG | 90 | 87 | 148 | 91 |
| Muc2 | 44,624 | A | +1G | 100 | 92 | 397 | 93 |
| Muc2 | 59,455 | G | T | 43 | 23 | 142 | 23 |
| Muc2 | 59,465 | T | C | 23 | 65 | 98 | 78 |
| Muc2 | 59,468 | G | A | 23 | 70 | 96 | 78 |
| Muc2 | 59,476 | G | C | 21 | 86 | 96 | 81 |
| Not annotated | 82,300 | G | T | 28 | 100 | 20 | 100 |
| Not annotated | 82,330 | A | +1G | 31 | 81 | 16 | 81 |
| Not annotated | 90,533 | G | T | 31 | 26 | 25 | 28 |

**Structural variants**
2 vector-vector breakpoints were found.

Vector: 7,963 (head) fused to Vector: 70,221 (tail) with 4bp homology

GATATGCTCTGAGCCAACTAAGGACAGCATCTGAGAGCTCCCAGTGTGGGGGACTAGAGGAGATATGCTAGGTCC
AAAGTAGGGGACTTGGCCTAGCTTGGCTGTGAGATACAGGCCAGACTGAAACAAGGCTAGATAACACAAAAATCT
CTAACCTACGTCTAACCCGTGAACTAGAAAGTGACTCTGAATGGAGAAATGATCTCTGAAAAGCTCAAGAAAATAA
TCTGGAGATGAAGTCGCCCTACATCAAAGAGGTCCTAAC

Vector: 9,561 (tail) fused to Vector: 90,890 (tail) with 4bp homology

CACGTATGTGTGCCTTCCAGGATCTTCTAGGCTGCGTACGCCTGTGGCTAAAAGGTAGCTCCAGGGAGTGGCACTG
CCCTTGCCTAGGTCAATGGCTCTATCCTGTTGAGGAGGGGCCACAGCAGGGCATCAGGATGGACTGCATCCCTGA
AATATGAGTCGGAAAAAGCCCCGTATTAAGTTGCTTTTGTTGGGTACTTGGTCATGTGAAGAGTAACTGACTAACA

#### Integration sites

**Whole genome coverage plot**

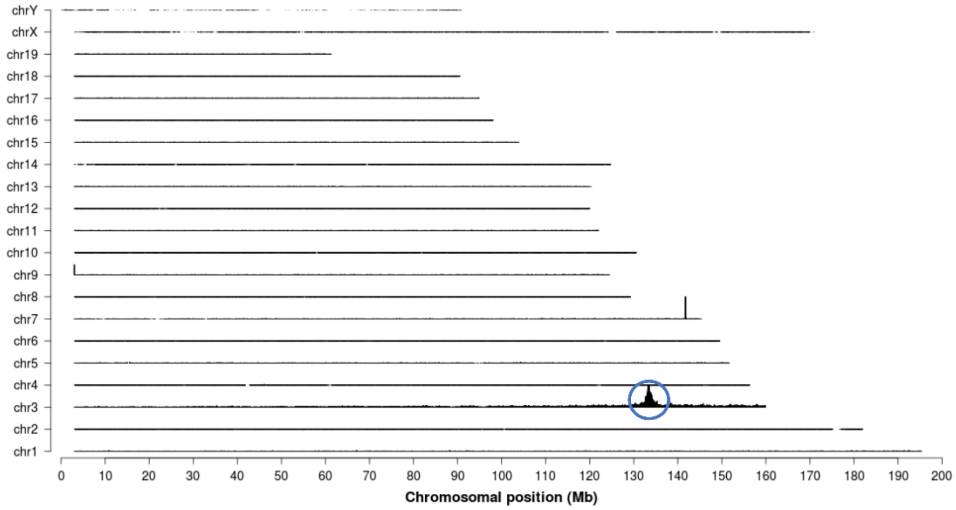

**Figure 2:** TLA sequence coverage across the mouse genome using primer set 3. The chromosomes are indicated on the y-axis, the chromosomal position on the x-axis. Identified integration site is encircled in blue.

As shown in figure 2, the vector has integrated in chromosome 3. The spike on chromosome 7 is due to the homology between the vector and the genome. Similar results were obtained with primer sets 1 and 2.

**Locus-wide coverage**

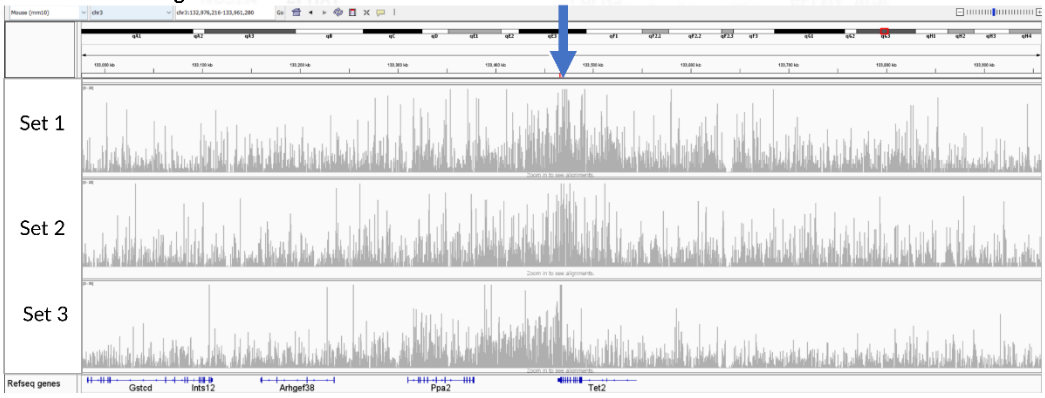

**Figure 3:** TLA sequence coverage (in grey) across the vector integration locus, mouse chr3:132,976,216-133,961,280. The blue arrow indicates the location of the breakpoint sequences. Y-axes are limited to 20x. 20x and 50x resp.

Coverage is observed across the vector integration site as shown in figure 3.

**Breakpoint sequences**

The following breakpoint sequences were identified marking the vector integration:

Vector: 194 (tail) fused to chr3: 133,465,565 (head) with 7 bp homology
GCGCTCGATTTCATCAGTTACCGTGTTTGAAGTGATCAGCGGCTTCATTACTTTTTTCAATGTGTAATCATTATTTAA
CTCTATGATACCGAGGGCGCCGTTCGCTAACTCAGCATCGCGTTTTTTAGTGCTCTGTCTGCAGCACCAAGGCTAAT
TATCACCTGCAATTAACAGACGTTCTTAAGCTGT

Vector: 90,141 (tail) fused to chr3: 133,465,642 (head) with 1 inserted base
CCTAGCACCCATGTGGCACCTCAGCCATCTGCAGCCCCAGTTCCGGGGAATCTAATGCTTTCTTCTAGCCTCTCTGG
GCACCAGACATACTTATCTACCTACCACCGTTTTACAGGATAGCAAGTAAGGTGAGCACCAAAAACTGTTATTTGTT
TTCTGTGGACTAGCATATCCATTCAGCTTCGGTTTGGATTGTAACAGGCCTGATAC

Vector: 9,387 (head) fused to chr3: 133,466,629 (head) with 3 bp homology
GtACATGCACACATCCCTGAAGATTGTTACACAAGCATAGCCAGACAAACACACACTGCATTTGTCACAGATGCTGT
GACCTGTCCCTACGGCCCATGCCATGAAAGACATACTG

Vector: 91,000 (head) fused to chr3: 133,466,728 (tail) with 1 bp homology
GATATAGAGTGTCAACTTGACATGATCTAGTAGAATCTGGGTGACAAAAACTCTGAATATATCTGTAAGGGAGCTC
CTGGATTAGGTTGACAGGCGGGTAAATATCTACCCCAAATGTGGGCATCACATTAGGCCAGACCACCAAGGACGA
CCTGTGAGCAGTATGTCTTTCATGGCATG

Chr7:141,697,592 (tail, endogenous location of Muc2) fused to chr3: 133,466,737 (head) with 2 bp homology
CCTGTGTGGACACAGATCTGGGCCCCATTCATCCCTTCCCACTTCTCTTTCATAGCATTTGCCGAAACGTCATACAA
ATGTGTTGTAAGGCCCTGTGACCTGAGTGAAAGCATATGGTGATGTAGTCACGGTAGAATCTGTAGTCACAGAC

Vector: 4,197 (tail) fused to chr3: 133,467,944 (tail) with 7 bp homology
CCGGGCACCTGTAAGGCAGGAAGCTACAACTCAGCATCCCCCACACAGCTCCTATCGAGGCTCTGATCAGGGCAG
GCGGGGCCAGCGGACATAGAGGTCACAGTTCCCGAATACCCCTCCCCCAGTCAGCACACACCTCAGATATTTATG
GAGATACCAACCATGT

Vector: 9,107 (head) fused to chr3: 133,468,175 (head) with 2 bp homology
GCCTGAGTGAAAGCTTCACTCTGTTCCTCAATTTCTCACTGAGTGAAGGCTGTTGGGTTTTAAGAAACTGCTAAGAG
GGCAGAAGTGGTGGCTCAGCGGTTAAGAGCACTTACTGTTCTTCCAAAGGACCGAAAGCTGCGGTTGTGCTATGG
AGAGGGTAGGAGAGAGCAGGTACAAACACATCTAAGAATGGAGATACTTAATATATGCAAGAGCATAGACAAGCC
CTGCAAGCAAGGACAAAGGAAA

Vector: 9,573 (head) fused to chr3: 133,468,815 (head) with 2 bp homology
CACACTTAACTCCCGGTCAGAGCTATCTCAGCTGTCAGAGCTGTTTGTTCAGTGCATTTGCTCTCCCAAGTCAAAGG
ACAAAAACACTATTGAGCCGTGATTTCCCACCAATCAAGTGTGCTGCAGAAGGATTTCTTTTTAGTAGCTCTAATTA
TAGACATACAGCAAGGGTTATTATTTATGGTGCCCAGTGCTCCTCCGTTGGGACTAAGGCAGAGTTTGATGAGTGT
GGAACATTACCTAAGCGAGAAC

In addition, the following fusions were identified indicating structural rearrangements:

Chr3: 133,466,631 (tail) fused to chr3: 133,465,615 (tail) with 3 bp homology

ACCACGGGAGAGATTAAATATAATGATAAAATAAACTATTTTGCTGGTGAGCTGACAGCCCCCCCCCAATAAACAT
ACACACAAACACAGCTTAAGAACGTCTGTTAATTGCAGGTGATAATTAGCCTTGGTGCTGCAGACAGAGCACTAA

The identified breakpoints shows that a complex genomic rearrangement has occurred in the region of the integration site.

From this data it is concluded that the vector has integrated in mouse chromosome 3: 133,465,565-133,468,815. According to the RefSeq the integrity of Tet2 may have been affected.

#### Copy number estimation

In this sample, the coverage on the vector-side is much higher than on the genome-side of the integration site (roughly 2-11 times). 1 complex integration site and 2 vector-vector junctions are found. The copy number is estimated to be 2-11 copies.

### QC information

| | |
|---|---|
| Internal project number: | 1435 |
| Lab technician: | Elaine Wong, PhD |
| Bio-informatician: | Karima Hajo, MSc |
| Analysis: | Cheryl Dambrot, PhD |
| QC-approval: | Judith Bergboer, PhD |
| Version report: | 1 |
| Date: | 02 07 2020 |

Signed by: Judith Bergboer, PhD

**Disclaimer**
Cergentis has used reasonable efforts in performing the services which are covered in this report. Cergentis does however not make any warranties, express or implied, by operation of law or otherwise, with respect to the functionality of these services. Without limiting the foregoing, Cergentis specifically disclaims all implied warranties of title, non-infringement, merchantability and fitness for a particular purpose.

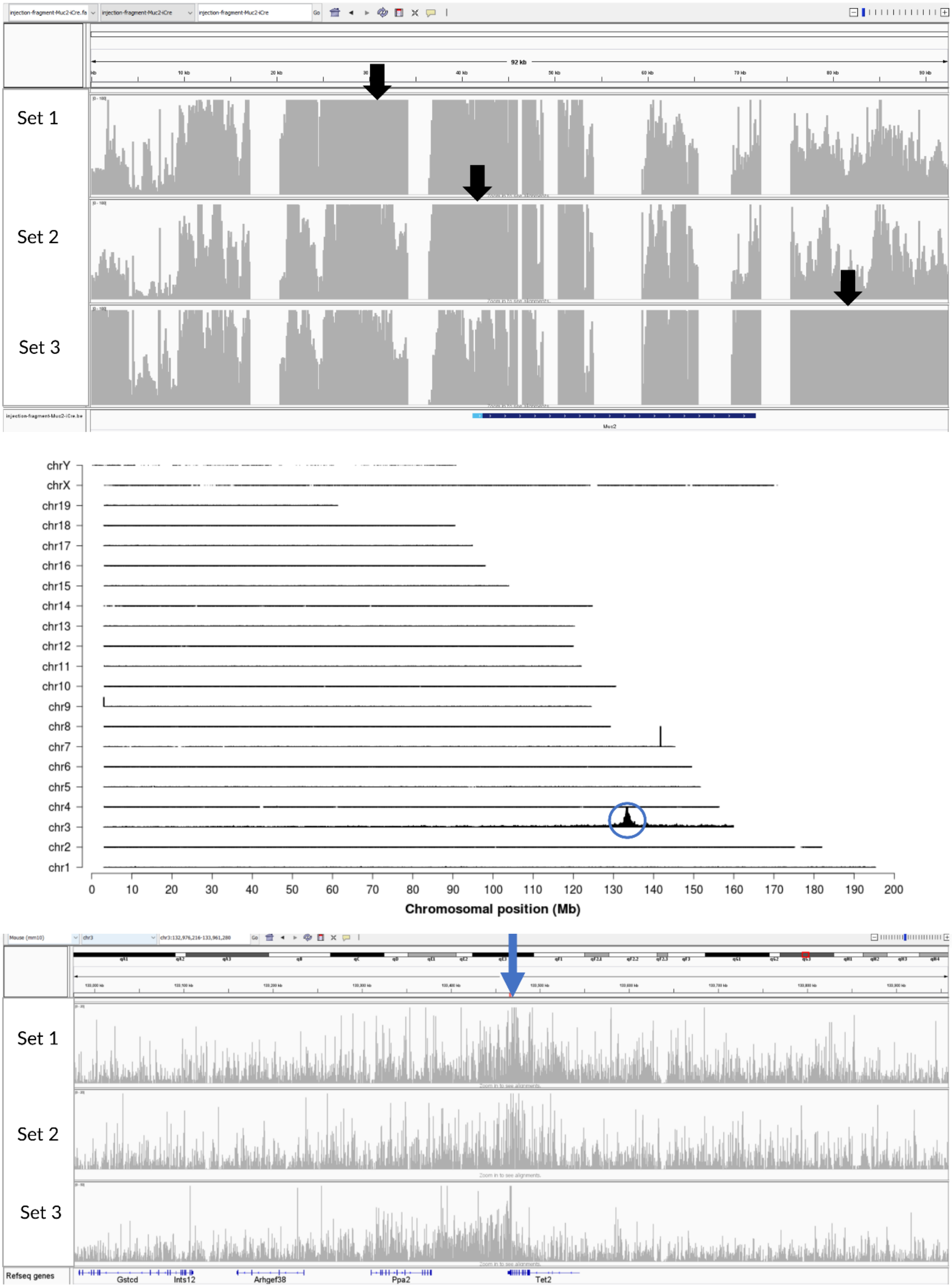
Set 1
Set 2
Set 3
chrY
chrX
chr19
chr18
chr17
chr16
chr15
chr14
chr13
chr12
chr11
chr10
chr9
chr8
chr7
chr6
chr5
chr4
chr3
chr2
chr1
0 10 20 30 40 50 60 70 80 90 100 110 120 130 140 150 160 170 180 190 200
Chromosomal position (Mb)
Set 1
Set 2
Set 3
Refseq genes
Gstcd
Ints12
Arhgef38
Ppa2
Tet2
